# ATP synthase K^+^- and H^+^-flux drive ATP synthesis and enable mitochondrial K^+^-uniporter function

**DOI:** 10.1101/355776

**Authors:** Magdalena Juhaszova, Evgeny Kobrinsky, Dmitry B. Zorov, H. Bradley Nuss, Yael Yaniv, Kenneth W. Fishbein, Rafael de Cabo, Lluis Montoliu, Sandra B. Gabelli, Miguel A. Aon, Sonia Cortassa, Steven J. Sollott

**Author notes:** Center for Scientific Review, NIH, Bethesda, MD, USA. Co-first author (Magdalena Juhaszova, Evgeny Kobrinsky, Dmitry B. Zorov). Correspondence, Lead Contact: Steven J. Sollott, M.D., Chief, Cardioprotection Section, Laboratory of Cardiovascular Science, Biomedical Research Center, Suite 100, National Institute on Aging, NIH, 251 Bayview Blvd, Baltimore, MD 21224-2816, USA.

## Abstract

ATP synthase (F_1_F_o_) synthesizes daily our body’s weight in ATP, whose production-rate can be transiently increased several-fold. Using purified mammalian F_1_F_o_-reconstituted proteoliposomes and isolated mitochondria, we show that F_1_F_o_ utilizes both H^+^- and K^+^-transport (because of >10^6^-fold K^+^ excess vs H^+^) to drive ATP synthesis with the H^+^:K^+^ permeability of ~10^6^:1. F_1_F_o_ can be upregulated by endogenous survival-related proteins (Bcl-xL, Mcl-1) and synthetic molecules (diazoxide, pinacidil) to increase its chemo-mechanical efficiency via IF_1_. Increasing K^+^- and H^+^-driven ATP synthesis enables F_1_F_o_ to operate as a primary mitochondrial K^+^-uniporter regulating energy supply-demand matching, and as the recruitable mitochondrial K_ATP_-channel that can limit ischemia-reperfusion injury. Isolated mitochondria in the presence of K^+^ can sustain ~3.5-fold higher ATP-synthesis-flux (vs K^+^ absence) driven by a 2.7:1 K^+^:H^+^ stoichiometry with unaltered OxPhos coupling. Excellent agreement between F_1_F_o_ single-molecule and intact-mitochondria experiments is consistent with K^+^-transport through ATP synthase driving a major fraction of ATP synthesis.

## Introduction

The family of ATPases shares a number of proteins with conserved functions and molecular composition (Cross and Muller, 2004). F-, A- and V-ATPases are true biological rotary engines that work as coupled motors: the F_1_/A_1_/V_1_ is chemically driven (i.e., effecting transduction of mechanical and chemical energy) and the membrane-embedded F_o_/A_o_/V_o_ is powered by the energy stored in a transmembrane ion gradient (Kuhlbrandt and Davies, 2016; Stock et al., 1999). Of these, a specialized group, the ATP *synthases*, is the major route to ATP synthesis. One of the best characterized members of ATPases is the F_1_F_o_-ATP synthase (F_1_F_o_) of E. coli, mitochondria and chloroplasts. It was demonstrated that both F_1_ and F_o_ subunits are required for ATP synthesis (Boyer, 1997).

ATP synthase operates as two rotary stepper generators coupled by a common shaft, the γ subunit (Abrahams et al., 1994; Boyer, 1997; Noji et al., 1997). The torque that is generated by ion flow through the F_o_ motor operates against the counter-torque in F_1_ driven by the energy of ATP hydrolysis. The *direction* of F_1_F_o_ is determined by which torque is larger: that of the driving force of the ion electrochemical potential or that produced by the ATP chemical potential. Under physiological conditions, F_o_ torque exceeds the F_1_-generated counter-torque at ambient ATP levels, and thus the system proceeds toward ATP synthesis. Although the principal function of the F_1_F_o_ is to harness the energy stored in electrochemical ion gradients to make ATP, it can nevertheless run backwards (as an ATP hydrolase) pumping ions in the opposite direction in the absence of the activity of a regulated inhibitory protein. This scenario would occur if, (1) the ATP levels would rise substantially relative to the ion gradient magnitude, or (2) the ion gradient becomes dissipated, as occurs during ischemia.

During ischemia, consuming substantial amounts of ATP at a time when its supply is limited would likely be detrimental in energetically-sensitive cells such as cardiomyocytes and neurons. It is known that Inhibitory Factor-1 (IF_1_), a small ~12kDa regulatory protein, limits the reversal of F_1_F_o_ function, and that during ischemia this helps to prevent excessive (or even futile) ATP consumption by damaged mitochondria to maintain ΔΨ_m_. Opening of an ATP-inhibited mitochondrial K^+^ channel (mK_ATP_), activated either by repetitive short periods of ischemia (“ischemic preconditioning”) or by K^+^ channel openers (KCO) such as diazoxide (Dz), serves as a critical link in a cascade of kinases preventing the deleterious effects of opening the mitochondrial permeability transition pore (mPTP), limiting cell damage and death after ischemia (Juhaszova et al., 2008; Juhaszova et al., 2004; Zorov et al., 2009). Interestingly, Dz binds to and enhances the inhibitory functions of IF_1_ (Contessi et al., 2004) suggesting a tendency to preserve ATP during ischemia that may lead to enhanced cell survival and resistance to damage.

Most ATPases harness the free energy of the transmembrane electrochemical proton gradient, Δμ_H_, but some use a Na^+^ gradient instead (e.g., see (Kaim and Dimroth, 1995)). While the mechanistic basis of ion-selectivity of various ATP synthases is a matter of considerable interest (Leone et al., 2015), it is even more intriguing to consider the possible significance for mitochondrial function of the accompanying “*non*-specific” ion flux via F_1_F_o_. The specificity of F_1_F_o_ for H^+^ over other cationic species was found to be extremely high (estimated >10^7^) (Feniouk et al., 2004). It can be calculated using the Goldman-Hodgkin-Katz equation (Hille, 2001) that for H^+^ selectivity values of 10^7^ and 10^8^, F_1_F_o_ would conduct a non-trivial ~24 and 2 K^+^, respectively, for every 100 H^+^ during normal ATP synthesis (at cytosolic pH=7.2 and K^+^=140 mEq/L) due to the >10^6^-fold excess of cytoplasmic K^+^ over H^+^. Given the large electrical force driving K^+^ to the mitochondrial matrix, it would make sense to harness this energy to generate ATP rather than to dissipate Δµ_K_ as heat. Because the activity of the respiratory chain is known to be regulated by intramitochondrial volume controlled by K^+^ influx (Garlid et al., 2003), the added benefit would be the direct coupling of respiratory chain activity and ΔΨ_m_ dissipation (caused by energy utilization/production) to an osmotic signal given by the amount of K^+^ traversing F_1_F_o_ to make ATP, facilitating the proportional matching between energy supply and demand. Finally, that part of the proton gradient and energy not being directly dissipated via ATP synthase because of the equivalent movement of charge as K^+^ would then be available to drive K^+^ efflux from mitochondria using the K^+^/H^+^ exchanger (KHE), thus restoring osmotic balance. These principles are fully compatible with Mitchell’s chemiosmotic mechanism (Mitchell, 1961; Nicholls and Ferguson, 2013).

We investigated the possible existence of a novel, regulated function set for ATP synthase based on the postulated ability to harness energy from K^+^ flux. This would enable K^+^ uniporter-like function and serve to facilitate energy supply-demand matching, while under certain circumstances, also to function as a mK_ATP_. We found that while retaining the high degree of H^+^-selectivity, the chemo-mechanical efficiency, and the monovalent cation conductance of F_1_F_o_ can be increased by certain KCOs and by endogenous pro-survival proteins, Bcl-xL and Mcl-1. This process requires IF_1_ and is regulated naturally by the concentration of ATP. We also demonstrated its role in protection signaling in intact cardiomyocytes.

## Results

### Potassium channel openers activate K^+^ flux into proteoliposome-reconstituted ATP synthase

First, to measure K^+^ flux into proteoliposome (PL)-reconstituted purified F_1_F_o_ (see Figure S1), the K^+^-sensitive fluorescent dye, PBFI, was trapped inside the vesicles under conditions shown in Figure 1A. In the presence of the protonophore FCCP (to enable charge balance necessary for K^+^ flux and to maintain membrane potential, ΔΨ_m_=0), the K^+^ channel opener (KCO) diazoxide (Dz) significantly enhanced the initial rate of K^+^ flux into PL; this effect was completely blocked both by the F_o_ inhibitor venturicidin B (Vent), and the mK_ATP_ blocker, 5-hydroxydecanoate (5-HD), while it was essentially absent in IF_1_-depleted F_1_F_o_ (Figure 1B).

**Figure 1.**
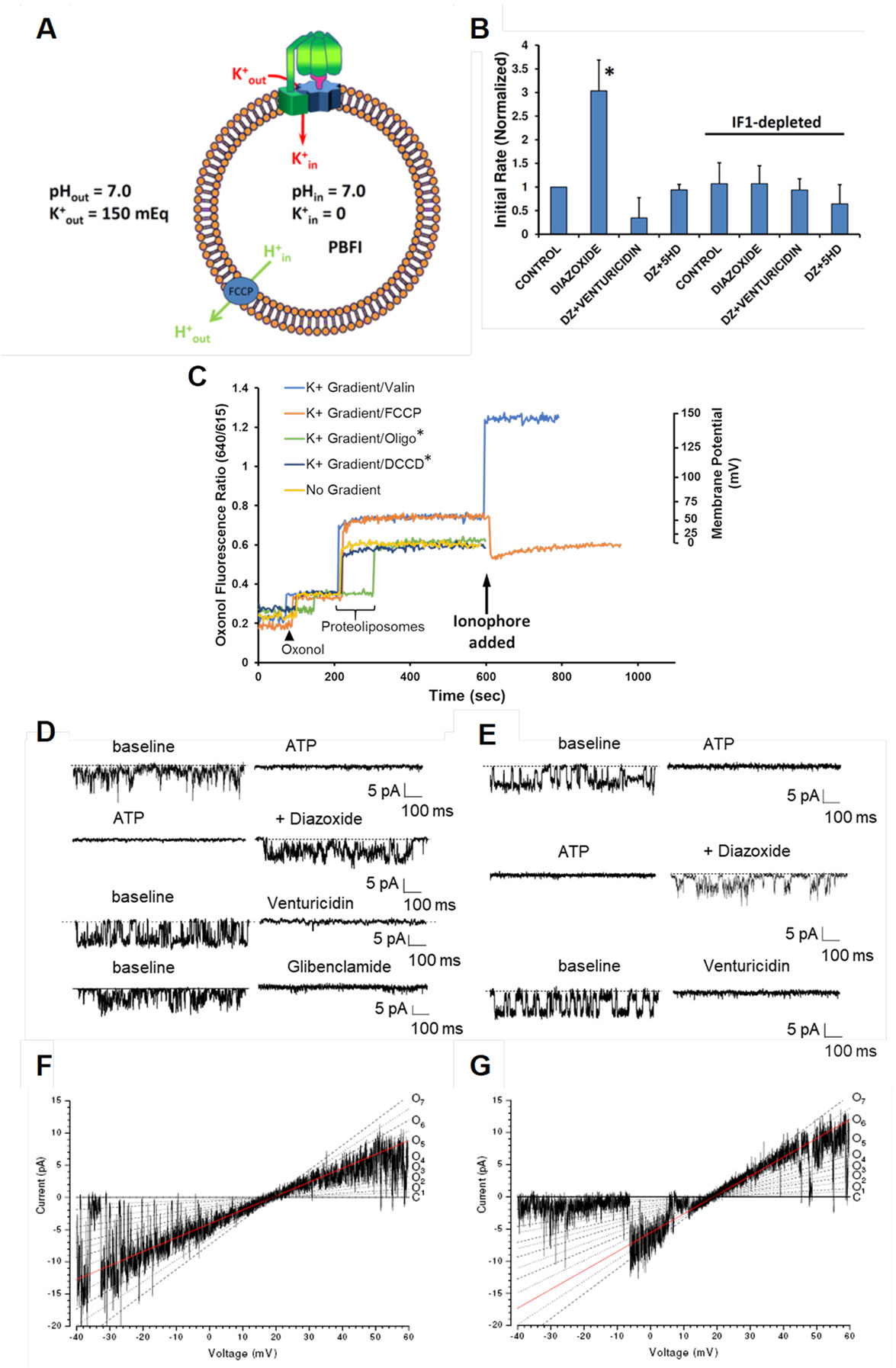
(**A**) Scheme of the PL system; ionic composition of the internal and external buffer, used in K^+^ transport experiments. (**B**) Kinetics of K^+^ flux into PL; effect of IF_1_ depletion. The KCO, Dz, significantly enhanced the rate of K^+^ flux into PL; this effect was blocked both by the F_o_ inhibitor, Vent, and the mK_ATP_ blocker, 5-HD, and was absent in IF_1_-depleted F_1_F_o_. * P<0.05. (**C**) Ratiometric fluorescence measurements of ΔΨ generated in F_1_F_o_-reconstituted proteoliposomes by a K^+^ gradient performed with oxonol VI (excitation 560, emission 640/615 nm). PL-reconstituted F_1_F_o_ develops a stable, non-zero K^+^ diffusion-potential (ΔΨ ~55 mV; orange and light blue traces) in a K^+^ concentration gradient (200 mM K^+^ _out_/0.5 mM K^+^ _in_; other conditions as in panel A except PBFI omitted) that was completely dissipated by 1 µM FCCP (orange trace). Addition of 10 nM valinomycin (instead of FCCP) resulted in a maximal K^+^ permeability and increase of the oxonol VI fluorescence ratio representing a membrane potential of ~150 mV (light blue trace). F_o_ inhibitors DCCD and Oligo (green and dark blue traces) prevented the development of ΔΨ by the K^+^ gradient; thus, ATP synthase (and not some other contaminating K^+^ conductance/channel) is responsible for ΔΨ under control conditions because it is K^+^-permeable. PL without a K^+^-gradient do not develop ΔΨ (yellow trace). The Nernst potential-calibrated values of ΔΨ were set by different K^+^ gradients using 10 nM valinomycin (right Y-axis). (**D-G**) Planar lipid bilayer experiments. (**D**) Unitary K^+^ currents from purified F_1_F_o_ and (**E**) from conventional mitochondrial membrane preparation (at −40 mV), reconstituted into lipid bilayers; pre-intervention baseline recordings are on the left, and the effect of the various compounds are shown (2 mM MgATP, 30 µM Dz, 4 µM Vent, 50 µM Glib). KCOs reverse ATP-inhibited permeation of F_1_F_o_ by K^+^ that can be blocked by Vent and Glib. (**F,G**) Unitary K^+^ currents elicited in response to a voltage ramp (14.1 mV/sec) distinguish multiple conductance levels represented by O_1_-O_7_ (C-closed state). The 216 pS conductance (O_5_) is predominantly active in the recording shown in panel **F**, while the 293 pS conductance (O_6_) is active during the recording in panel (**G**) (highlighted by red lines).

The reconstituted purified F_1_F_o_ subjected to a transmembrane K^+^ gradient at ΔpH=0 generated a stable ΔΨ_m_, reaching a maximal voltage of ~55 mV (Figure 1C). The fact that this ΔΨ_m_ remains stable until being dissipated by FCCP, in principle, rules out the possibility of contamination by an unidentified K^+^/H^+^ antiport or another charge-exchange mechanism. To further elucidate this possibility, we utilized an electrophysiological approach to rule out the presence of any other cation-selective channel activity in our F_1_F_o_ preparations (Figure S2A,B). It has been suggested that a mitochondrial ROMK potassium channel might act as the pore-forming subunit of a cytoprotective mK_ATP_ channel (Foster et al., 2012). Our immunoblotting with anti-ROMK antibody ruled out ROMK channel contamination of the isolated F_1_F_o_ (Figure S2C).

### Measurement of unitary K^+^ and H^+^ currents from F_1_F_o_

In unitary ion channel recordings from lipid-bilayer reconstitution experiments with purified F_1_F_o_, Dz reversed ATP-inhibited ion flux that can be blocked by the F_o_ inhibitor, Vent, by the mK_ATP_ inhibitor, glibenclamide (Figure 1D) and by F_o_ inhibitors oligomycin and *N,N’*-dicyclohexylcarbodiimide (DCCD; see below). Considering that similar findings are obtained in conventional mitochondrial membrane reconstitution studies (i.e., without using a purified F_1_F_o_), for the first time we show that the unitary ion currents derived from conventional mK_ATP_ preparations, which display the same characteristics as the purified F_1_F_o_ complex, can be largely inhibited by Vent (Figure 1E) at levels that do not affect sarcolemmal mK_ATP_ currents. Unitary currents from purified F_1_F_o_ exhibit multiple conductance levels (Figure 1F,G) in agreement with single channel behavior of conventional mK_ATP_ preparations. This complex behavior may arise from the multiple ion-binding positions on the F_o_ c-ring. A comprehensive literature search regarding the single channel characteristics of conventional mKATP channel preparations reconstituted into lipid bilayers indicates that multiple levels are frequently observed (five distinct peaks/conducting states between 20 pS and 120 pS in symmetric 150 mM K glutamate (Jiang et al., 2006; Nakae et al., 2003) and conductances between 100-275 pS in symmetric 100 mM KCl (Grigoriev et al., 1999; Mironova et al., 1999) which is largely consistent with the present data.

The ion-specificity of the observed unitary currents requires closer examination because mammalian F_1_F_o_ is thought to make ATP only by H^+^ flux. In the present experiments, the abundance of K^+^ over H^+^ was ~10^6^:1 (comparable to that occurring in cells), so it is reasonable to consider that K^+^ permeation may be contributing as well. Furthermore, the potential contribution of anion permeation cannot be excluded *a priori*. Because only permeant ions contribute to the net ion-current reversal potential (*E*_rev_), this was examined in detail under various conditions to assess the possibility of anion permeation and to interpret the changes in cation permeation activated by KCOs. Since complete substitution of the substantially larger and rather non-permeant Hepes anion for Cl^−^ causes no change in *E*_rev_ (17.9 ± 0.7 vs 18.6 ± 0.5 mV, respectively, *P*=ns), we concluded that the impact of Cl^−^ permeation via F_1_F_o_ is insignificant compared to cations. Additionally, the current-voltage relationship of purified F_1_F_o_ was examined in a pH gradient (pH_cis_=8.0, pH_trans_=7.2, buffered by TEA^+^-Hepes, in the absence of K^+^ or any other small cation aside from H^+^) yielding a reversal potential identical to the expected Nernst potential for H^+^ as the only permeant species, in agreement with the idea that the OH^−^ anion is non-permeant. Since the measured unitary ion currents under control conditions consist only of H^+^ and K^+^, we next assessed their relative contributions. Under ionic conditions where the reversal potential for H^+^ (*E*_H_) was 0 mV and that for K^+^ (*E*_K_) was +28 mV, *E*_rev_ was found to be ~+18 mV indicating that both cations must be permeant and contributing to the total currents observed. In this case, *E*_rev_ is given by:

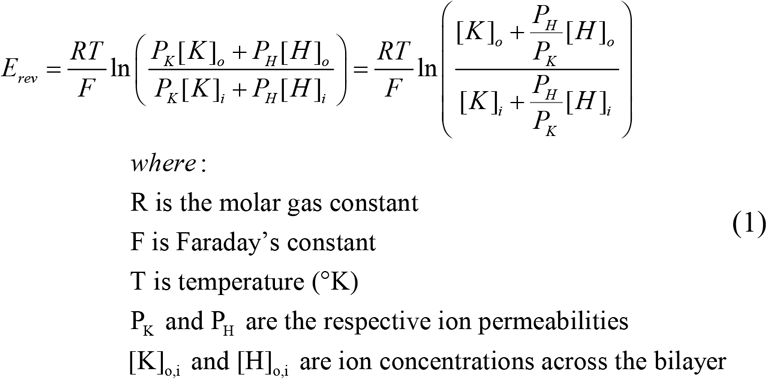

Maintaining steady voltages at each of the ion-reversal potentials (to remove the driving force from the specific cation, thus rendering pure unitary current from the other cation) produced macroscopic H^+^ and K^+^ currents (at +28 and 0 mV, respectively) of the same characteristics and open probability (P_o_) as at − 20 or −40 mV (Figure 1F,G). Importantly, the KCO Dz increases P_o_ and amplitude for both H^+^ and K^+^ currents (Figure 1D,E) without causing any change in *E*_rev_ (18.3 ± 1.3 vs 18.5 ± 0.9 mV with Dz, *P*=ns; similar data was obtained with Na^+^: 18.1 ± 1.0 vs 18.9 ± 1.0 mV; n=10). This indicates that while the permeability for H^+^ and K^+^ after Dz increases for both cations, the ratio (*P*_H_/*P*_K_) remains unchanged (see eq.1). From the Goldman-Hodgkin-Katz (GHK) formalism, the total ion current is related to the individual ion permeability as follows:

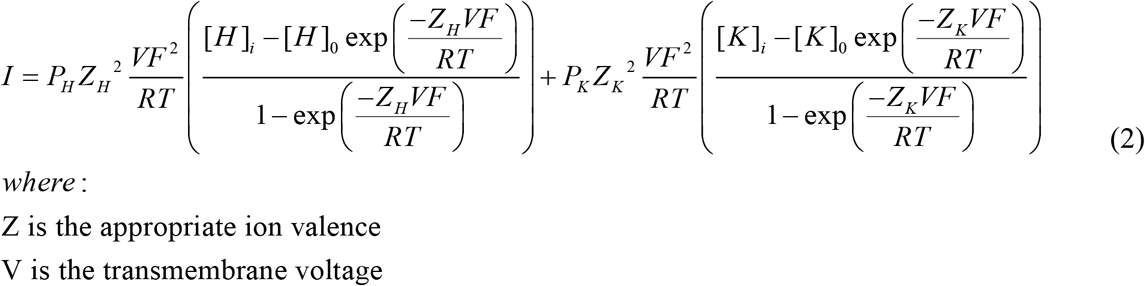

We determined that the baseline values for *P*_H_ and *P*_K_ (5.2±0.9 ×10^−11^ and 8.7±2.9 ×10^−17^ m^3^/s, respectively) each increases ~3.5-fold after Dz (to 2.2±1.3 ×10^−10^ and 3.0±1.4 ×10^−16^ m^3^/s, respectively, n=4, both *P*<0.05 *vs*. each ion’s baseline value), thus keeping the selectivity of F_1_F_o_ for H^+^ over K^+^ at ~10^6^:1. Regarding the single ion channel behavior of the conventionally-prepared mK_ATP_, close inspection of the electrophysiological data referred in eight published papers allowed us to extract baseline values for *P*_H_ and *P*_K_ (mean values of 4.9±1.1 ×10^−11^ and 1.9±0.5 ×10^−16^ m^3^/s, respectively) (Bednarczyk et al., 2005; Dahlem et al., 2004; Grigoriev et al., 1999; Jiang et al., 2006; Mironova et al., 1981; Mironova et al., 2004; Nakae et al., 2003; Zhang et al., 2001) which compare extremely well with those obtained here for F_1_F_o_.

The importance of F_1_F_o_ as a major K^+^ pathway can be realized from Eq. 2 which shows the sum of the K^+^ and H^+^ current components in the GHK formulation. For sufficiently large ΔΨ_m_ magnitudes (>100 mV for the present case), a rearrangement of Eq. 2 expressing the ratio of K^+^ to H^+^ conducted by F_1_F_o_ simplifies to the limiting value, [(*P*_*K*_·[K^+^]_*cytosol*_)/(*P_H_*·[H^+^]_*cytosol*_)] at *negative* ΔΨ_m_ (the direction of ATP *synthesis*). This means that F_1_F_o_ may potentially conduct an average of 3.7 K^+^ for every H^+^ during *normal* ATP synthesis (at cytosolic pH=7.2 and K^+^=140 mEq/L; (see section: **Quantitative comparison of H^+^ and K^+^ current magnitudes through ATP synthase in** Supplemental information).

### ATP synthesis by proteoliposome-reconstituted F_1_F_o_ in a K^+^ gradient

To investigate whether ATP synthase can harness energy from K^+^-flux, purified F_1_F_o_ reconstituted into PL was subjected to a transmembrane K^+^ gradient (Figure 2A). Under conditions in which ΔΨ=0, and Δμ_H_ and H^+^ were unable to drive ATP synthesis due to the presence of protonophore, FCCP, we show that ATP synthesis driven by the K^+^ gradient occurs under these conditions, and was increased several-fold by the KCOs, Dz or pinacidil. This ATP synthesis was inhibited by the specific F_o_ inhibitor, Vent, the mK_ATP_ blocker, 5-HD, and abolished by the K^+^ ionophore, valinomycin (Val) Figure 2B-D). Interestingly, the F_1_F_o_ is not selective among alkali ions after KCO-activation, since we observed a comparable degree of ATP synthesis in a Na^+^ gradient, akin to that in a similar K^+^ gradient (Figure 2E). Moreover, this effect seems to be restricted to small cations, since there is no ATP generated by KCO activation of the F_1_F_o_ in a comparable gradient of TEA^+^Cl^−^. Similar K^+^ gradients with either Cl^−^ or SO_4_^2−^ as counterions yielded comparable ATP amounts (not shown). It is important to note that FCCP is necessary to enable ATP synthesis driven by the K^+^ gradient because it equilibrates the transmembrane charge (via passive inward diffusion of H^+^) owing to Δµ_K_-driven K^+^ efflux via F_1_F_o_. Omitting FCCP eliminates ATP synthesis ruling out the possible contamination of the PL’s by an unsuspected protein which might cause an underlying K^+^/H^+^ antiport activity (see Figure S2 for additional details). Thus, any possibility that the observed ATP synthesis was due to F_1_F_o_ surreptitiously harnessing H^+^ flux energy (somehow derived from the original K^+^ gradient energy) is ruled out by this FCCP result, because ATP synthesis would instead have been prevented by the protonophore-elicited dissipation of H^+^ energy. This evidence also shows that K^+^ can drive ATP synthesis by the same mechanism and path as H^+^.

**Figure 2.**
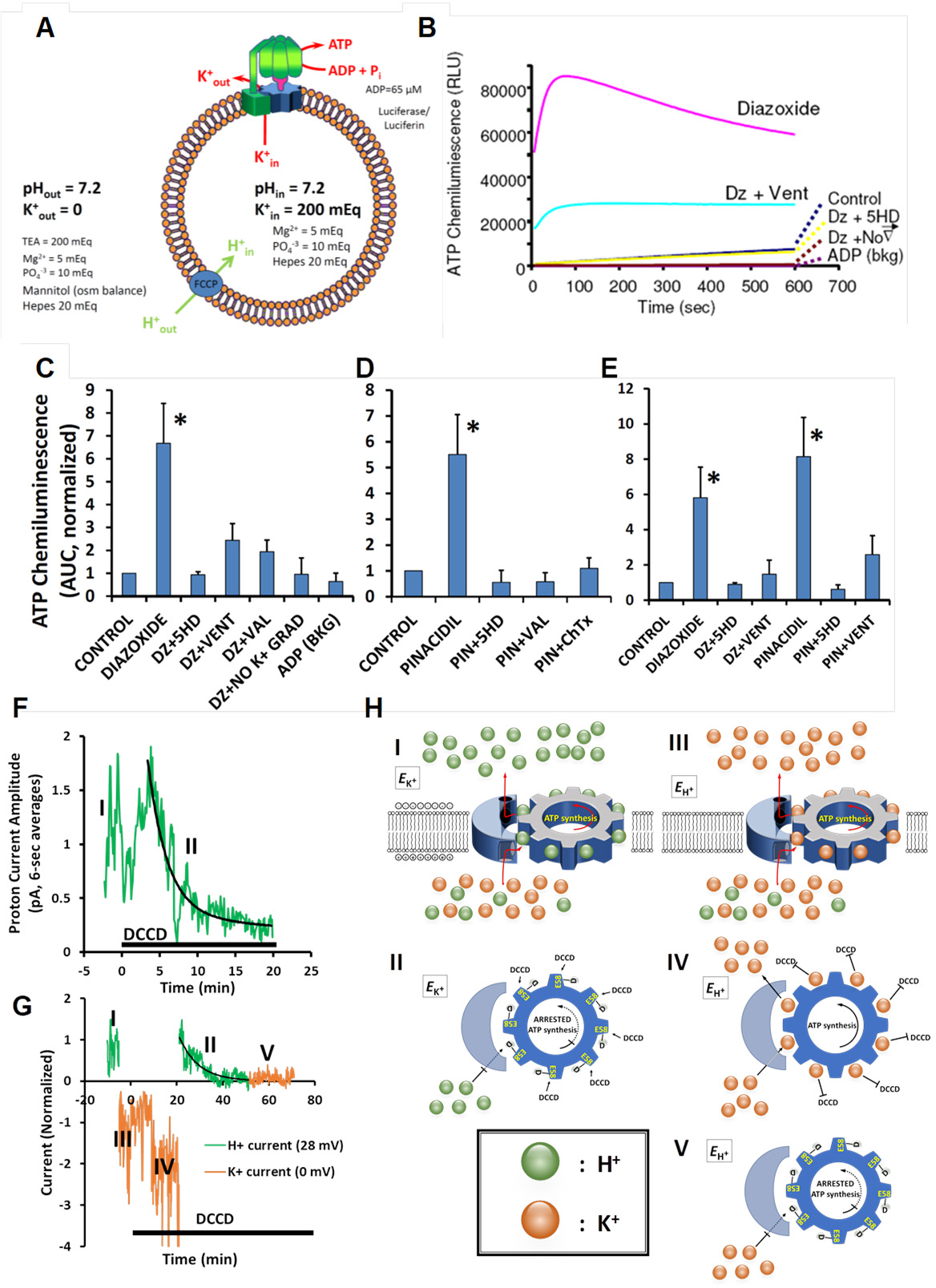
(**A**) Scheme of the PL and reaction conditions to measure K^+^ transport-coupled ATP synthesis (and hydrolysis). Δμ_H_=0 for ATP synthesis and ΔΨ=0 by using FCCP; n.b., omitting FCCP eliminates ATP synthesis ruling out the presence of an unsuspected, underlying K^+^/H^+^ antiport mechanism. (**B,C**) K^+^-gradient-driven ATP production in PL is activated by Dz and (**D**) pinacidil, and (**B-D**) attenuated by inhibitors of F_1_F_o_, inhibitors of the mK_ATP_ and BK_Ca_ channels, and by K^+^ gradient dissipation (Val or No K^+^ grad). (**E**) Na^+^-gradient-driven ATP production in PL is activated by Dz and pinacidil and attenuated by inhibitors of complex V and the mK_ATP_. These experiments prove that the entity activated by KCOs is the F_1_F_o_. * P<0.05. (**F-H**) Effect of DCCD on lipid bilayer-reconstituted F_1_F_0_ activity during conduction of pure H^+^ (at +28 mV, green traces) and K^+^ (at 0 mV, orange traces) currents (*cis*: 150 mEq K^+^, pH 7.2/ *trans*: 50 mEq K^+^, pH 7.2; 1 kHz-recorded data for multiple-minute experiments are plotted as 6-sec averages). (**F**) Continuous recording of unitary H^+^ current (phase ‘I’; green trace); H^+^ current is completely blocked by DCCD with t_½_~6 min (phase ‘II’; exposure to DCCD is indicated by the black horizontal bar). (**G**) Baseline performance of F_1_F_o_ initially driven by H^+^ current (phase I; green), then switched to K^+^ current (phase ‘III’, orange), followed by addition of DCCD (black bar, initiated at “0” time) which fails to block K^+^ current-driven activity after 20 min (phase ‘IV’, orange), then switched back to H^+^ current (at 20 min DCCD exposure, phase ‘II’, green) leading to rapid inhibition of activity, then back to K^+^ current (at 50 min DCCD, phase ‘V’, orange) displaying persistence of inhibition. (**H**, Panels **I-V**) Mechanistic schema of ‘phases’ of F_1_F_o_ driven by K^+^ (orange) or H^+^ (green) currents corresponding to those depicted in panels F,G. (**H_I_**) H^+^-driven rotation at the reversal potential for K^+^ (*E*_K+_); (**H_II_**) inhibition of H^+^-driven rotation after DCCD reaction with protonated E58; (**H_III_**) K^+^-driven rotation at the reversal potential for H^+^ (*E*_H+_); (**H_IV_**) inability of DCCD to inhibit K^+^-driven rotation due to occupancy of E58 by K^+^ (rather than by a H^+^ required for DCCD reaction and formation of the acylurea adduct); (**H_V_**) once the enzyme is deactivated by DCCD during prior H^+^ current passage (and formation of a stable acylurea adduct on E58), subsequent K^+^ flux is also blocked. Together, these results serve as another line of proof that mammalian F_1_F_o_ can operate utilizing K^+^ flux, and that both K^+^ and H^+^ travel the same route within the complex on the *c*-ring.

To further investigate whether the K^+^ and H^+^ paths through F_1_F_o_ are the same, we utilized the inhibitor DCCD that acts by specifically and covalently modifying the *protonated* form of a conserved glutamate carboxylate (E58) located in the binding pocket of the membrane-embedded c-ring (Kluge and Dimroth, 1993; Pogoryelov et al., 2010; Symersky et al., 2012) and performed measurement of unitary ion channel currents from F_1_F_o_ under conditions where we could generate and select pure K^+^ and H^+^ currents by setting specific membrane potentials. Since E58 is critical for coordinating ion binding necessary for the H^+^ gradient to generate ATP, we found that DCCD rapidly inhibits ion flux (t_½_ ~6 min) when being driven by pure H^+^ current (Figures 2F,H_I,II_), as expected. So long as the *c*-ring binding pockets are H^+^-bound, continuing active H^+^ flux is not required for DCCD’s block since we found that the inhibition fully develops when the *c*-ring is “stalled” without ion flow after running pure H^+^ current (Figure S3).

We hypothesized that E58 in the *deprotonated* state is also critical for K^+^ binding related to its generation of ATP, and that occupancy of the same binding site by K^+^ (likely by preventing glutamate protonation via charge neutralization and physical exclusion) would specifically prevent F_1_F_o_ from reacting with, and being inactivated by, DCCD (Figures 2G,H_III,IV_). We employed K^+^-only current conditions (i.e., under the present experimental conditions at 0 mV where there is no H^+^ driving force, and hence, no H^+^ current) for 5 min *before* exposure to DCCD (Figure 2G orange trace labelled ‘III’, and Figure 2H_III_). We found that the maintenance of K^+^-flux and binding *prevented* the subsequent DCCD block for at least 15 min (Figure 2G, orange trace labelled ‘IV’, and Figure 2H_IV_), and for as long as 1 hour under continuous K^+^ currents (Figure S3), indicating protection of E58 by K^+^ from the inhibitory reaction with DCCD. Similar protection by Na^+^ from inhibition by DCCD was described for this conserved, essential Glu site of the *c*-subunit from *P. modestum* (Kluge and Dimroth, 1993). Returning to driving F_1_F_o_ with H^+^ currents after the “protective” K^+^ current protocol, in the continuing presence of DCCD, lead to rapid and irreversible inhibition of the enzyme (Figure 2G green trace labelled ‘II”, Figure 2H_II_, and Figure S3). Once the enzyme was deactivated by DCCD during H^+^ current passage, we found that K^+^ flux is also blocked (Figure 2G orange trace labelled “V” and Figure 2H_V_; Figure S3), consistent with the DCCD reaction forming a stable acylurea adduct on E58. Together, these results serve as another line of proof that mammalian F_1_F_o_ can operate utilizing K^+^ flux, and that both K^+^ and H^+^ travel the same route within the complex on the *c*-ring.

### Direct demonstration of K^+^-driven ATP synthesis by simultaneous measurements of K^+^ currents and single-photon detection of ATP: “Single Molecule Bioenergetics”

To directly demonstrate that ATP synthesis can be driven specifically by K^+^ flux through F_1_F_o_ and the energy stored in an electrochemical K^+^ gradient, we designed a unique system for simultaneous measurement of unitary K^+^ ion currents (by voltage clamp) together with ATP synthesis (from low light-level detection of bioluminescence) generated by single molecules of ATP synthase (Figure 3A-F). Single molecules of purified F_1_F_o_ were reconstituted in the lipid bilayer formed on a 30-60 µm glass pipette tip (Figure 3A). Pure K^+^ current-driven ATP synthase activity due to Δµ_K_ produced by the experimental K^+^ gradient (at 0 mV, when Δµ_H_=0) was accompanied by a significant increase in the photon rate (i.e., real ATP generation) over background (at 18 mV with F_1_F_o_ stalled and no K^+^ or H^+^ currents) that in turn was significantly inhibited by specific F_o_ blockers, Vent/Oligo (representative experiment in Figure 3B; concatenated results in Figure 3C). *This experiment provides unambiguous and definitive proof of K^+^-driven ATP production by single molecules of mammalian ATP synthase under conditions reasonably matching the physiological K^+^ ionic milieu*. In this experiment, K^+^, the only monovalent cation capable of performing electrochemical work, is thermodynamically driven by its preset gradient in the direction of ATP synthesis, whereas protons are incapable of performing work because the experiment is performed under circumstances constraining Δµ_H_=0. That the entity that synthesizes ATP as a result of the work performed by K^+^ flux is significantly blocked by the specific F_o_ inhibitors, Vent and Oligo, provides corroborating proof that ATP synthase is responsible for the observed synthesis of ATP, and not some unknown (or undescribed) complex.

**Figure 3.**
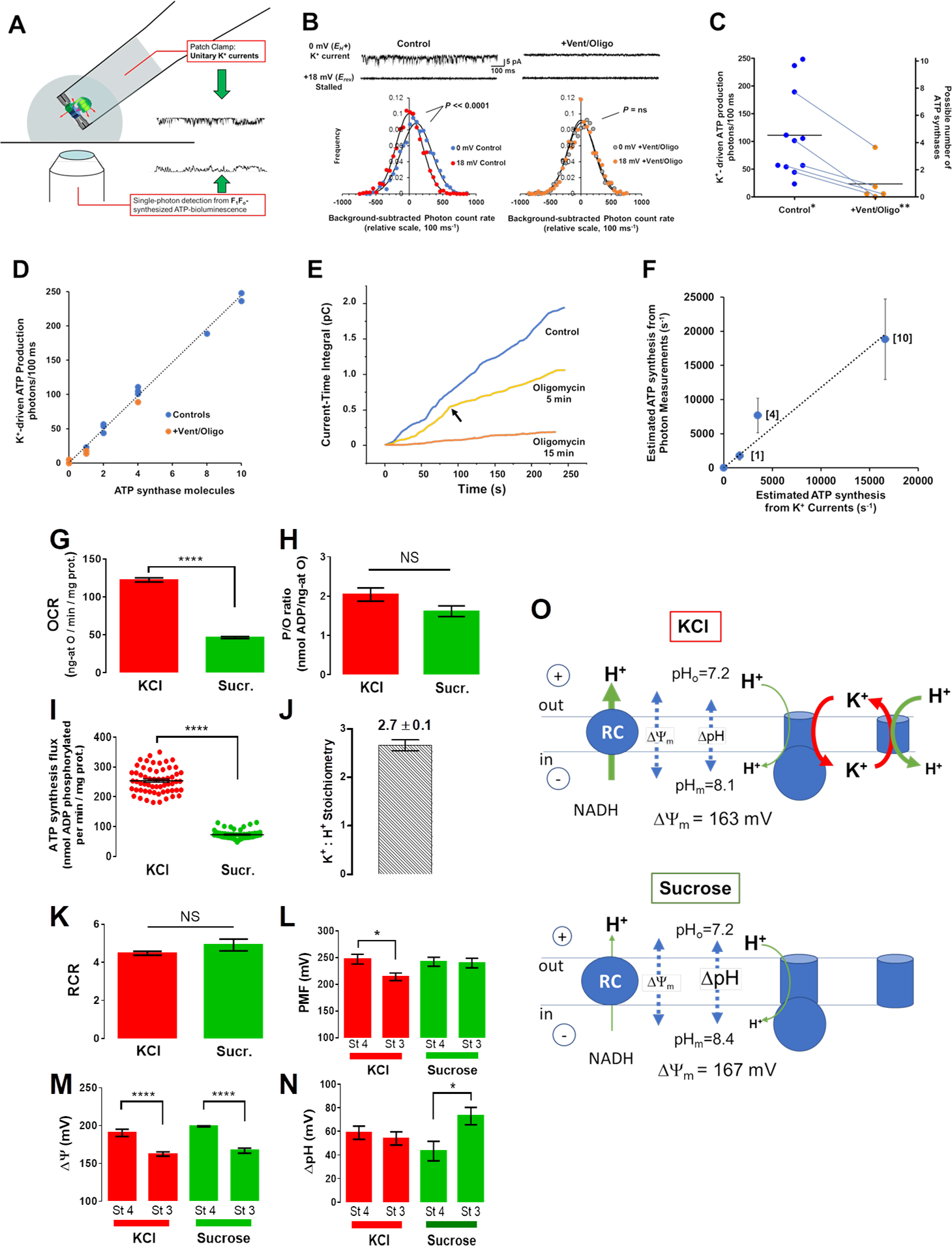
(**A-F**) Direct, “single molecule bioenergetics” demonstration of K^+^-driven ATP synthesis by F_1_F_o_ using simultaneous measurements of K^+^ currents and single-photon detection of ATP. (**A**) Experimental scheme for simultaneous measurements of unitary K^+^ currents (by voltage clamp) and low light level detection of ATP synthesis activity (luciferase/luciferin bioluminescence) from reconstituted single molecules of F_1_F_o_ in a lipid bilayer formed on a 30-50 µm glass pipette (*cis*: 150 mEq K^+^, pH 7.2/ *trans*: 50 mEq K^+^, pH 7.2). (**B**) Representative experiment performed as described in (A). Top traces: K^+^ current in control (left) and that after F_1_F_o_ inhibition by Vent+Oligo (right), together with corresponding measurements with F_1_F_o_ activity stalled at *E*_rev_. Bottom panels: Frequency histograms of detected bioluminescence photons from ATP generated by K^+^-driven F_1_F_O_ (background-subtracted photon counts measured during K^+^ current (0mV) and at the E_rev_ (18mV); Control experiment (left) vs Vent+Oligo block (right)). Note that the Control photon-rate frequency histogram during active K^+^ currents (at 0 mV, *E*_H+_) shows the production of ~50 photons/100 ms above that when stalled (no currents at +18 mV, *E*_rev_; n=1000 observations each histogram, *P*<<0.0001 for paired comparison), and that both the 0 mV current and associated photon production are abolished with +Vent/Oligo (*E*_H+_ vs *E*_rev_; n=1000 observations each histogram, *P*=ns). (**C**) Concatenated experimental results: K^+^-driven photon production in control (0 mV vs stalled at +18mV, n=11 paired observations, \**P*<<0.0001 by Fisher’s combined probability test) and that after F_1_F_o_ inhibition by Vent+Oligo (paired observations with control, n=5, \*\**P*<<0.0001 by Fisher’s combined probability test); measured photon production rates from K^+^-driven ATP synthesis appeared to be quantized in relation to the number of incorporated F_1_F_o_ (“Possible number of ATP synthases” on right y-axis; see text for further details). (**D**) Analysis of apparent quantal nature of K^+^-driven photon (ATP) production rate vs the proposed number of bilayer-incorporated ATP synthase molecules (from experimental results, panel (**C**); regression analysis slope of ~25 photons/100 ms/ATP synthase, R^2^=0.99. (**E**) Evidence for F_1_F_o_ lipid bilayer insertions as functional dimers. Time course of the cumulative K^+^ current-time integral recording (running ATP synthesis index) from a representative F_1_F_o_ experiment in control (blue trace) and during Oligo inhibition (yellow and orange traces); arrow (yellow trace, 5 min Oligo) indicates abrupt change of the current-time integral slope (to half of the control value) consistent with the initial activity of an F_1_F_o_ dimer being reduced by one functional unit to the equivalent of a monomer; after 15 min Oligo (orange trace), the activity of the remaining F_1_F_o_ (monomer) becomes fully inhibited (n.b., slight deviation from a zero-slope is a typical baseline artifact of cumulative electrical drifts in 4+ minute electrical recordings, and because it occurs in lipid bilayer recordings without protein incorporation, does not represent residual ATP synthase activity). (**F**) Relationship between ATP synthesis rates calculated from single photon vs K^+^ current measurements (see text for details). The number of F_1_F_o_ molecules is shown in brackets for several independent experiments; the average ATP synthesis rate is ~1000 ATP/sec/F_1_F_o_ driven solely by K^+^ current in the nominal absence of ATP (thus without significant counter-torque on F_1_, in contrast to physiological conditions). ATP synthesis rates estimated independently from single photon vs K^+^ current measurements are in excellent agreement, deviating from unity by <18%, R^2^=0.98. (**G**) Seahorse-instrument oxygen consumption rate (OCR) of isolated rat heart mitochondria under state 3 was determined in KCl- or sucrose-based assay medium in the presence of 5/5mM G/M and 0.5mM ADP. The oligomycin-sensitive OCR was obtained by subtracting from state 3 OCR the state 4 OCR measured in the presence of 10μM oligomycin; n=60/3 experiments, *P*<0.0001. (**H**) P/O ratio was measured using high resolution respirometry as described in Methods under similar conditions as those described in panel G; n=9/3 experiments, *P*=ns. (**I**) The ATP synthesis flux was calculated as the product of the oligomycin-sensitive OCR times the P/O ratio in K^+^- or sucrose-based assay medium; n=60/3 experiments, *P*<0.0001. (**J**) The K^+^/H^+^ stoichiometry was calculated from the difference between the ATP synthesis flux in the presence of KCl *vs* in the presence of sucrose, in ratio to the ATP flux in sucrose; n=46/3 experiments. (**K**) Respiratory control ratio (RCR) was obtained from the OCR ratio state 3/state 4; n=60/3 experiments, *P*=ns. (**L**) The proton motive force (PMF) was determined according to: PMF = ΔΨ_m_ - Z·ΔpH using ΔΨ_m_ (panel (**M)**; n=8/3 experiments; *P*<0.0001) and Z·ΔpH (panel (**N**); n=8/3 experiments; *P*<0.05) obtained under the indicated conditions with Z=2.303·RT/F (see also Supplemental Experimental Procedures). (**O**) Summary scheme describing the ion fluxes and their respective driving forces, ΔΨ_m_ and ΔpH, determined under state 3 respiration in K^+^- or sucrose-based medium (see also Supplemental Experimental Procedures). \**P*<0.05; \*\*\*\**P*<0.0001

The range of measured photon production rates from these experiments varied by an order of magnitude (~25 to 250 photons/100 ms) and appeared to be quantized (allowing for “noise”) with a “greatest common factor” of ~25 photons/100 ms which is probably consistent with the activity of 1 monomer of ATP synthase (Figures 3C right *y*-axis, D). We expected that there could be occurrences of multiple ATP synthases since these latter experiments were prepared with an increased amount of purified F_1_F_o_ intended to increase both the potential for multiple insertions of ATP synthase into the lipid bilayer and the detected bioluminescence signal over noise. Plotting the K^+^-driven photon production rate vs the proposed number of ATP synthase molecules (from 1 to 10 ATP synthases), and performing regression analysis of all experimental results, rendered a slope of ~25 photons/100 ms/ATP synthase (Figure 3D). The data pattern suggests that most of the insertions observed are of ATP synthase *dimers* and multiples thereof, whereas isolated monomers appeared to be relatively rare (accounting for <10% of observed occurrences). There are several additional lines of evidence for the idea of ATP synthase dimer-insertions: (1) examining the time course of inhibition of K^+^ current flux by oligomycin to find a quantal pattern of inhibition. In experiments producing ~50 photons/100 ms at baseline (indicated as dimers in Figure 3C) the current-time integral (CTI, the precise “electrical analog” of cumulative ATP synthesis) shows in a representative example that at ~5 min of oligomycin exposure the time-dependent slope of CTI (proportional to ATP synthesis *rate*) is abruptly reduced by exactly half, consistent with the activity of a dimer being reduced by one functional unit to the equivalent of a monomer (Figure 3E, yellow trace, arrow), and subsequently, after 15 min, the activity of the remaining ATP synthase becomes fully inhibited (Figure 3E, orange trace). (2) Purified ATP synthase is isolated mainly as dimers in the current protocol, as gauged by the abundance of ATPase activity in Clear Native PAGE in-gel assay (Figure S1). (3) ATP synthases form rows of natural dimers along cristae in mitochondria (e.g., as shown in five different species: bovine, *Y. lipolytica*, *P. anserina*, *S. cerevisiae*, and potato (Davies et al., 2011) and are isolated as natural dimers by other workers (e.g., as shown using transmission electron microscopy (Couoh-Cardel et al., 2010).

We sought to further quantify and compare the ATP synthesis rates independently derived from single molecule electrical and bioluminescence measurements. Bioluminescence-emitted photon measurements, concatenated with the efficiencies of the ATP-luciferase reaction, light path and camera detection characteristics, together with the fraction of the solid angle collected from an isotropically-emitting point source by the objective lens, produce a quantitative estimate of ATP production rate (see Supplementary Methods). Additionally, an independent estimate of ATP synthesis rate from these same experiments can be derived from the parallel unitary current measurements since 3 ATP’s are made from the flux of 8 ions through mammalian ATP synthase. Excellent agreement was obtained for single molecule ATP synthesis rates calculated from photon vs K^+^ current measurements incorporating the range of 1, 4 and 10 ATP synthases (<13% deviation from the line of identity; Figure 3F, r^2^=0.93), yielding the consistent value of ~1,900 ATP/s/ATP synthase driven by K^+^ (±50, SEM of n=11), which is in good agreement with published data (e.g., (Watt et al., 2010)). Note that ATP synthase is operating in these experiments against a negligible counter-torque because the ambient [ATP] is negligible, which is experimentally necessary to resolve the extremely low levels of bioluminescence generated from the driven activity of single molecules of ATP synthase. In the presence of “physiological” levels of ATP (~3-4 mM) the resultant chemical counter-torque exerted at F_1_ against the electrical torque driving F_o_ would reduce ion flux (conductance) and correspondingly ATP synthesis by about an order of magnitude (see section on “Regulation of mechano-chemical efficiency of F_1_F_o_” below, and Figure 5H), so the equivalent ATP synthesis rate in these K^+^-driven experiments would be ~190 ATP/s/ATP synthase (±5, SEM of n=11) in an intracellular milieu, which is in good agreement with published data (Diez et al., 2004; Etzold et al., 1997; Junesch and Graber, 1991; Soga et al., 2012).

### K^+^-driven ATP synthesis in isolated heart mitochondria

Measurements of K^+^ flux, a stable K^+^-diffusion potential, and purely K^+^-driven ATP synthesis in proteoliposome- and single molecule-reconstituted ATP synthase (Figures 1, 2, 3A-F), together with unitary K^+^ and H^+^ currents sustained by purified F_1_F_o_ reconstituted in lipid-bilayers (Figure 1D, F, G), show that ATP synthesis can happen under physiological conditions (cytosolic pH=7.2 and K^+^=140 mEq) in which F_1_F_o_ is driven by up to ~ 3.5 K^+^ for every H^+^. These findings predict that, in mitochondria, K^+^ transport carried by F_1_F_o_ would share a sizable portion of the ATP synthesis flux driven by Δµ_H_ with a high K^+^:H^+^ stoichiometry.

To assess these predictions, we studied the effect of physiological levels of K^+^ on the ATP synthesis flux and the corresponding K^+^:H^+^ stoichiometry in isolated mitochondria at constant osmolality (260 mOsm) in the presence vs absence of K^+^ (isosmotically substituting sucrose for K^+^). We quantified the Oligo-sensitive oxygen consumption rate (OCR) with high-throughput Seahorse respirometry (Figure 3G), and ADP/O ratio with high resolution respirometry (Oroboros) (Figure 3H). At similar respiratory control ratio (RCR~5) (Figure 3K), ATP synthesis was 3.5-fold higher in the presence of K^+^ than in its absence (Figure 3I) with a K^+^:H^+^ stoichiometry of 2.7:1 (Figure 3J). More robust ATP synthesis with K^+^ resulted from a 2.65-fold higher respiratory flux (Figure 3G) at ADP/O ratio 2.0 and 1.6 (Figure 3H) in the presence and absence of K^+^, respectively. Under similar conditions, a significant mitochondrial K^+^ influx happens during state 4-to-3 transitions (Figure S4B). The OCR values were within the range of reported data under similar conditions (Table S1).

Next, we employed radioactive tracers to measure the protonmotive force (PMF), and its individual components, ΔΨ_m_ and ΔpH, in isolated rat heart mitochondria in the presence or absence of K^+^ at constant (260mOsm) osmolality under states 4 and 3 respiration (Figures 3L-N). While ΔΨ_m_ was similar under both respiratory states and in absence or presence of K^+^ (Figure 3M), ΔpH was significantly higher under state 3 compared to state 4 respiration in the absence of K^+^ whereas it remained constant in its presence (Figure 3N). This resulted in 0.3 pH units increase in the matrix when K^+^ was absent (pHi 8.4 *vs*. 8.1) (Figure 3O) with pH 7.2 in the medium. Unlike in K^+^ absence, where both ATP synthase and the respiratory chain (RC) are operating at slower rates with flux through the RC higher than through ATP synthase resulting in matrix alkalinization driven by similar ΔΨ_m_, in K^+^ presence, the matrix pH elevation does not happen due to the additional flux of K^+^ through F1Fo accompanied by enhanced ATP synthesis and H^+^ entering mitochondria through the K^+^/H^+^ exchanger (Fig. 3). Importantly, PMF, as an energy-proportional index, exhibited a significant decrease under state 3 respiration in the presence of K^+^, consistent with the additional output of ATP, whereas it remained constant in its absence (Figure 3L).

Using dynamic, simultaneous, fluorometric monitoring of ΔΨ_m_, NAD(P)H and volume (90° light scattering) rather than steady state measurements, we investigated possible effects of matrix alkalinization on carbon substrate oxidation during state 4→3 transition in mitochondrial suspensions subjected to the addition of ADP pulses of increasing concentration (Fig. S4H-M). Under both conditions of K^+^ presence and absence, the RC flux is controlled downstream of NADH, e.g., by the RC activities, ATP synthase, etc., (i.e., the “pull” condition) (Cortassa et al., 2003; Wei et al., 2011) as can be judged by the *in-phase* response of mitochondrial ΔΨ_m_, NAD(P)H and volume, in response to the increase in energy demand exerted by the ADP pulses, ruling out a matrix alkaline-pH limitation on substrate oxidation in either condition (Fig. S4H-D).

Mitochondrial volume changes associated with the state 3→4 transition, as measured with radioactive tracers, were greater in the K^+^-containing medium (1.25±0.15 to 0.81±0.16) than in sucrose (1.27±0.19 to 1.06±0.13 µl/mg prot) reflecting a more dynamic and higher rate of ADP influx and conversion to ATP, thus higher amplitude of volume changes in salt medium associated with the transition to fully energized state (see also Figure S4L-O). These results were corroborated by our fluorometric simultaneous measurements of ΔΨ_m_ and volume of isolated heart mitochondria, where we show that, in the presence of K^+^, mitochondria exhibit significantly faster ATP synthesis as revealed by the shorter time lapse utilized to fully consume the ADP added (roughly 1000 sec *vs*. 1600 sec: compare H and I in Figure S4; see also panels N,O).

Together, our data indicate that mitochondria synthesize 3.5-fold higher rates of ATP synthesis in the presence of K^+^ compared to osmotically-matched conditions in which this cation is absent. The 2.6-fold higher respiratory flux is mainly responsible for the higher ATP synthesis flux driven by a 2.7:1 K^+^:H^+^ stoichiometry. Thus, in the presence of K^+^, both Δµ_K_ and Δµ_H_ energies (Δµ_H_ producing Δµ_K_ almost entirely through the driving force of ΔΨ_m_) are utilized to synthesize ATP with the resulting matrix-accumulated K^+^ being continuously restored via the Δµ_H_-driven K^+^/H^+^ exchanger (Figure 3O, top). In the absence of K^+^, ATP synthase relies only on H^+^, thus obligating a greater contribution of the ΔpH component of the PMF to ATP synthesis (as evidenced by an additional matrix alkalization of 0.3 units which is not observed in the presence of K^+^) (Figure 3O, bottom; see also Discussion). The presence of any significant inner membrane K^+^ leak pathway (e.g., through an unspecified channel) can be ruled out as an explanation for this K^+^ influx since it would have significantly uncoupled OxPhos, lowering RCR and P/O ratio in the presence of K^+^. Instead, OxPhos coupling was unaltered by K^+^ presence, and the data is entirely consistent with K^+^ transport through the ATP synthase driving the observed increase in ATP synthesis. Thus, there is excellent agreement between the functional data obtained from purified F_1_F_o_ single molecule experiments and ATP synthase studied in the intact mitochondrion. Altogether, these results are fully consistent with predictions arising from experiments performed with purified ATP synthase reconstituted into proteoliposomes and lipid bilayers. It is important to point out that this tight consistency of results obtained between the technically diverse range of experiments presented here (i.e., from single molecule to intact organelle approaches) *rules out* that the behavior of purified, isolated F_1_F_o_ develops an artifactual K^+^ conductance (e.g., due to some *hypothetical* loss of an important regulatory component, etc.), because ATP synthase undoubtedly remains naturally and functionally unaltered in the intact mitochondrial experiments.

### F_1_F_o_-mediated mitochondrial K^+^ influx regulates respiration and mitochondrial volume in cells

The data presented so far indicate that K^+^ flux-driven ATP synthesis can proceed with PL- and single molecule-reconstituted purified F_1_F_o_, as well as in isolated mitochondria, in addition to that normally achieved directly by the flux of a H^+^ gradient. Additionally, in the presence of KCOs F_1_F_o_ achieves a proportionally greater flux of K^+^ ions suggesting that it may function as a recruitable mK_ATP_.

To investigate whether these findings also apply to living cells we examine each one of the following manifestations of mK_ATP_ activation in cardiomyocytes in response to KCOs, based on previous work by others (Garlid et al., 2003; Halestrap, 1989; Sato et al., 1998) and our group (Juhaszova et al., 2004; Juhaszova et al., 2009): (1) flavoprotein (FP) oxidation, (2) modulation of mitochondrial regulatory swelling (i.e., due to mitochondrial K^+^ accumulation), (3) volume activation of respiration (as a consequence of #2), (4) inhibition of GSK-3β activity via ser-9 phosphorylation, and (5) increased mPTP reactive oxygen species (ROS)-threshold. We tested mK_ATP_ activation by Dz in myocytes with IF_1_ knocked down by ~75% through gene silencing (Figures 4B,C), compared to cells treated with control siRNA (Figures 4A,C. Dz produced an equivalent increase in FP oxidation in control myocytes (Figure 4D) as compared to a blunted FP response from IF_1_ siRNA treated cells (Figure 4E), consistent with F_1_F_o_ functioning as a mK_ATP_ regulated by IF_1_. KCO-driven activation of mK_ATP_ causes mitochondrial swelling (Garlid et al., 2003) and increases respiration (Halestrap, 1989; Korge et al., 2005). Using a single cardiac myocyte imaging technique (Juhaszova et al., 2004), we found that KCO Dz, HOE694 (HOE; NHE-1 inhibitor), and the δ-opioid peptide, DADLE, each cause a rapid ~2.5-4% increase in the average volume of mitochondria throughout the cardiomyocyte and increase in respiration (Figures 4F-K,L). In cardiac myocyte suspension, we found that pharmacologic agents that cause mitochondrial swelling (Dz, HOE, and DADLE) increased oxygen consumption (VO_2_) over baseline by about 10%, 25-30%, and 35%, respectively, when utilizing glucose, the medium- and long-chain fatty acid octanoate or palmitate, respectively, and that by preventing this volume increase (e.g., using the Cl^−^ channel inhibitor, IAA-94), the accompanying increase in respiration was similarly eliminated (Juhaszova et al., 2004). Thus, volume activation of respiration is a direct correlate of mitochondrial regulatory volume swelling. Using the same logic as the preceding section, since DADLE causes similar and rapid increases in respiration (as Dz) but is known to *not* activate the mK_ATP_, it was employed as a negative control in the next series of experiments. We found that Vent completely prevented the Dz-related increase in cardiomyocyte swelling and respiration (Figures 4I,L), while the actions of HOE or DADLE were unaffected (Figures 4J-L). Thus, only the specific effect of Dz acting through the mK_ATP_ causes mitochondrial swelling leading to an increase in respiration, but not that of DADLE, requires the function of F_o_.

**Figure 4.**
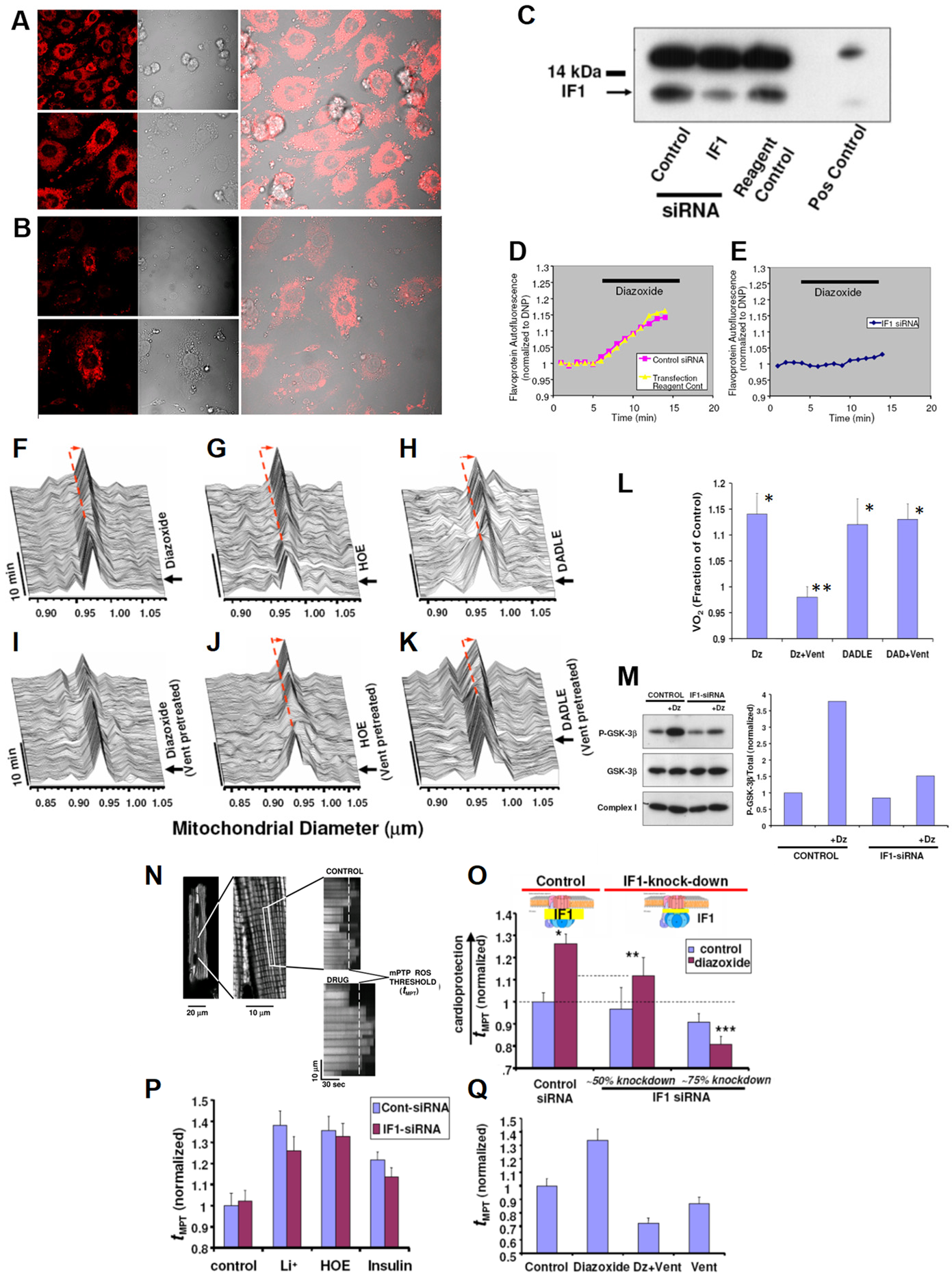
Knockdown of IF_1_ expression in neonatal cardiomyocytes using RNA interference. (**A**) IF_1_ immunocytochemical labeling of control, and (**B**) IF_1_ siRNA treated cells. (**C**) Western blot analysis of control vs siRNA treated samples; positive control corresponds to adult rat heart. (**D;E**) FP autofluorescence (normalized to dinitrophenol, DNP) as marker of mK_ATP_ activity. (**D**) Dz induced FP oxidation in control, and (**E**) No effect of DZ was observed in IF_1_ siRNA treated cells. **(F-K)** *In situ* monitoring of the amplitude and kinetics of regulatory mitochondrial swelling (resulting from increased mitochondrial K^+^ influx and/or retention) in intact cardiomyocytes, based on Fourier analysis of laser linescan transmittance imaging. (**F**) KCO, Dz; (**G**) The NHE-1 inhibitor, HOE, and (**H**) the δ-opioid peptide, DADLE, induced mitochondrial swelling. (**I**) The F_o_ inhibitor, Vent, blocked Dz-induced mitochondrial swelling, while it had no effect on swelling induced by (**J**) HOE or (**K**) DADLE. Arrow indicates the time point of drug addition. (**L**) Mitochondrial respiration (indexed by oxygen consumption, VO_2_ with respect to Dz and Vent treatment as in (**D**) in myocytes. (**M**) mK_ATP_ (Dz)-protection signaling via GSK-3β requires IF_1_ (see text for details). While the KCO, Dz, causes a robust increase in P-GSK-3β in control cells, this was largely prevented in IF_1_-siRNA treated cells. (**N**) Measurements of the mitochondrial permeability transition ROS threshold (t_MPT_, the index of cardioprotection) in myocytes (used in **O-Q**); typical positive t_MPT_ effect of a drug is illustrated vs Control. (**O**) t_MPT_ decreases in proportion to the degree of IF_1_ knock-down, compared to control cells. (**P**) GSK-3β-dependent protection signaling which does not require mK_ATP_ activated K^+^ flux (i.e., Li^+^, HOE, insulin) is unaffected by IF_1_-knock-down. (**Q**) Block of F_o_ by Vent prevents mK_ATP_ (Dz)-mediated cardioprotection. * P<0.05 vs paired Control; **,*** P=ns vs paired Control.

### Effects on mPTP ROS-threshold

The mPTP is a key end-effector of protection signaling: the threshold for mPTP-induction by ROS being significantly reduced after ischemia-reperfusion injury and contributing to cell death, but beneficially increased by preconditioning, postconditioning and other forms of protection signaling, contributing to cell survival (Juhaszova et al., 2004; Juhaszova et al., 2009). We showed that cell protection involves convergence of a multiplicity of potential and distinct upstream pathways (including opening of mK_ATP_), each acting via inhibition of GSK-3β on the end effector, the mPTP complex, to limit its induction (see Figure 4M). We have found that Dz, HOE, Li^+^ (the direct pharmacologic inhibitor of GSK-3β), and insulin, each cause a significant increase of the ROS-threshold for mPTP induction, **t_MPT_** (Juhaszova et al., 2004) (Figures 4N-Q). Since HOE, Li^+^ and insulin each cause protection via mK_ATP_-independent mechanisms they were employed as negative controls in the next series of experiments. The degree of protection (i.e., prolonged **t_MPT_**) afforded by HOE, Li^+^, and insulin was largely unaffected by IF_1_-knockdown (Figure 4P). In stark contrast, the effect of Dz was decreased in direct relation to the degree of IF_1_-knockdown, i.e., **t_MPT_**-increase was reduced by about half with ~50% IF_1_-knockdown, and completely abolished with ~75% IF_1_-knockdown (Figure 4O). We conclude that mK_ATP_-related protection signaling to the mPTP requires the functional presence of IF_1_, thus implicating the role of ATP synthase. Furthermore, similarly to IF_1_-knockdown, Vent blocked the protection by Dz (Figure 4Q). However, while blockage of F_o_ by Vent prevents mK_ATP_ (Dz)-mediated cardioprotection, it does not do so in the case of DADLE or HOE.

### Regulation of F_1_F_o_ by Bcl-xL and MCl-1

Suspecting that the effect of Dz and pinacidil via IF_1_ could be naturally operating under the control of yet-to-be discovered endogenous ligands of IF_1_ we set out to find them. We examined IF_1_ for conserved survival protein-related homology domains since IF_1_ is known to have a “minimal inhibitory domain” sequence of 33 amino acids that binds to the β-subunit of F_1_ (Gledhill et al., 2007). We found that IF_1_ contains a conserved BH3-like domain (residues 32-46) that significantly overlaps its minimal inhibitory sequence (residues 14-47) (Figure 5A), and that Bcl-xL and Mcl-1, which are each known to have a BH3-binding groove, exert effects comparable to Dz on the H^+^ and K^+^ ion currents sustained by ATP synthase (Figures 5B,C). Furthermore, the effect of Bcl-xL and remarkably also of Dz, are reversed by a 26 AA peptide consisting of the BH3-domain of Bad (BH3 peptide, known to have nM affinity for Bcl-xL, but 1-2 orders less so for Bcl-2 (Kelekar et al., 1997; Petros et al., 2001)). A single AA substitution in the BH3 peptide (L12A), that reduces the affinity for Bcl-xL by almost 2 orders of magnitude (Wang et al., 2000), eliminated the inhibitory effects (Figure 5C). Notably, unlike Bcl-xL, Bcl-2 has no effect on F_1_F_o_ H^+^ and K^+^ ion currents, which agrees with their known affinities for the BH3-domain of Bad, respectively. In binding experiments measuring changes in intrinsic tryptophan fluorescence, we found that Bcl-xL has a high affinity (sub-nM K_d_) for the ligand IF_1_, whereas Bcl-2’s affinity is several orders of magnitude lower (see Supplement, section Protein binding (K_d_) measurements). This data suggests that IF_1_ harbors a functionally-active BH3 domain homologous to that of Bad that overlaps with part of IF_1_’s inhibitory domain and functions as the area of binding to the β-subunit of F_1_. Additionally, Bcl-xL and Mcl-1, but not Bcl-2, serve as endogenous regulatory ligands of ATP synthase via interaction with IF_1_ at the BH3-like domain.

**Figure 5.**
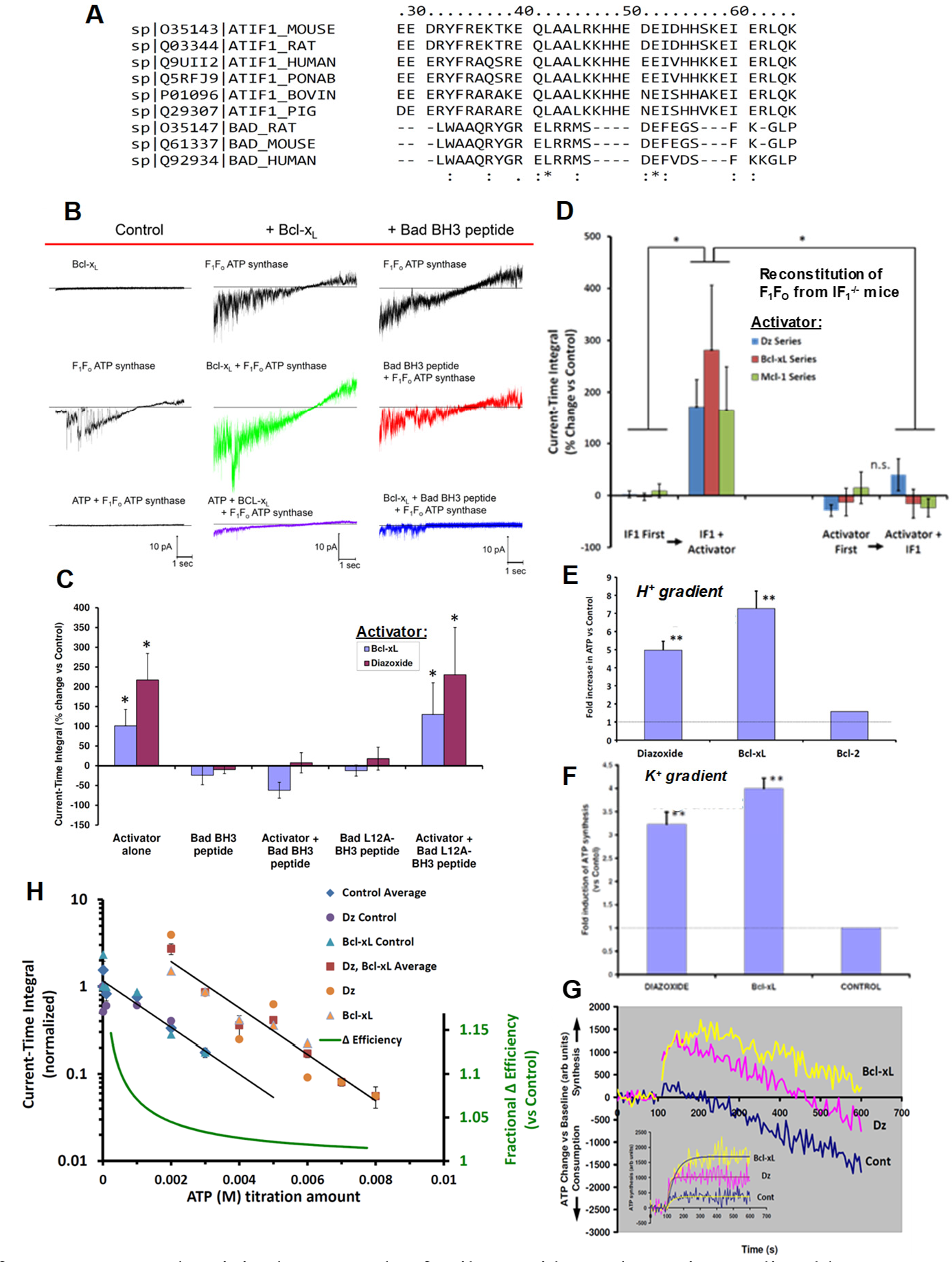
Regulation of F_1_F_o_ current and activity by Dz, Bcl-2 family peptides and proteins mediated by IF_1_. (**A**) AA alignment of the 26-residue BAD BH3 peptide and F_1_F_o_ inhibitory factor IF_1_ from mouse, rat, bovine, pig, monkey and human. The consistency of the alignment is indicated in the last row (asterisk shows complete conservation) (see also Fig S6). BH3 peptide L12 aligns with L42 in full length IF1. (**B**) Effect of Bcl-xL (20 nM) and BH3 peptide (20 nM) on the voltage ramp (from −60 mv to +60 mV) evoked F_1_F_o_ currents. Column headers about the red line denote three experimental groups; Bcl-xl and BH3 peptide middle and right columns, respectively; control currents (left column). Bottom traces (left and middle) correspond to 2 mM ATP inhibited F_1_F_o_ current. (**C**) Augmentation of the current-time integral of voltage ramp evoked F_1_F_o_ currents by Bcl-xL and Dz (30 µM) is reversed by a 26 AA peptide consisting of the BH3-domain of Bad. Control peptide with a single AA substitution L12A has no effect. (**D**) Reconstitution of F_1_F_o_ from IF_1_^−/−^mice. Addition of IF_1_ (100 nM) restores the stimulatory effect of Dz, Bcl-xL and Mcl-1 on the current-time integral of voltage ramp evoked F_1_F_o_ currents from IF_1_^−/−^ mice. The order of addition of IF_1_ and Dz, Bcl-xL or Mcl-1 is varied among the various groups as indicated. (**E**) F_1_F_o_ activity (ATP synthesis) driven by a H^+^ or (**F**) a K^+^ gradient in PL. (**G**) ATP production/consumption kinetics (chemiluminescence traces) in a K^+^ gradient in PL. (**H**) Dose-response of ATP inhibition of F_1_F_o_ (H^+^) currents and Dz and Bcl-xL activated F_1_F_o_ currents: *x-axis-(linear)* ATP concentration used for inhibition of F_1_F_o_ currents, *y-axis (log)* normalized current-time integral of F_1_F_o_ currents. Dz and Bcl-xL produced a parallel shift in the F_1_F_o_ activity vs control resulting in the energy of an additional ~2.8 mM ATP being required to provide sufficient counter-torque to limit F_1_F_o_ to the same level of function as under control conditions. The relative change in efficiency was calculated as the ratio of the free energy of ATP hydrolysis during activation by Dz and Bcl-xL over that under basal conditions. * P<0.05, ** P<0.02.

### Regulation of ATP synthase by Bcl-xL, Mcl-1 and Dz requires IF_1_

Thus far, we have discussed the role and function of IF_1_ in intact cells, organelles and purified single molecules of F_1_F_o_, as well as in IF_1_-knockdown experiments. Next, we examined the regulation of F_1_F_o_ by Bcl-xL, Mcl-1 and Dz in the absence of IF_1_, and upon reconstitution with IF_1_. We measured H^+^ and K^+^ currents in F_1_F_o_ isolated from IF_1_^−/−^ mice (see Supplement section: Generation of IF_1_^−/−^ mice, and Figure S5 regarding confirmatory proof of lack of IF_1_ expression) and the baseline properties of the ionic currents were essentially like that of WT control. Importantly, upon reconstitution with IF_1_, *P*_*H*_ and *P*_*K*_ as well as total current reversal potential were found to be unchanged. One notable difference, however, was the ability of graded mM ATP amounts (by the energy transferred via its hydrolysis) to produce sufficient mechanical counter-torque (exerted by F_1_ on the γ shaft) in excess of the oppositely-directed electrogenic mechanical torque exerted by F_o_, causing a net reversal of electrical current in the IF_1_^−/−^ case (resulting in ATP hydrolysis-generated reverse ion pumping). The latter (i.e., ATP-generated reverse ion pumping) was not observed in parallel experiments with WT and is entirely consistent with the known function of IF_1_ to limit the waste of futile ATP hydrolysis by impaired mitochondria under circumstances when ΔΨ_m_ would drop below levels needed to synthesize ATP (Walker, 1994).

Assessing the current-time integral function (CTI, which in the direction of negative current is the direct analog of the amount of ATP synthesized) after reconstitution of F_1_F_o_ with IF_1_ in IF_1_-/-, we observed a small *increase* of ~11-14% in CTI (*vs* baseline). Thus, it is notable that IF_1_ does not cause a *net* inhibitory drag on the energy transfer in F_1_F_o_ likely due to frictional losses in the direction of ATP synthesis (see also discussion below regarding BH3 peptide effects). In contrast to WT, neither Dz, Bcl-xL nor Mcl-1 exerted a significant positive augmentation of CTI in the absence of IF_1_, and subsequent IF_1_ reconstitution was similarly ineffective (Figure 5D). A likely explanation for the apparent ineffectiveness of IF_1_ added after Bcl-xL, Mcl-1 or Dz can be given by the possible interference exerted by these molecules on the intrinsic disorder of IF_1_ (Bason et al., 2014), hindering its interaction at the F_1_’s binding cleft and γ shaft into a functionally active complex. In one case, the 5-fold excess of IF_1_ used leaves effectively no free Bcl-xL because of the high affinity of this pair, and presumably only free IF_1_, rather than bound, can reconstitute into F_1_F_o_. In the case of Dz, this molecule could directly affect the intrinsic disorder of IF_1_ or the binding cleft preventing effective reconstitution. On the other hand, prior reconstitution of F_1_F_o_ with IF_1_ restores the WT behavior entirely, with Dz, Bcl-xL and Mcl-1 manifesting a robust augmentation of CTI (Figure 5D). Taken together, this data allows us to conclude that IF_1_ is required for these mediators to augment F_1_F_o_ activity (for the same driving force).

As stated earlier, the positive effects of Bcl-xL and Mcl-1 on WT F_1_F_o_ currents could be reversed by the BH3 peptide (but not by the L12A variant, null-acting control BH3 peptide). Since this BH3 peptide, as well as IF_1_, likely binds to the same region of F_1_-β, but because of its short length is unable to reach to the γ shaft, we examined the functional effects upon binding F_1_F_o_ in the absence of IF_1_. We found that in IF_1_-deficient F_1_F_o_, the BH3 peptide alone exerts a robust positive effect comparable to that of Dz, Bcl-xL and Mcl-1 (i.e., doubling to tripling the activity; not shown), but the L12A-modified BH3 peptide had no effect, suggesting that IF_1_ likely produces significant frictional drag via its constitutive contact with the γ shaft that is fully offset by some function-augmenting mechanism achieved by the portion of IF_1_ bound to F_1_-β (see below and Discussion).

### Regulation of mechano-chemical efficiency of F_1_F_o_

We have shown that F_1_F_o_, conducting univalent cations at a fixed driving energy, Δμ, can be upregulated to increase the total ion flux against a constant load without slip or leak via the IF_1_-dependent actions of synthetic small molecules such as Dz and pinacidil, and endogenous proteins such as Bcl-xL and Mcl-1. The absence of slip is revealed by complete inhibition of currents at high Δμ by excess ATP. This activity enables ATP synthase to function as a recruitable mK_ATP_, whereby the triggered increase of mitochondrial K^+^ influx and matrix volume upregulate respiration and produce redox activation of local signaling inhibiting GSK-3β and resulting in desensitization of the mPTP to damaging levels of ROS (Juhaszova et al., 2004; Zorov et al., 2014). These data, together with the results showing that Bcl-xL and Dz (and pinacidil and Mcl-1) are each capable of increasing the amount of ATP synthesized by reconstituted F_1_F_o_ (WT, IF_1_-competent) utilizing either K^+^ or H^+^ gradients (Figures 5E-G), suggest that these IF_1_-dependent effectors have increased the mechano-chemical efficiency of the ATP synthase. To investigate this, we examined the titration curve of the CTI (at each of the ion-reversal potentials for H^+^ and K^+^, in single ATP synthase molecules) as a function of the counter-torque on the γ shaft applied by F_1_ resulting from the hydrolysis energy derived from increasing ATP concentrations, in the presence of Dz or Bcl-xL as compared to controls. The data obtained are well described by a log-linear relationship between CTI and [ATP]. Dz and Bcl-xL produced a parallel upward shift of 5.6-fold in the F_1_F_o_ activity *vs* control (Figure 5H) indicating that the hydrolysis energy of an additional ~2.8 mM ATP is required to provide sufficient counter-torque to constrain the F_1_F_o_ to the same level of function as under control conditions. Based on considerations of energy conservation, the additional ATP synthesis might be driven by extra energy that was not lost to viscous drag and intermolecular friction. Together, these results agree with the idea that both Bcl-xL and Dz increase the mechano-chemical efficiency of ATP synthase (e.g., by ~7% at 1 mM, ~5% at 2 mM, and ~3% at 4 mM ambient ATP).

## Discussion

Up to the present, it has been a central tenet of bioenergetics that mammalian ATP synthase operates solely on proton flux through F_o_ to make ATP. The present work significantly revises that concept. We found that despite the high degree of F_o_’s H^+^ selectivity *vs*. K^+^ (~10^6^:1), the abundance of cytoplasmic K^+^ over H^+^ being >10^6^:1 enables ATP synthase to harness Δμ_K_ (electrical gradient energy) and conduct a significant number of K^+^ for every H^+^ in the synthesis of ATP. Specifically, our finding is supported by the following evidence: 1) under physiological pH=7.2 and K^+^=140 mEq conditions, purified F_1_F_o_ reconstituted in proteoliposomes exhibiting a stable (non-zero) ΔΨ_m_ in the presence of the K^+^ gradient, can synthesize ATP solely driven by the free energy stored in the experimentally-defined K^+^ gradient by up to ~3.5 K^+^ for every H^+^; 2) purely K^+^-driven ATP synthesis from single F_1_F_o_ molecules measured by bioluminescence photon detection could be directly demonstrated along with simultaneous measurements of unitary K^+^ currents by patch clamp, both blocked by specific F_o_ inhibitors, Vent/Oligo; 3) in the presence of K^+^, compared to osmotically-matched conditions in which this cation is absent, isolated mitochondria display 3.5-fold higher rates of ATP synthesis, at the expense of 2.6-fold higher rates of oxygen consumption, these fluxes being driven by a 2.7:1 K^+^:H^+^ stoichiometry; the excellent agreement between the functional data obtained from purified F_1_F_o_ single molecule experiments and ATP synthase studied in the intact mitochondrion under unaltered OxPhos coupling by K^+^ presence, is entirely consistent with K^+^ transport through the ATP synthase driving the observed increase in ATP synthesis; 4) the chemo-mechanical efficiency of ATP synthase can be endogenously up-regulated by certain members of the Bcl-2 family, and pharmacologically by certain K^+^ channel openers, acting via IF_1_, an intrinsic regulatory factor of ATP synthase, in a process that increases the monovalent cation conductance of F_1_F_o_ while retaining its high degree H^+^-selectivity.

As the currency of metabolic energy, the flux of ATP catalyzed by ATP synthase generates every day (in humans) ~2 × 10^26^ ATP molecules, corresponding to a mass of about 80 kg. In other words, each day we synthesize and recycle the equivalent of our own body weight in ATP (Rich, 2003). In the human heart, the estimated daily amount of ATP generated (6 – 35 kg) is much more than its own weight (~300 g) (Ashrafian et al., 2007; Taegtmeyer, 1994). This entails an extraordinary process of energy supply-demand matching which occurs at extremely high rates. The increase in energy demand from rest to maximal can be 5- to 10-fold in normal people, depending on the intensity of physical activity, and reach 20-fold in well-trained human athletes (Weibel and Hoppeler, 2005). We propose that the dynamic range of ATP synthesis flux enabled through the newly discovered K^+^ mechanisms described here are essential to fulfill to a great extent the broad span of energy demand needs, matched by the supply of a highly efficient mechanism of energy generation, and evolutionarily tuned by endogenous regulators from the Bcl2 family of proteins.

Although what we propose remains fully compatible with Mitchell’s chemiosmotic mechanism ((Mitchell, 1961); reviewed in (Nicholls and Ferguson, 2013)), our findings have major bioenergetic implications. Electrophysiological measurements indicate that purified F_1_F_o_ reconstituted into the lipid bilayer could conduct up to 3.7 K^+^ for every H^+^, in the absence of any other K^+^ conducting pathway as demonstrated in section **Measurement of unitary K^+^ and H^+^ currents from F_1_F_o_**. The demonstration that F_1_F_o_ in isolated mitochondria utilizes ~ 3 K^+^ for every H^+^ transferred, yielding ~2 K^+^ per ATP (based on 8 ions driving the *c*-ring per 3 ATPs in mammalian F_1_F_o_ together with our measurement of ~3:1 K^+^:H^+^), indicates that ATP synthase acts as a K^+^-uniporter, i.e., the primary way for K^+^ to enter mitochondria. In isolated mitochondria the major fraction of the total K^+^ flux (likely significantly exceeding 60% considering the accompanying large differences in matrix volume changes) is sustained by the ATP synthase (Figures 3G-O; Figure S4D), thus showing that F_1_F_o_ is a major mitochondrial K^+^ influx pathway. Since K^+^ entry is directly proportional to ATP synthesis and regulates matrix volume and respiration, in turn it directs the matching between energy supply and demand. The K^+^ flux can be enhanced, halted, or even reversed depending on ATP concentration based on thermodynamic energy balance (and the function of IF_1_).

We show that F_1_F_o_, conducting H^+^ and K^+^, can be upregulated (even at the same driving energy, Δμ) to increase the total ion-flux (at constant H^+^:K^+^) against a constant load without slip or leak via the IF_1_-dependent actions of endogenous pro-survival proteins, Bcl-xL and Mcl-1, and of synthetic small molecules, Dz and pinacidil (Figure 5; conceptualized in Figure 6C). By harnessing Δμ_K_, driven essentially by ΔΨ_m_, and continuously converted (restored) from respiratory chain-generated Δμ_H_ through the activity of the KHE, F_1_F_o_ generates additional ATP proportional to the amount of energy that would have been dissipated as heat by the same K^+^ current in passing (in a hypothetical scenario) through a separate entity functioning only as a K^+^ uniporter. In other words, letting K^+^ enter via a non-ATP generating process would not be as energetically effective as using the F_1_F_o_ as the K^+^-influx mechanism. Thus, once the K^+^ is eventually extruded by the KHE using H^+^ influx, the equivalent energy of that H^+^ will have been harnessed in form of ATP made by the K^+^ influx through the F_1_F_o_ (Figure 6B).

**Figure 6.**
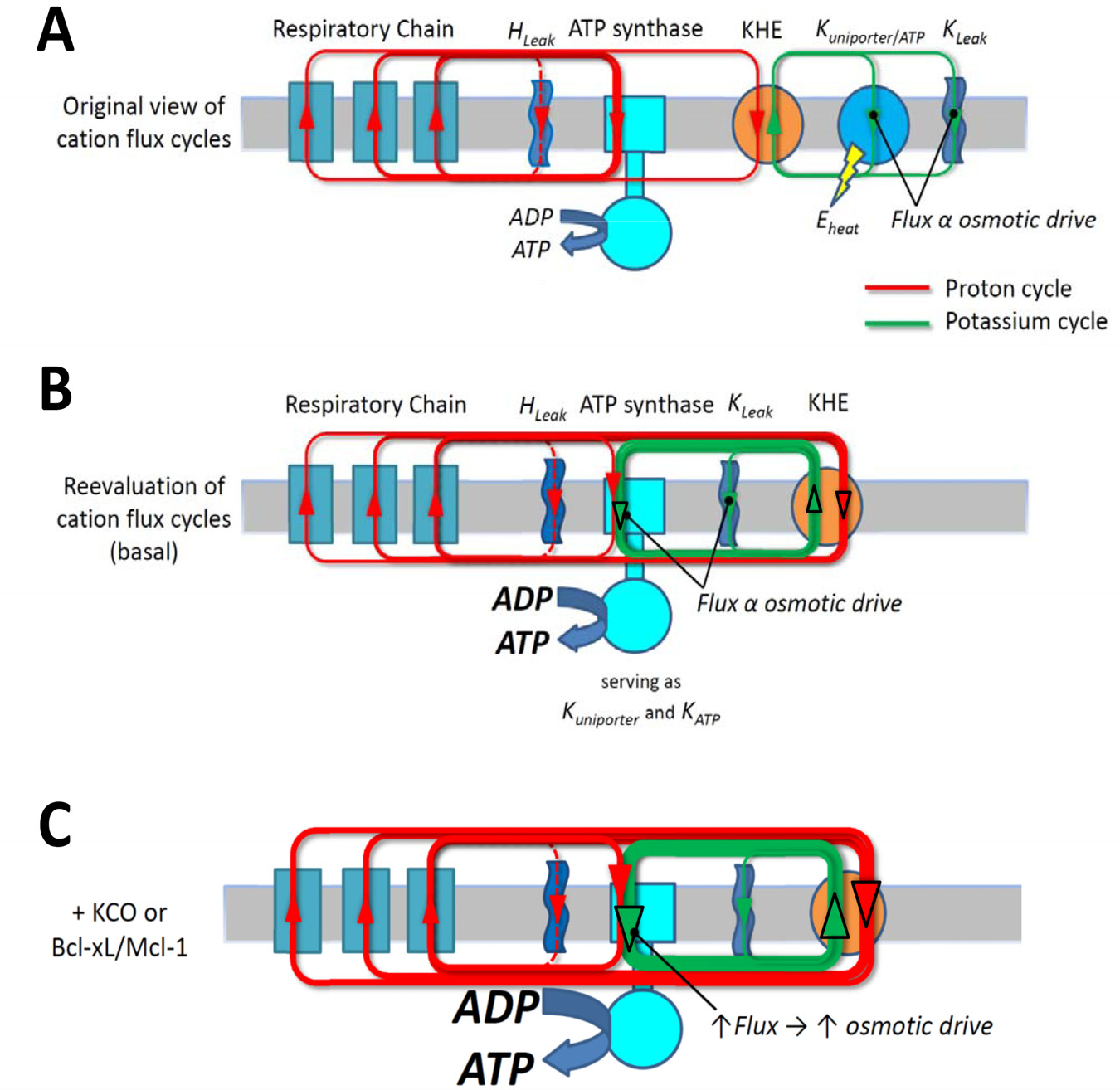
Scheme of the H^+^ and K^+^ transport across the inner mitochondrial membrane. From the energetic standpoint, all the energy available to perform work and execute the ionic movements derives from the original H^+^ gradient established by proton pumps in the respiratory chain. A central point is the obligatory preservation of charge and mass balance under the steady state circuits. In the “original view of cation flux cycles” (**A**), a certain (majority) of the H^+^ gradient is being harnessed by F_1_F_o_ directly to make ATP, whereas a certain amount of K^+^ is entering the matrix through an ordinary K^+^ channel mechanism (a “mK_ATP_-uniporter” channel), driven by ΔΨ, and extruded via KHE utilizing the energy remaining in the fraction of the H^+^ gradient not harnessed by F_1_F_o_. The equivalent energy of this fraction being used to extrude K^+^, and a large fraction of that non-ATP-producing energy would essentially be dissipated as heat in the constant cycle of K^+^ recirculation (green circuit in A). In the new mechanism (**B**) the same amount of energy available in the original H^+^ gradient but largely lost as heat is entirely available to produce ATP, simply by having the mK_ATP_-uniporter mechanism reside inside, and as natural part of, F_1_F_o_ with the traffic of H^+^ or K^+^ contributing its energy to producing ATP. The remainder of the H^+^ gradient energy is now utilized to remove all the K^+^ that entered via F_1_F_o_ (n.b., this exchange process of extruding K^+^, restoring Δμ_K_, is the way that the H^+^ gradient energy is still the original, entire driving force for ATP production). However, the gain is that more ATP is produced for the same input energy by not wasting some of that energy on maintaining what was originally thought to be a separate K^+^ cycle that does not/cannot generate any ATP. Engineered this way, it is a better, tightly coupled system of energy supply-demand matching through the K^+^ cycle utilizing F_1_F_o_ because the matrix influx of K^+^ is truly directly proportional to ATP synthesis. Any transient increase in F_1_F_o_ activity will thus lead to transient K^+^ accumulation (due to a natural kinetic lag in the activity increase in KHE reacting to the matrix K^+^ accumulation). This will lead to the attraction of a counter-ion and change of the osmotic drive yielding a “volume-activation of respiration” response which previously has been documented in detail (Juhaszova et al., 2004). The scheme depicted in (**C**) integrates the implications of modestly enhancing the chemo-mechanical efficiency of F_1_F_o_ (by KCO’s or Bcl-xL/Mcl-1). For the driving energy of the same H^+^ gradient the F_1_F_o_ flux increases, enabling increased respiration and a directly increased K^+^ flux cycle (yielding an increased volume signal) and enhanced ATP generation (**C**) vs the basal conditions (B).

Because a transient change in K^+^ influx would need to be matched by influx (or retention) of a counter-ion (e.g., Cl^−^) to produce an osmotic imbalance signal, both KHE and the counter-ion transport pathways are also important control steps in matrix volume regulation. Dysfunction of mitochondrial KHE activity leads to aberrations in matrix K^+^ and mitochondrial volume regulation that in turn may affect fission/fusion and mitophagy. Such pathology is evident in the Wolf-Hirschhorn syndrome, a genetic insufficiency of mito-KHE activity (1/50,000 incidence, characterized by microcephaly, growth retardation, intellectual disability, and epileptic seizures among other severe manifestations (Zotova et al., 2010)).

The K^+^ uniporter function also enables F_1_F_o_ to operate as an on-demand, recruitable mK_ATP_, whereby triggered increases of mitochondrial K^+^-influx and matrix-volume upregulate the signaling cascade resulting in desensitization of the mPTP, enhancing cell survival (Juhaszova et al., 2004). Nature usually operates important pathways with built-in redundancy so that other mitochondrial K^+^ channels may contribute to these mechanisms, including the Ca^2+^-activated K^+^ channel, BK_Ca_ (Xu et al., 2002), and a ROMK channel which may also mimic mK_ATP_ channel function (Foster et al., 2012), but these pathways are likely fine-tuning mechanisms.

Our data also unveil that F_1_F_o_ operates at increased efficiency (by up to ~7% at normal ATP levels) in response to KCOs, Bcl-xL and Mcl-1, yielding both increased ATP output and matrix K^+^ influx for the same Δμ_H_ (Figure 5; depicted in 6C). Dz and Bcl-xL cause a rightward shift in the ATP-dependence of the CTI (a quantitative index of ATP synthesis), such that the hydrolysis energy of an additional ~2.8 mM ATP is required to provide enough counter-torque to constrain the F_1_F_o_ to the same level of function as in controls. This means that an additional ~2.8 mM ATP can be produced for the same input energy at normal ambient levels of ATP. This provides quantitative proof that both Bcl-xL and Dz increase the mechano-chemical efficiency of F_1_F_o_ (Figure 5H). Recent work found that Bcl-xL interacts with the F_1_F_o_ (Alavian et al., 2011; Formentini et al., 2014), specifically with the β-subunit of ATP synthase decreasing an ion leak within the F_1_F_o_ complex and concluded that this was responsible for increasing net transport of H^+^ by F_1_F_o_ (Alavian et al., 2011). These latter findings and conclusions are non-trivially different from our experiments: (1) we do not observe ion leak (or slip) at all, regulated or otherwise, in F_1_F_o_ in the presence or absence of Bcl-xL, i.e., Bcl-xL does not inhibit an ion leak that is not present in ATP synthase, and (2) the increase in ATP synthetic capacity in response to Bcl-xL is specifically due to an increase in mechano-chemical efficiency of ATP synthase *per se*, and not by changing an ion leak into useful energy. These latter findings lead us to conclude that essential mitochondrial homeostatic and pro-survival mechanisms result from a regulated IF_1_-mediated increase in chemo-mechanical efficiency of F_1_F_o_ conducting both K^+^ and H^+^. Our results add a significant dimension to the known, and apparently diverse biological function sets of F_1_F_o_. Additionally, it was proposed that a certain triggered rearrangement of F_1_F_o_ dimers is functionally responsible for other major biological functions such as the mitochondrial cristae arrangements (Strauss et al., 2008) and possibly the formation of the mPTP (Giorgio et al., 2013).

Our findings raise the question of how IF_1_ might control the activity of ATP synthase to engage physiologic/homeostatic and survival-promoting mechanisms. Overall, our data are consistent with a “minimal inhibitory domain” of IF_1_ (residues 14-47 in bovine IF_1_ (van Raaij et al., 1996)) binding to the β-subunit of F_1_ in an “IF_1_ ligand-binding cleft” (adjacent to the F_1_ α-subunit interface), forming at its proximal end an α-helix loop that interacts with the F_1_ γ-rotor shaft which is responsible for limiting ATPase activity. With the evidence of a significant modulatory role by certain Bcl-2 members, we examined this domain for conserved survival protein-related homology domains. Bcl-xL and Mcl-1 are each known to have a BH3-binding groove with high affinity for certain domains of BH3. Together with the result of the high affinity binding of IF_1_ to Bcl-xL, our data agrees with IF_1_ harboring a functionally-active BH3-like domain homologous to that of Bad and coincident with IF_1_’s inhibitory domain that functions as the binding patch to the β-subunit of F_1_. Binding of Bcl-xL and Mcl-1, but not Bcl-2, via IF_1_ interaction, endogenously regulate F_1_F_o_ activity. This may explain why the effects of Bcl-xL, Mcl-1, and Dz, are reversed by the BH3 peptide, but not by the same peptide with a single AA change (L12A) (Kelekar et al., 1997; Wang et al., 2000) (Figures 5B,C). Specifically, the BH3 peptide may compete and displace IF_1_ from its binding site on F_1_F_o_, as well as interfere with its binding to Bcl-xL or Mcl-1. Moreover, we have shown that, unlike in WT, neither Dz nor Bcl-xL significantly increased CTI in F_1_F_o_ from IF_1_-/-. Alternatively, prior reconstitution of F_1_F_o_ in the presence of IF_1_ entirely rescued the WT behavior, with both Dz and Bcl-xL strongly augmenting the ion currents (Figure 5D). These data allow us to conclude that the higher ATP synthase activity elicited by these effectors (for the same driving force) requires IF_1_, and that the mere removal from its binding site does not suffice to enhance the enzyme activity. We propose that in the normal basal state IF_1_ has two mechanical and *nearly offsetting* effects on the function of ATP synthase operating in the synthesis direction: (1) a net *negative*, frictional drag-like effect of the IF_1_ molecule originating at its proximal end where it engages the γ shaft in its natural rotation, and (2) a net *positive* effect created somehow by the presence of the long α-helical stretch that engages the IF_1_ binding cleft on F_1_-β, the latter effect being mimicked by the BH3-peptide. It has been shown that Bcl-xL can interact forming 3D-domain swapped (3DDS) homodimers (O’Neill et al., 2006) as well as heterodimers with other survival-regulating proteins. These interactions can significantly affect the residual function of both partners (Rajan et al., 2015), and certain BH3-only proteins can bind to and partially unfold Bcl-xL, changing its interactions with other binding partners and thereby biasing cell survival-signaling (Follis et al., 2013). Thus our two-fold proposal implies that (1) Bcl-xL/Mcl-1 (via their intrinsic BH3-binding grooves) tightly bind to IF_1_ at its minimal inhibitory/BH3-like domain to displace it from its binding cleft at F_1_-β, and (2) this interaction triggers a specific unfolding and rearrangement of the Bcl protein’s α2 helix, enabling an increase of its potential range-of-motion. This could allow the helix from the Bcl protein to participate in an energetically favorable rearrangement with F_1_F_o_ by binding to the empty IF_1_ binding cleft. We propose a possible model of this interaction (Bcl-protein’s α2 helix containing its BH3 domain engaging the IF_1_ binding cleft on F_1_-β; Figures7A-D) that would cause the Bcl-xL/Mcl-1-mediated increase of F_1_F_o_ function in the presence of IF_1_, analogous to that obtained with the BH3 peptide added to the IF_1_ deficient F_1_F_o_ (Figure 5D). The mechanism by which a short IF_1_/BH3-(like) helical-peptide structure occupying the natural IF_1_ binding groove can enhance the chemo-mechanical efficiency of F_1_F_o_ is of considerable interest, but how it specifically works remains a matter of future study.

The origin of IF_1_ in relation to the evolution of F_1_F_o_ is also an interesting question. There are conserved “IF_1_ domains” that can be found embedded in a variety of larger proteins across Archaea, Bacteria, and Eukaryotes (Geer et al., 2002), suggesting ancient origins for this domain. Although F_1_F_o_ exists in all major lifeforms, IF_1_ as a separate entity is only known to regulate synthase function in Eukaryotes. It is tempting to speculate that when the early bacterium became a mitochondrion as a functional organelle of the eukaryotic cell, some 2 billion years ago, it brought along the genetic information for IF_1_, which might have evolved to prevent the mitochondrion from wasteful ATP consumption in the host cell. We examined these bacterial IF_1_-progenitors and they have regions homologous to the BH3-like domains that we found in eukaryotic IF_1_’s. The Bcl family is also ancient, some 2 billion years extant and resident in eukaryotic lifeforms. Bayesian phylogenetic analysis shows that IF_1_ is an ancient member of the Bcl family and today may be most closely related to BH3-containing proteins (e.g., Bad, PUMA, Bcl-xL; Figure 7E). This may explain how the Bcl-2 protein family has come to regulate F_1_F_o_ function as part of its repertoire of survival-regulating functions.

**Figure 7.**
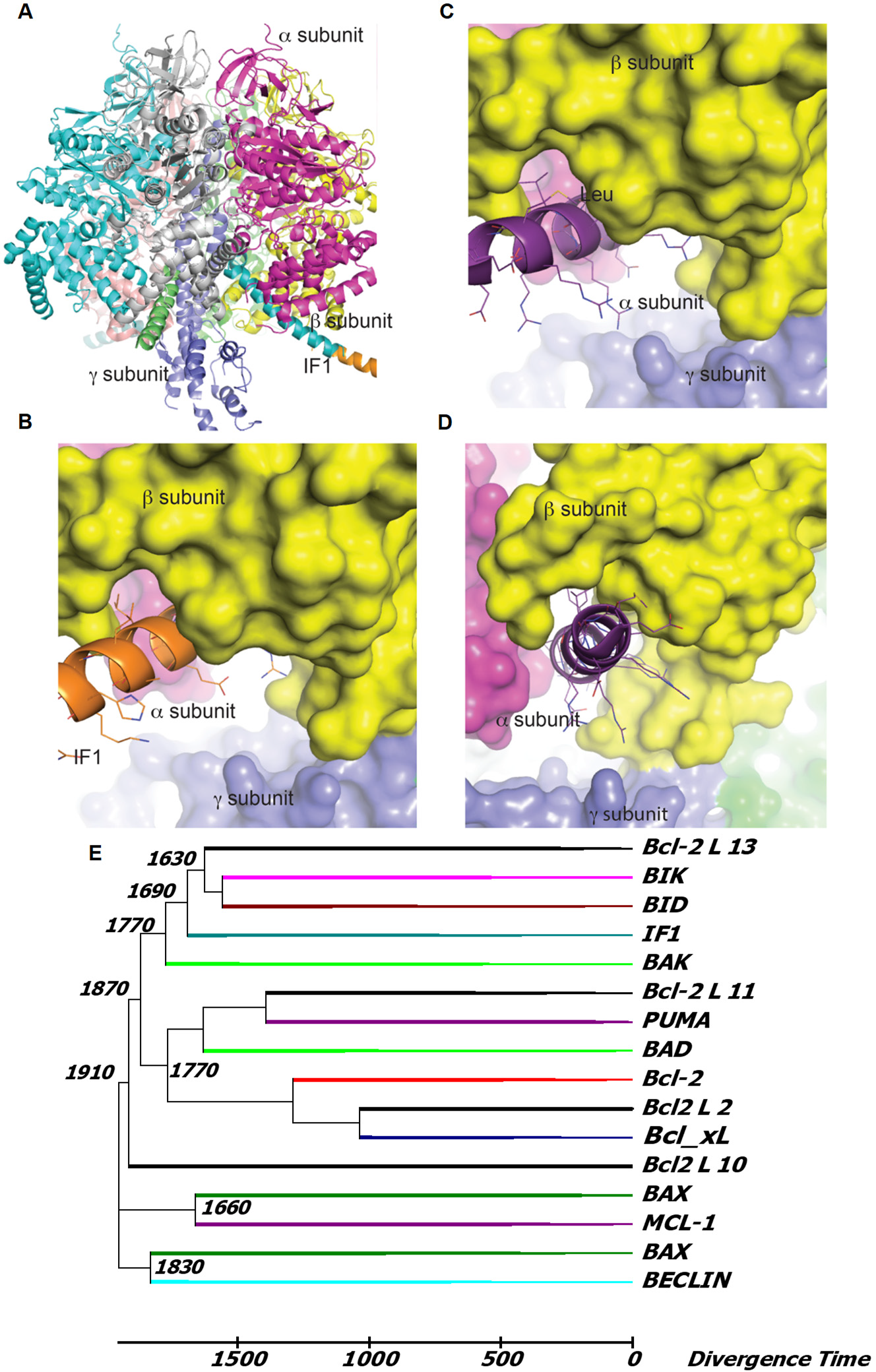
Interaction of F_1_ ATPase with IF_1_ or a BH3 modeled peptide. (**A**) Ribbon model of the crystal structure of bovine F1 (PDB ID 4Z1M) emphasizing two of the three β subunits (cyan and magenta) and the γ subunit (blue) in interaction with the long α-helix of the inhibitor protein IF_1_ (orange). The IF_1_ domain (residues 18-51 are shown in cyan; residues 23-70 from PDB ID 1GMJ are shown in orange) interacts with the β subunit marked in yellow. (**B**) Surface representation of subunits β (yellow), γ (blue) and α (magenta) of F_1_ ATPase interacting with IF_1_ peptide. (**C**) As panel (**B)** with the α_2_-helix peptide containing BH3 domain from the BAD protein (PDB ID 1G5J, as it binds BCL-XL) in ribbon representation at the IF_1_ groove in F_1_ showing the aliphatic side chain of Leu 42 in IF_1_ and Leu 12 in BH3-BAD peptide (corresponding to Leu114 in human BAD). (**D**) Same as (C) at an approximately 90° orientation. (**E**) Phylogenetic tree of the BH3 extended peptides (35 residues) from Bcl-2 proteins and IF1 across eukaryotes. Sequence alignment was computed with Clustal Omega (Sievers et al., 2011) and the tree and divergence times (in Myr) were calculated by MEGA 6.0 (for further details, see Supplemental Information; Figs S6 and S7, and Table S2).

In conclusion, we demonstrated that F_1_F_o_ utilizes the ion gradient energy not only of H^+^ but also of K^+^ to drive ATP synthesis. The essential mitochondrial homeostatic and pro-survival mechanisms discussed here, including F_1_F_o_ operation as a primary mitochondrial K^+^ uniporter to facilitate energy supply-demand matching, and as a recruitable mK_ATP_ channel to protect from pathological opening of the mPTP, result from regulated function of ATP synthase conducting both K^+^ and H^+^. The specific mechanisms by which KCOs and certain Bcl-2 family proteins engage IF_1_ to produce an increase in the chemo-mechanical efficiency of ATP synthase will require additional investigation.

## Experimental procedures

### Detailed methods are available in the Supplemental Experimental Procedures

#### Cell isolation, and purification, characterization and reconstitution of F_1_F_o_

Myocytes were isolated from neonatal and adult rat, and mouse hearts by enzymatic dissociation. Mice carrying an inactivated Atpif1 allele were obtained from the European Mouse Mutant Archive (EMMA), bred to homozygosity and are referred to as IF_1_-/-throughout the text.

F_1_F_o_ was purified according to manufacturer protocol (Mitosciences) and reconstituted into liposomes and planar lipid membranes.

#### ATP measurements

Bioluminescent assays which employ the luciferin-luciferase ATP-dependent reaction were used to evaluate the ATP production by PL.

#### Electrophysiological measurements

The Planar Lipid Bilayer workstations (Warner Instruments) were used to characterize electrophysiological properties of the reconstituted F_1_F_o_.

Simultaneous measurements of unitary K^+^ currents and single photon detection of ATP synthesis activity (bioluminescence) from reconstituted F_1_F_O_ into a lipid hydrogel-hydrogel interface bilayer formed on a 30-50 µm glass pipette are described in **the Supplemental Experimental Procedures**.

#### Membrane potential measurements

The potential-sensitive fluorescent probe oxonol VI was used to monitor F_1_F_O_-generated Δψ in a K^+^ gradient using a PTI spectrofluorometer (Photon Technology International Inc.)

#### Confocal microscopy experiments

mPTP ROS-threshold induction, mitochondrial volume determination, FP autofluorescence measurements and immunoflourescence microscopy were performed as described before (Juhaszova et al., 2004; Zorov et al., 2000).

#### Statistics

All experiments were performed at least in triplicate, with cell number greater than 12 in each independent experiment unless stated otherwise. All data are mean ± SEM. Comparisons within groups were made by an appropriate one-way ANOVA or Student t test, and P value ˂0.05 was considered as statistically significant.

## Supporting information

Juhaszova et al_ supplemental Revised

## Author contributions

Conceptualization, M.J., D.B.Z. and S.J.S.; Methodology, M.J., E.K., H.B.N., K.W.F., L.M., M.A.A., S.C. and S.J.S.; Software, Y.Y., S.B.G. and S.C.; Formal Analysis, Y.Y., S.B.G., S.C. and S.J.S.; Investigation, M.J., E.K., D.B.Z., H.B.N., M.A.A. and S.C.; Resources, R.dC., L.M., S.B.G. and S.J.S.; Writing-Original Draft, S.J.S.; Writing-Review & Editing, M.J., E.K., D.B.Z., K.W.F., R.dC., S.B.G., M.A.A., S.C. and S.J.S.; Visualization, M.J., E.K., Y.Y., S.B.G., M.A.A., S.C. and S.J.S.; Supervision, S.J.S.

## Acknowledgments

We thank E.G. Lakatta for useful discussions, D. Boyer for animal husbandry, L. Rezanka for mice genotyping and M.J. del Hierro Sanchez for assistance in obtaining the transgenic IF_1_^−/−^ mice. This work was supported entirely by the Intramural Research Program, National Institute on Aging, NIH.

## Conflict of interest

The authors declare that they have no conflict of interest.

