## Supplementary material for "ATP synthase K^+^- and H^+^-flux drive ATP synthesis and enable mitochondrial K^+^-uniporter function": Juhaszova et al_ supplemental Revised

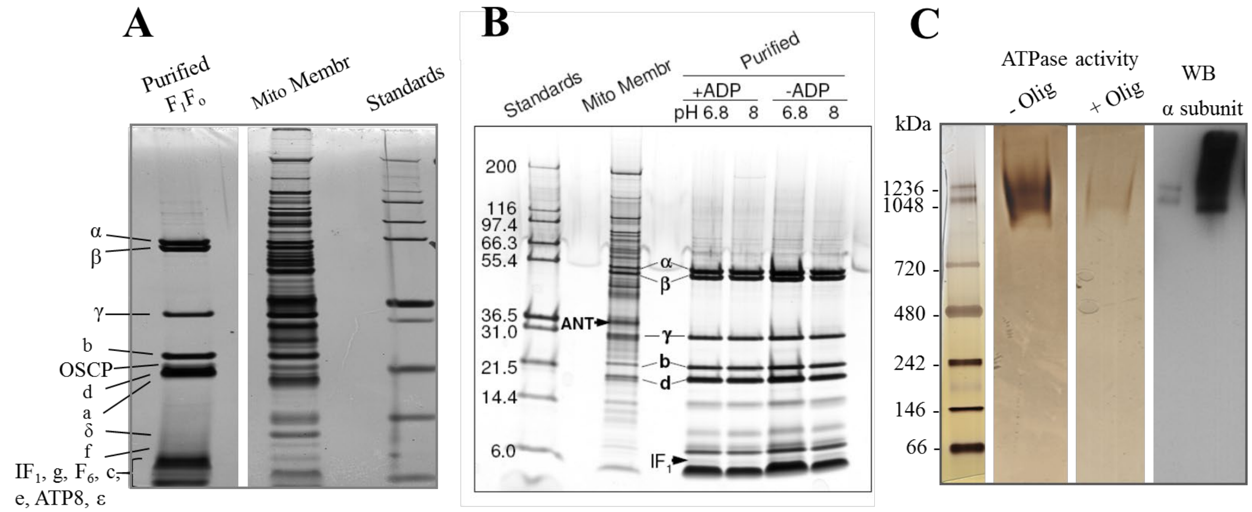


**Figure S1 – related to Figure 1. Purity and protein characterization of the isolated $F_1F_o$ complex. (A,B)** $F_1F_o$ was purified according to manufacturer protocol (Mitosciences) and reconstituted into liposomes and planar lipid membranes. The purity of the enzyme complex was assessed by gel electrophoresis, silver staining and Western blotting (under denaturing conditions), which shows the absence of virtually all other unrelated mitochondrial proteins, including ANT, one of the most abundant inner mitochondrial membrane proteins. **(B)** Importantly, the binding of the small (8 kD) $IF_1$ to the synthase complex is reversible and pH-dependent, with approximately half of the $IF_1$ being bound at pH 8 as compared to pH 6.8. The synthase complexes during and after isolation were maintained at pH 7 to keep $IF_1$ binding, except in protocols for which $IF_1$ was intentionally removed by a short wash in alkaline (pH 10) buffer. **(C)** The isolated $F_1F_o$ showed mostly dimers (possibly even higher order multimers) when run on clear-native gels, and the "in-gel" oligomycin-sensitive ATPase activity, as seen in the clear-native gel, was largely restricted to the dimers and higher.

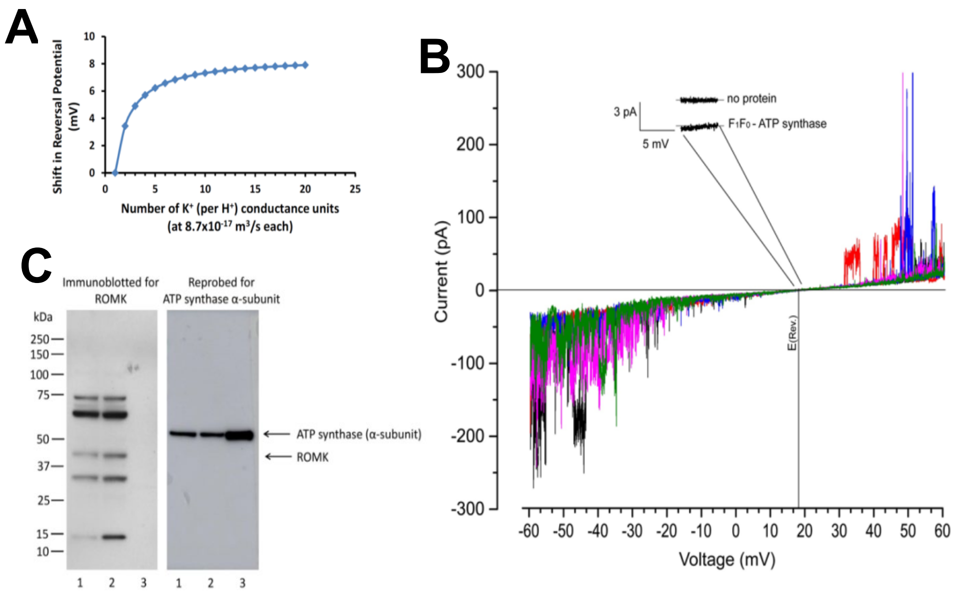


**Figure S2 – related to Figure 1. Absence of functionally relevant $K^+/H^+$ antiport or independent $K^+$ channel activity contaminating the $F_1F_o$ proteoliposome (PL) reconstitution experiments. (A)** Shift in Reversal Potential ($E_{rev}$) as a function of a hypothetical variation in the stoichiometry of $F_1F_o$ $H^+$ conductance units vs number of $K^+$ channels ("$K^+$ conductance" units)**.** All electrophysiological data confirms the absence of coincidental co-immunoprecipitation with a contaminating $K^+$ channel (or a $Na^+$ channel). The results of a calculation based on manuscript equation 1 show a theoretical shift in $E_{rev}$ as a function of a hypothetical variation on the stoichiometry of "$H^+$ conductance" and "$K^+$ conductance" units: note that a ratio of 1:1 yields the obtained $E_{rev}$ in the present experiments (a ratio of 1:0 represents the absence of a "$K^+$ conductance unit" and $E_{rev}$ will equal the reversal potential for $H^+$ which is set by ionic conditions to be zero; this result has never been seen in any of our experiments). A ratio of 2:1 means two "$K^+$ conductance" units per "$H^+$ conductance", and so on. If purified $F_1F_o$ complex displayed a 2:1 ratio we would observe a ~3.5 mV positive shift in $E_{rev}$, which was never observed. That $E_{rev}$ is invariant from experiment to experiment (our resolution is < 1 mV) confirms a fixed 1:1 stoichiometry. Furthermore, the $K^+$ conductance is quantitatively inhibited in parallel with that of $H^+$, without any shifts in $E_{rev}$, by the titration with ATP (Fig 5H). Finally, the "$H^+$ conductance" and "$K^+$ conductance" units are all completely inhibited by both Oligo and Vent. If there was a contaminating $K^+$ (or $Na^+$) channel co-immunoprecipitated with $F_1F_o$, that channel activity would remain after these inhibitors, as we know of no $K^+$ (or $Na^+$) channels that are coincidentally *fully* sensitive and inhibited by Oligo and Vent, but such a residual channel activity never happens. Additionally, given the apparent 1:1 stoichiometry with $F_1F_o$ of such a hypothetical $K^+$ (or $Na^+$) channel, its abundance should be similar, for example, to the γ-subunit, but there is no such unidentified band of similar abundance in silver-stained gels run on the immunoprecipitated $F_1F_o$ used in the present experiments (Fig S1).

**(B) Biophysical evidence ruling out contamination of different $K^+$ channels separate and distinct from $F_1F_o$.** Currents were recorded in $F_1F_o$-reconstituted bilayers in the voltage-ramp mode. Five random examples of the voltage-ramp data are given (depicted in different colors). Inset shows the current noise amplitude around $E_{rev}$ with reconstituted $F_1F_o$ which remains identical to that without reconstituted proteins. In the hypothetical scenario of the presence of a separate $K^+$ channel distinct from $F_1F_o$, the noise amplitude at $E_{rev}$ created by the continuous, independent channel-gating activities of the two putative independent channels (whose time-averaged currents exact summate to zero at $E_{rev}$) would be significantly larger than pure bilayer noise, and distinct from $F_1F_o$. In contrast, if ions entering and turning the $F_1F_o$ c-ring are sharing the common path for both $K^+$ and $H^+$ permeation, independent gating would not occur - there would be no channel-gating activity at all at $E_{rev}$ – therefore, the noise amplitude at $E_{rev}$ would be equivalent to that of the plain bilayer without reconstituted proteins. The absence of any channel-gating at $E_{rev}$ (Fig inset) rules out the possibility that $K^+$ is being conducted by a separate $K^+$ channel co-immunoprecipitated with $F_1F_o$. **(C) Immunocaptured $F_1F_o$ is devoid of contamination with ATP-dependent ROMK potassium channel.** Immunoblot analysis of rat heart mitochondria (10 µg; lane 1), left ventricle tissue lysate (10 µg; lane 2) and immunocapture purified rat heart $F_1F_o$ (~5 µg) separated on a NuPage 4-12% Bis-Tris gel prior to immunoblot assay and tested with anti-KCNJ1 (Sigma Prestige, anti-KCNJ1 Cat#:HPA026962; this antibody has been employed to identify mitochondrial ROMK channel by (Foster et al., 2012)), the blot was then re-probed with anti-ATP synthase subunit α antibody. Arrows point to location of the ROMK channel and the ATP synthase subunit α. While left ventricle and cardiac mitochondria extracts showed the expected weak but positive band ~45 kDa, there was no immunolabelling whatsoever in our $F_1F_o$ complex. Thus there is no ROMK contaminating our preparations to yield any functional artefact.

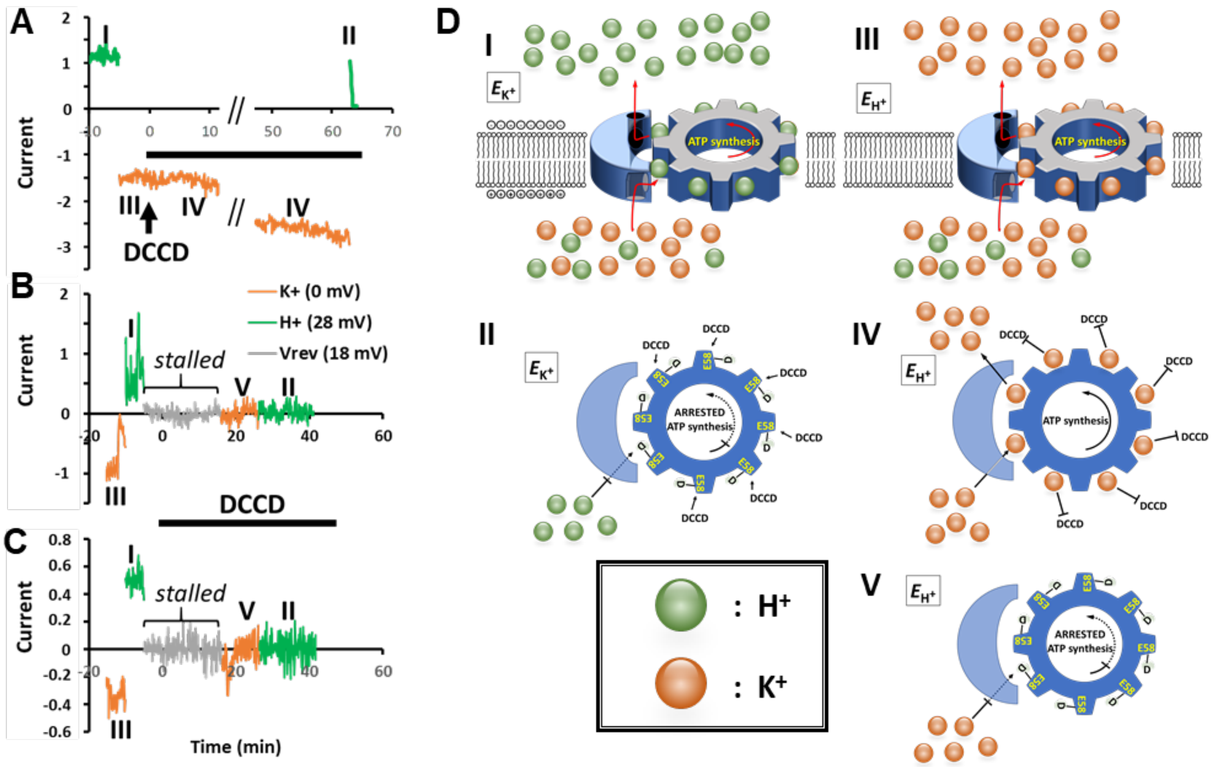


**Figure S3 – related to Figure 2 F-H. Regulation of $F_1F_o$ unitary currents by DCCD (sequel).** (**A**) Continuous recording of $H^+$ unitary current (green; mode I; normalized), followed by $K^+$-only current (orange; mode III; 0 mV; normalized), $K^+$ ions block (mode IV) the DCCD access to the essential Glu58 on the c-ring, allowing the ATP synthesis to continue; the protection of Glu58 is stable for an extend time frame (> 60 min). Switching to the proton mode in the presence of DCCD (II; 28 mV) results in an immediate inhibition of the $H^+$ current and an arrest of the ATP synthesis. (**B-C**) Two examples of the DCCD inhibition of the protonated $F_o$. $K^+$ current (III; 0 mV; normalized) followed by the switch to $H^+$ mode (I; 28 mV) and then to stalled (18mV); the DCCD application during stalled mode (in the presence of $H^+$) resulted in a complete inhibition of the current and an arrest of the ATP synthesis.

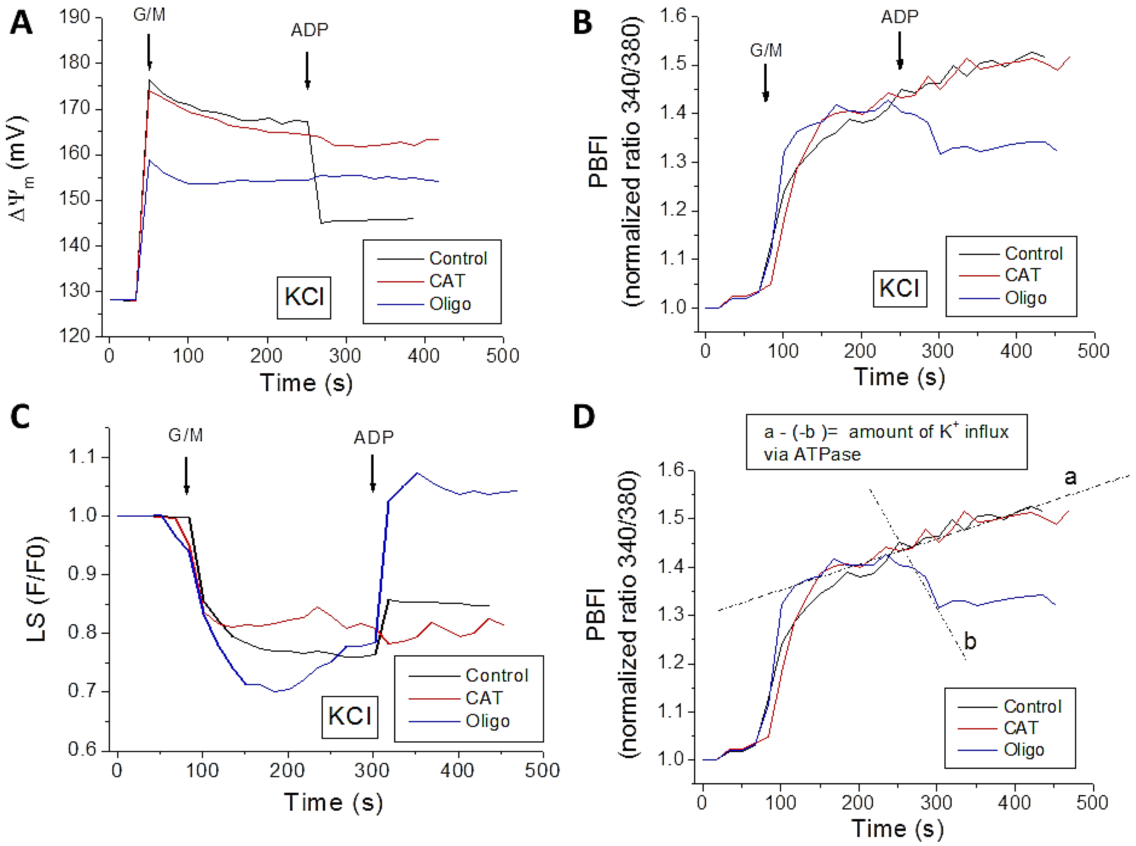


**Figure S4 A-D, related to Figure 3G-O. $K^+$ fluxes during the state 4 → 3 transition in isolated mitochondria from guinea pig heart.** Freshly isolated mitochondria loaded with 20 µM PBFI-AM as described before (Aon et al., 2010) were preincubated in the absence (Control) or presence of 10 µM oligomycin (Oligo) or 10 µM carboxyatractyloside (CAT; ADP/ATP translocase inhibitor) (in the cuvette of the spectrofluorimeter, with stirring at 37ºC for 3min) followed by energization with the $Na^+$ salts of substrates glutamate/malate (G/M, 5 mM/5 mM) (state 4 respiration) and, subsequently, with 1mM ADP (state 3 respiration), added as indicated with arrows. Mitochondria were assayed in the same medium above described containing 137 mM KCl (instead of sucrose) and 2 mM Pi. Mitochondrial $\Delta\Psi_m$ (**A**), PBFI (**B**) and swelling (light scattering, LS) (**C**), were monitored simultaneously as described above. (C, D) Volume changes during mitochondrial swelling-contraction, and of the $K^+$ flux sustained by the ATP synthase in state 3 respiration, were estimated in control and oligomycin-preincubated mitochondria. The relative volume changes (measured as LS) were: *swelling* triggered by G/M (~ 0.3 for Oligo and ~0.2 for control and carboxyatractyloside, CAT), and *contraction* by ADP (~ 0.1 for control and ~ 0.3 for Oligo) additions (**C**). On these bases, the volume change quantification can be estimated considering that 1.77 µl/mg mitochondrial protein (see (Aon et al., 2010) p.75, 2nd column, 2nd paragraph) correspond to the volume in state 4 respiration after G/M addition. A Δvolume of 30% contraction after ADP addition can be estimated equivalent to 0.5 µl/mg mito prot., thus from 1.77 to 1.24 µl/mg mito protein, sustained by a $K^+$ flux equivalent to 60% of the $K^+$ flux at $V_{max}$.

(D) Estimation of the $K^+$ flux sustained by the ATP synthase in state 3 respiration in the presence of Oligo: In KCl, $V_{max}$ of $K^+$ uptake rate = 172±17 nmol $K^+$/min/mg prot. (see (Aon et al., 2010), their Fig 2A).

Slopes (PBFI ratio/min) = Control: 0.00102; +Oligo: - 0.00169

$dK^+/dt$ = influx – efflux = flux balance from control.

Flux balance from control + efflux = influx

0.00102 + 0.00169 = 0.00271 PBFI ratio/min = 0.0183 PBFI ratio/min/mg prot. (148 µg mitochondrial prot. per assay)

$K^+$ flux = 0.0183 * 1000/0.175 = 104 nmol $K^+$/min/mg prot.

For a $V_{max}$ uptake rate of $K^+$ = 172 nmol/min/mg prot.; 104 nmol $K^+$/min/mg prot. represents 60% of the total $K^+$ flux at $V_{max}$.

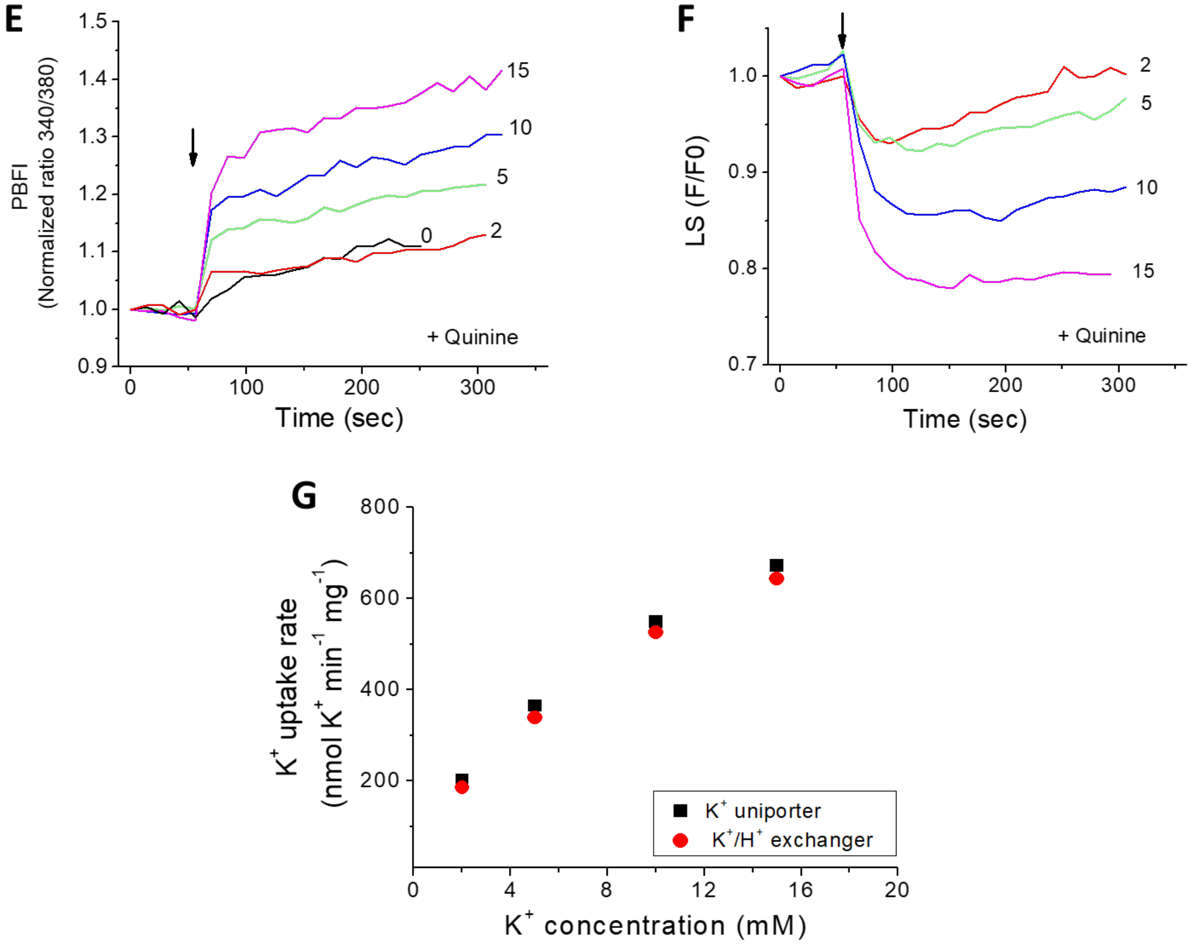


**Figure S4 E-G, related to Figure 3G-O. $K^+$ uniporter and $K^+/H^+$ exchanger fluxes in the absence or presence of quinine in isolated mitochondria from guinea pig heart.**

Freshly isolated PBFI-loaded mitochondria were assayed for $K^+$ fluxes in the same medium described in the legend of Fig S4 A-D, but containing 250 mM sucrose (instead of KCl), thus enabling us to study $K^+$ fluxes in a controlled fashion. After pulses of 2 to 15 mM $KH_2PO_4$ (indicated by arrow), $K^+$ fluxes were quantified in mitochondria energized with the $Na^+$ salts of G/M (5 mM/5 mM), and in the absence (Control) or presence of 50 µM quinine, a $K^+/H^+$ exchanger inhibitor (Garlid et al., 1986; Martin et al., 1986). Mitochondrial swelling (LS: **F**) and PBFI fluorescence (**E**) were determined in control and quinine-preincubated (**E, F**) mitochondria. (**G)** depicts the $K^+$ fluxes sustained by the $K^+$ uniporter and the $K^+/H^+$ exchanger as measured in the presence of quinine or its absence, respectively. The flux through the $K^+/H^+$ exchanger was calculated as the difference between the fluxes measured in the presence of quinine (uniporter) minus in its absence (where both uniporter and exchanger are active).

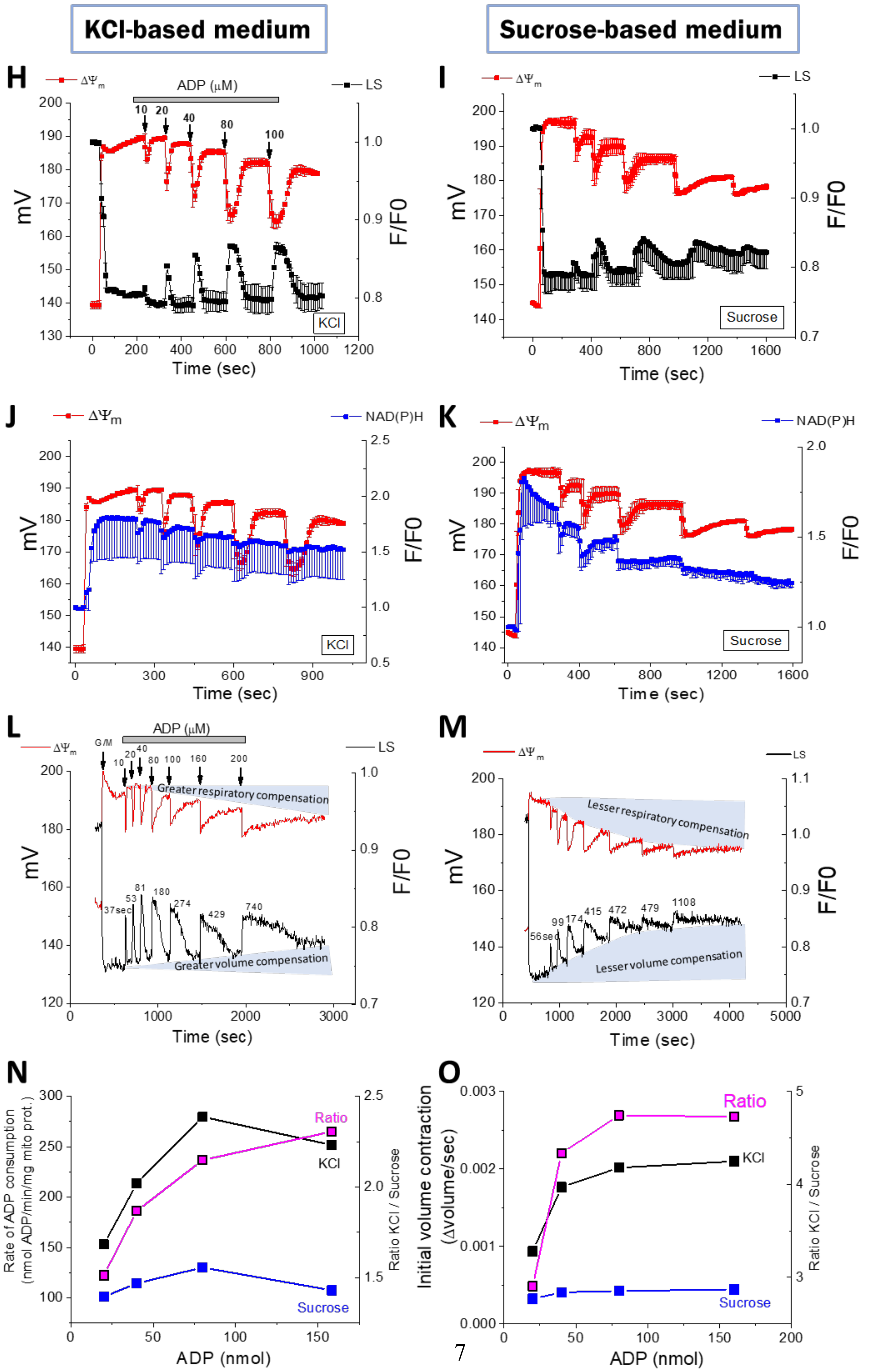
KCl-based medium
Sucrose-based medium
H
ADP (μM)
10 20 40 80 100
ΔΨm
LS
mV
F/F0
Time (sec)
KCl
I
Sucrose
J
NAD(P)H
K
L
G/M
Greater respiratory compensation
Greater volume compensation
37sec 53 81 180 274 429 740
M
Lesser respiratory compensation
Lesser volume compensation
56sec 99 174 415 472 479 1108
N
Rate of ADP consumption (nmol ADP/min/mg mito prot.)
Ratio KCl / Sucrose
ADP (nmol)
Ratio
Sucrose
O
Initial volume contraction (Δvolume/sec)

**Figure S4 H-O, related to Figure 3G-O. Mitochondrial response to sequential additions of ADP in KCl- and sucrose-based media.** (**H-M**) Multiple ADP additions, as indicated, lead to successive cycles of shrinkage and swelling as judged by 90º light scattering (LS, black lines) and concomitant changes in $\Delta\Psi_m$ (TMRM fluorescence, red lines) and NAD(P)H (blue lines) monitored with a PTI QuantaMaster spectrofluorometer (Photon Technology International Inc.). (**H, J, L**) KCl-based medium, and (**I, K, M**) sucrose medium at matched (260 mOsm) osmolality. **(H-K**) correspond to measurements performed with the same mitochondrial preparations utilized for the measurements of mitochondrial respiratory and energetic performance shown in Figure 3G-O; error bars were obtained from 3 independent mitochondrial preparations from 3 different hearts.

(**L-M)** correspond to a representative experiment where it is shown that mitochondria in $K^+$-based medium display larger amplitude excursions and higher frequency responses (as given by the numbers close to the LS traces, which correspond to the length of the condensed-orthodox cycle in seconds, i.e., state 4-to-3 transitions), after consecutive ADP additions, reflecting better adaptation to changes in mitochondrial energy supply and a more efficient energy generation. The shaded background schematically highlight the poorer respiratory compensation and increased degree of uncompensated volume contraction in the sucrose-based media *vs.* $K^+$-based media.

**(N)** and (**O**), derived from **L** and **M**, display initial rates of ADP phosphorylation ~2.3 times faster in KCl than in sucrose (see KCl/sucrose ratio in G, magenta line), and initial rates of volume contraction ~4.7 times larger in KCl than in sucrose (see ratio in (**H)**, magenta line), underlying the better efficiency of ATP synthase in the presence of $K^+$.

**Supplemental Table S1, related to Figure 3G. Comparison of Oligomycin-sensitive Oxygen Consumption Rates (OCR) from rat heart isolated mitochondria**

| Rat heart mitochondria (strain, sex, mitochondrial location) | Oligomycin-sensitive OCR (ng at O $min^{-1}$ $mg^{-1}$ prot) | Reference |
|---|---|---|
| Sprague Dawley; male | 122 | This work |
| Sprague Dawley; male | 250 | (Pelikan et al., 1987) |
| Wistar; female | 100 | (Ribeiro et al., 2017) |
| Wistar; male; intermyofibrillar | 250 | (Ribeiro et al., 2016) |
| Wistar; female; intermyofibrillar | 110 | (Ribeiro et al., 2016) |
| Wistar; male; subsarcolemmal | 150 | (Ribeiro et al., 2016) |
| Wistar; female; subsarcolemmal | 110 | (Ribeiro et al., 2016) |
| Sprague Dawley; female | 200 | (Vanasco et al., 2014) |
| Sprague Dawley; female | 130 | (Vanasco et al., 2012) |
| Sprague Dawley; male | 250 | (Makazan et al., 2009) |

For comparison, the literature was surveyed using the following criteria: a) similarity in the incubation medium, i.e., presence of KCl, absence of $Ca^{2+}$, pH, substrates, temperature; b) compatibility of units of measurement; c) accounted for oligo-sensitive respiration (i.e., in all cases state 4 respiration was subtracted from state 3 respiration to compare with our data of OCR oligo-sensitive)

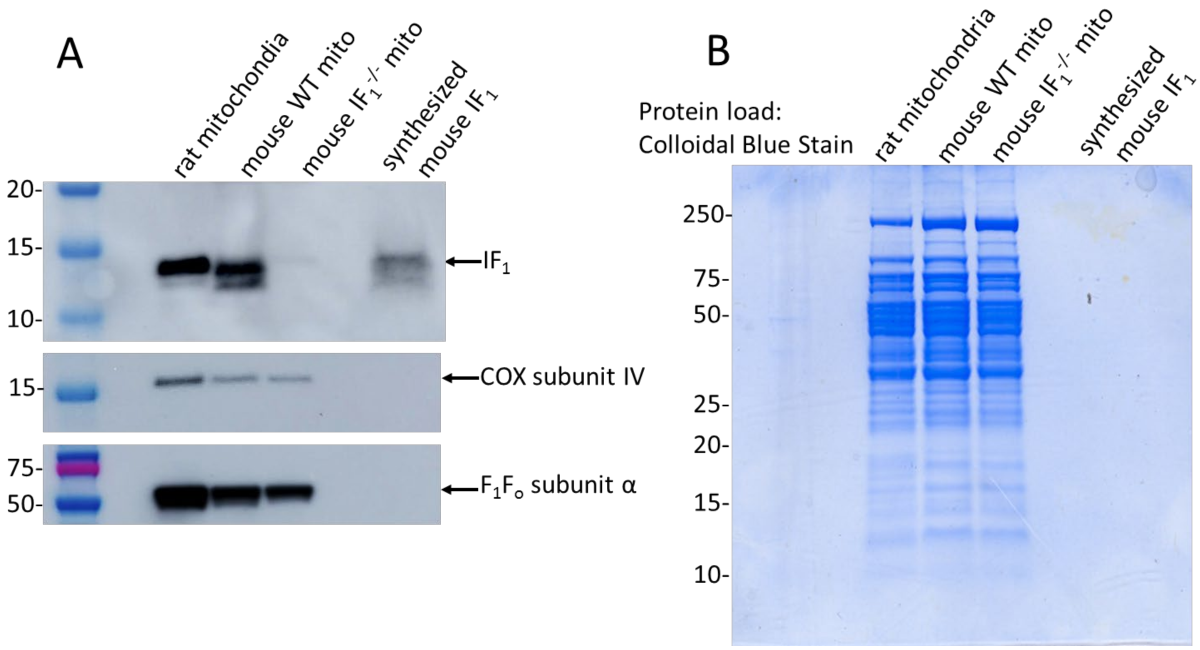


**Figure S5-related to Figure 5D. Phenotype verification of $IF_1$ KO mouse**. (**A**) $IF_1$ protein expression was not detected by an anti-$IF_1$ antibody (Cell Signaling Technology) in $IF_1^{-/-}$ mouse mitochondria by western blotting while the blot was positive for $IF_1$ protein expresion in rat and mouse wild type (WT) mitochondria (15 µg of proteins were loaded per well) and synthetized mouse $IF_1$ peptide (purified >98%; 50 ng). As expected, all three mitochondrial preparations showed the presence of other mitochondrial proteins (the same blot was reprobed for COX subunit IV (Cell Signaling Technology, clone 3E11) and $F_1F_o$ subunit alpha (Abcam). (**B**) The gel was stained with Colloidal Blue Stain (Invitrogen) to demonstrate the equal total mitochondrial protein load into each well.

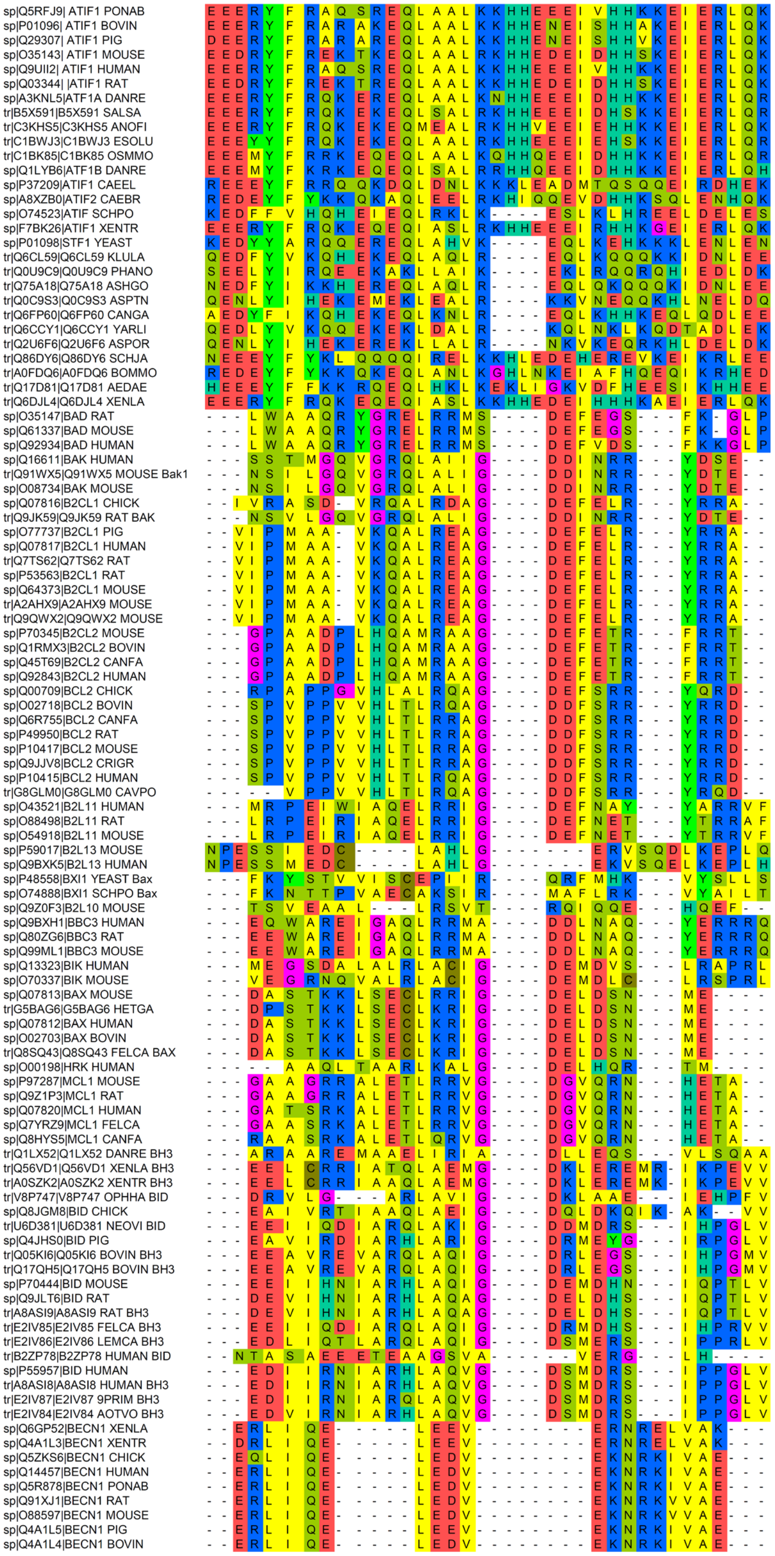


**Figure S6, related to Figure 5A**. **Multiple sequence alignment of BH3 motifs of Bcl-2 proteins and IF1.** The alignment was obtained with Clustal Omega. Details of the 107 sequences corresponding to fragments of the IF1 (28 correspond to IF1 sequences) and Bcl-2 family proteins are provided in Table S2.

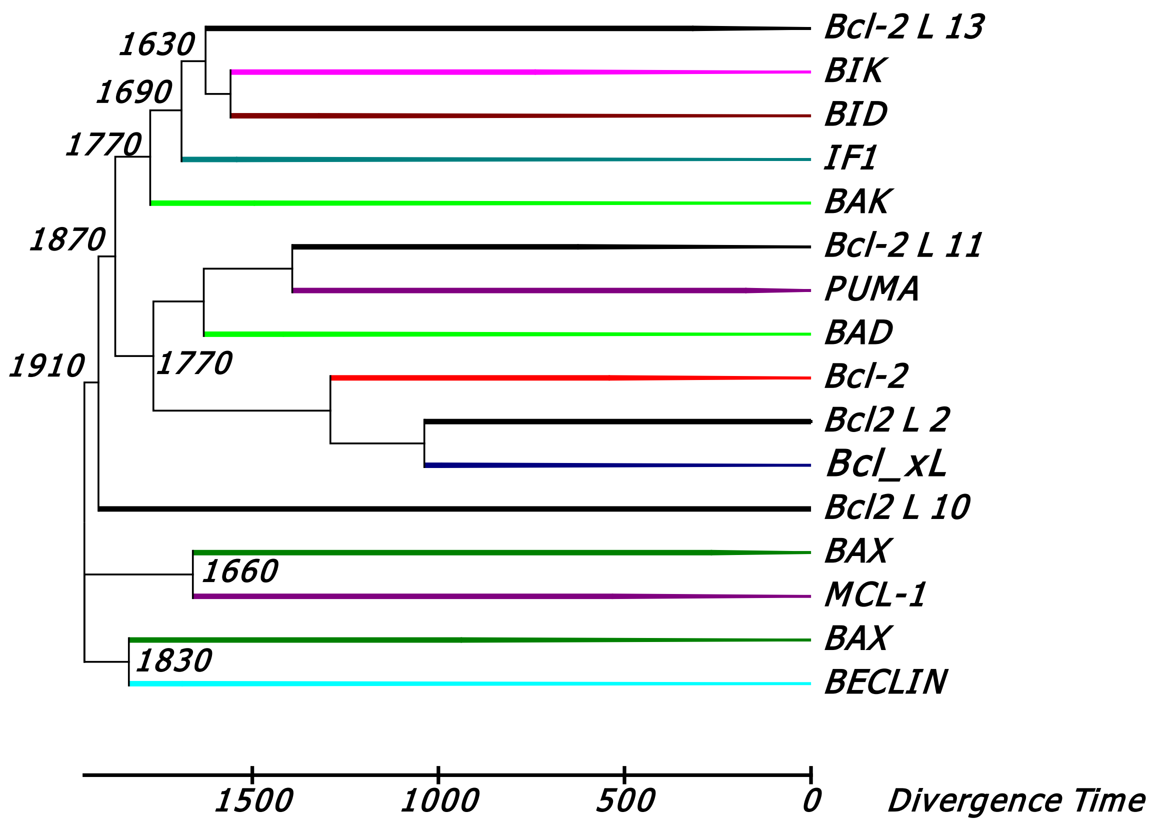


**Figure S7 A, related to Figure 7. Phylogenetic tree of IF1 and BH3 motif containing proteins.**
The tree was calculated with the Neighbor-Joining statistical method (Saitou and Nei, 1987) from the alignment performed by Clustal Omega (Fig S6). The calibration of the divergence times (in millions of years, Myr) was performed as described in Methods and to calculate the branching times the method RelTime (Tamura et al., 2012) was applied. Only positions with more than 95% site coverage were considered for the analysis, which in this case corresponds to 14 amino acids. The tree construction was performed with MEGA 6.0 (Tamura et al., 2013)

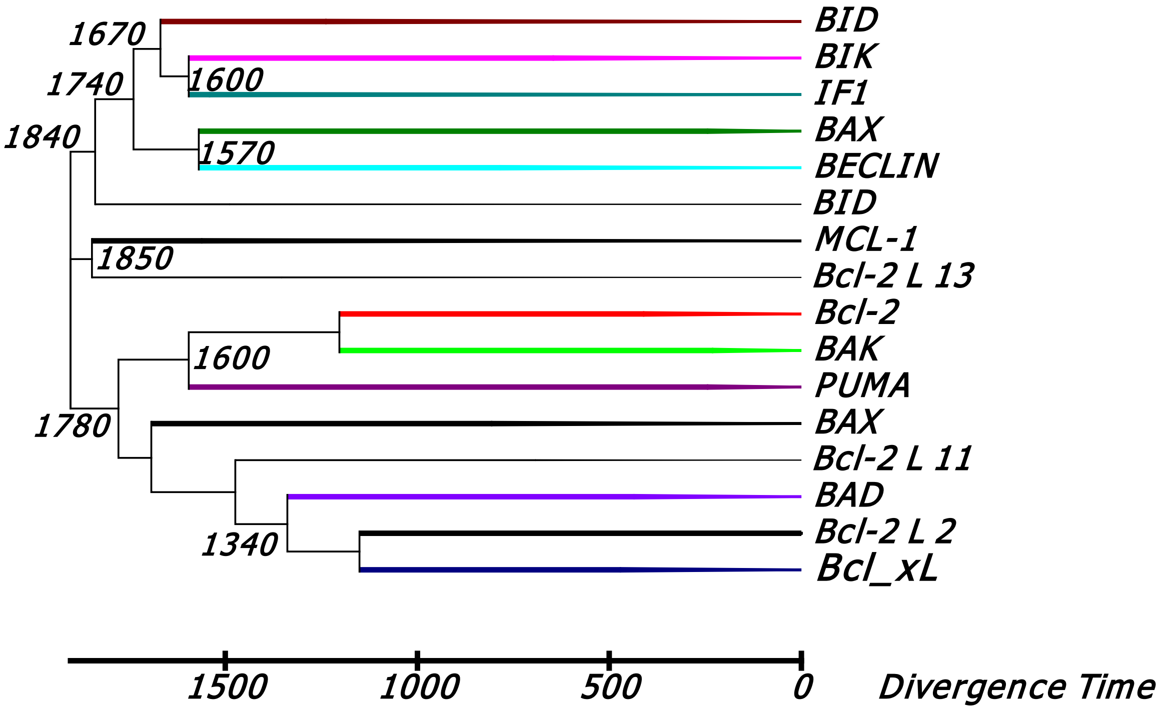


**Figure S7 B, related to Figure 7. Phylogenetic tree of IF1 and Bcl-2 protein BH3 motif containing fragments.** The tree was calculated with the Neighbor-Joining statistical method from the alignment performed by Bali-Phy with a LG substitution model (Le and Gascuel, 2008) and the Insertion/deletion model RS07 (Redelings and Suchard, 2007). The calibration of the divergence times was performed as described in methods and to calculate the branching times the method RelTime was applied. Only positions with more than 95% site coverage were considered for the analysis, which in this case corresponds to 9 residues. The tree construction was performed with MEGA 6.0. The bootstrap consensus tree obtained in Mega 6.0 was very similar to that calculated by the joint estimation of Bali-phy.

**Supplemental Table S2, related to Figure 7E**

Key for the input sequences used in the phylogenetic analysis

| | | | |
|---|---|---|---|
| sp\|Q1LYB6\|ATF1B | DANRE | ATIF1 | *Danio rerio* |
| sp\|A3KNL5\|ATF1A | DANRE | ATIF1 | *Danio rerio* |
| sp\|P01096\|ATIF1 | BOVIN | ATIF1 | *Bos Taurus* |
| sp\|P37209\|ATIF1 | CAEEL | ATIF1 | *Caenorhabditis elegans* |
| sp\|Q9UII2\|ATIF1 | HUMAN | ATIF1 | *Homo sapiens* |
| sp\|Q5RFJ9\|ATIF1 | PONAB | ATIF1 | *Pongo abelii* |
| sp\|A8XZB0\|ATIF2 | CAEBR | ATIF1 | *Caenorhabditis briggsae* |
| sp\|O74523\|ATIF | SCHPO | ATIF1 | *Schizosaccharomyces pombe* |
| sp\|Q03344\|ATIF1 | RAT | ATIF1 | *Rattus norvegicus* |
| sp\|F7BK26\|ATIF1 | XENTR | ATIF1 | *Xenopus tropicalis* |
| sp\|Q29307\|ATIF1 | PIG | ATIF1 | *Sus scrofa* |
| sp\|O35143\|ATIF1 | MOUSE | ATIF1 | *Mus musculus* |
| sp\|P01098\|STF1 | YEAST | Putative ATPase inhibitor | *Saccharomyces cerevisiae* |
| tr\|Q6CL59\|Q6CL59 | KLULA | Putative ATPase inhibitor | *Kluyveromyces lactis* |
| tr\|Q0U9C9\|Q0U9C9 | PHANO | Putative ATPase inhibitor | *Phaeosphaeria nodorum* |
| tr\|Q75A18\|Q75A18 | ASHGO | Putative ATPase inhibitor | *Ashbya gossypii* |
| tr\|Q0C9S3\|Q0C9S3 | ASPTN | Putative ATPase inhibitor | *Aspergillus terreus* |
| tr\|Q6FP60\|Q6FP60 | CANGA | Putative ATPase inhibitor | *Candida glabrata* |
| tr\|Q6CCY1\|Q6CCY1 | YARLI | Putative ATPase inhibitor | *Yarrowia lipolytica* |
| tr\|Q2U6F6\|Q2U6F6 | ASPOR | Putative ATPase inhibitor | *Aspergillus oryzae* |
| tr\|Q86DY6\|Q86DY6 | SCHJA | Putative ATPase inhibitor | *Schistosoma japonicum* |
| tr\|A0FDQ6\|A0FDQ6 | BOMMO | ATIF1 | *Bombyx mori* |
| tr\|Q17D81\|Q17D81 | AEDAE | Putative ATPase inhibitor | *Aedes aegypti* |
| tr\|B5X591\|B5X591 | SALSA | ATIF1 | *Salmo salar* |
| tr\|C1BWJ3\|C1BWJ3 | ESOLU | ATIF1 | *Esox Lucius* |
| tr\|C3KHS5\|C3KHS5 | ANOFI | ATIF1 | *Anoplopoma fimbria* |
| tr\|C1BK85\|C1BK85 | OSMMO | ATIF1 | *Osmerus mordax* |
| tr\|Q6DJL4\|Q6DJL4 | XENLA | atpif1 | *Xenopus laevis* |
| sp\|Q16611\|BAK | HUMAN | BAK1 | *Homo sapiens* |
| tr\|Q91WX5\|Q91WX5 | MOUSE | Bak1 | *Mus musculus* |
| sp\|O08734\|BAK | MOUSE | Bak1 | *Mus musculus* |
| sp\|Q07816\|B2CL1 | CHICK | BCL2L1 | *Gallus gallus* |
| tr\|Q9JK59\|Q9JK59 | RAT | Bak1 | *Rattus norvegicus* |
| sp\|O77737\|B2CL1 | PIG | BCL2L1 | *Sus scrofa* |
| sp\|Q07817\|B2CL1 | HUMAN | BCL2L1 | *Homo sapiens* |
| tr\|Q7TS62\|Q7TS62 | RAT | Bcl2l1 | *Rattus norvegicus* |
| sp\|P53563\|B2CL1 | RAT | Bcl2l1 | *Rattus norvegicus* |
| sp\|Q64373\|B2CL1 | MOUSE | Bcl2l1 | *Mus musculus* |
| tr\|A2AHX9\|A2AHX9 | MOUSE | Bcl2l1 | *Mus musculus* |
| tr\|Q9QWX2\|Q9QWX2 | MOUSE | Bcl2l1 | *Mus musculus* |

| | | | |
|---|---|---|---|
| sp\|P70345\|B2CL2 | MOUSE | Bcl2l2 | *Mus musculus* |
| sp\|Q1RMX3\|B2CL2 | BOVIN | BCL2L2 | *Bos Taurus* |
| sp\|Q45T69\|B2CL2 | CANFA | BCL2L2 | *Canis familiaris* |
| sp\|Q92843\|B2CL2 | HUMAN | BCL2L2 | *Homo sapiens* |
| sp\|Q00709\|BCL2 | CHICK | BCL2 | *Gallus gallus* |
| sp\|O02718\|BCL2 | BOVIN | BCL2 | *Bos Taurus* |
| sp\|Q6R755\|BCL2 | CANFA | BCL2 | *Canis familiaris* |
| sp\|P49950\|BCL2 | RAT | Bcl2 | *Rattus norvegicus* |
| sp\|P10417\|BCL2 | MOUSE | Bcl2 | *Mus musculus* |
| sp\|Q9JJV8\|BCL2 | CRIGR | BCL2 | *Cricetulus griseus* |
| sp\|P10415\|BCL2 | HUMAN | BCL2 | *Homo sapiens* |
| tr\|G8GLM0\|G8GLM0 | CAVPO | BCL2 | *Cavia porcellus* |
| sp\|O43521\|B2L11 | HUMAN | BCL2L11 | *Homo sapiens* |
| sp\|O88498\|B2L11 | RAT | Bcl2l11 | *Rattus norvegicus* |
| sp\|O54918\|B2L11 | MOUSE | Bcl2l11 | *Mus musculus* |
| sp\|P59017\|B2L13 | MOUSE | Bcl2l13 | *Mus musculus* |
| sp\|Q9BXK5\|B2L13 | HUMAN | BCL2L13 | *Homo sapiens* |
| sp\|Q92934\|BAD | HUMAN | BAD | *Homo sapiens* |
| sp\|Q61337\|BAD | MOUSE | Bad | *Mus musculus* |
| sp\|O35147\|BAD | RAT | Bad | *Rattus norvegicus* |
| sp\|P48558\|BXI1 | YEAST | Bax | *Saccharomyces cerevisiae* |
| sp\|O74888\|BXI1 | SCHPO | Bax | *Schizosaccharomyces pombe* |
| sp\|Q9Z0F3\|B2L10 | MOUSE | Bcl2l10 | *Mus musculus* |
| sp\|Q9BXH1\|BBC3 | HUMAN | BBC3 | *Homo sapiens* |
| sp\|Q80ZG6\|BBC3 | RAT | Bbc3 | *Rattus norvegicus* |
| sp\|Q99ML1\|BBC3 | MOUSE | Bbc3 | *Mus musculus* |
| sp\|Q13323\|BIK | HUMAN | BIK | *Homo sapiens* |
| sp\|O70337\|BIK | MOUSE | Bik | *Mus musculus* |
| sp\|Q07813\|BAX | MOUSE | BAX | *Mus musculus* |
| tr\|G5BAG6\|G5BAG6 | HETGA | BAX | *Heterocephalus glaber* |
| sp\|Q07812\|BAX | HUMAN | BAX | *Homo sapiens* |
| sp\|O02703\|BAX | BOVIN | BAX | *Bos Taurus* |
| tr\|Q8SQ43\|Q8SQ43 | FELCA | BAX | *Felis catus* |
| sp\|O00198\|HRK | HUMAN | HRK | *Homo sapiens* |
| sp\|P97287\|MCL1 | MOUSE | Mcl1 | *Mus musculus* |
| sp\|Q9Z1P3\|MCL1 | RAT | Mcl1 | *Rattus norvegicus* |
| sp\|Q07820\|MCL1 | HUMAN | MCL1 | *Homo sapiens* |
| sp\|Q7YRZ9\|MCL1 | FELCA | MCL1 | *Felis catus* |
| sp\|Q8HYS5\|MCL1 | CANFA | MCL1 | *Canis familiaris* |
| tr\|Q1LX52\|Q1LX52 | DANRE | BID | *Danio rerio* |
| tr\|Q56VD1\|Q56VD1 | XENLA | BID | *Xenopus laevis* |
| tr\|A0SZK2\|A0SZK2 | XENTR | BID | *Xenopus tropicalis* |
| tr\|V8P747\|V8P747 | OPHHA | BID | *Ophiophagus hannah* |
| sp\|Q8JGM8\|BID | CHICK | BID | *Gallus gallus* |
| tr\|U6D381\|U6D381 | NEOVI | BID | *Neovison vison* |

| | | | |
|---|---|---|---|
| sp\|Q4JHS0\|BID | PIG | BID | *Sus scrofa* |
| tr\|Q05KI6\|Q05KI6 | BOVIN | BID | *Bos Taurus* |
| tr\|Q17QH5\|Q17QH5 | BOVIN | BID | *Bos Taurus* |
| sp\|P70444\|BID | MOUSE | BID | *Mus musculus* |
| sp\|Q9JLT6\|BID | RAT | BID | *Rattus norvegicus* |
| tr\|A8ASI9\|A8ASI9 | RAT | BID | *Rattus norvegicus* |
| tr\|E2IV85\|E2IV85 | FELCA | BID | *Felis catus* |
| tr\|E2IV86\|E2IV86 | LEMCA | BID | *Lemur catta* |
| tr\|B2ZP78\|B2ZP78 | HUMAN | BID | *Homo sapiens* |
| sp\|P55957\|BID | HUMAN | BID | *Homo sapiens* |
| tr\|A8ASI8\|A8ASI8 | HUMAN | BID | *Homo sapiens* |
| tr\|E2IV87\|E2IV87 | 9PRIM | BID | *Saimiri boliviensis* |
| tr\|E2IV84\|E2IV84 | AOTVO | BID | *Aotus vociferans* |
| sp\|Q6GP52\|BECN1 | XENLA | Beclin-1 | *Xenopus laevis* |
| sp\|Q4A1L3\|BECN1 | XENTR | Beclin-1 | *Xenopus tropicalis* |
| sp\|Q5ZKS6\|BECN1 | CHICK | Beclin-1 | *Gallus gallus* |
| sp\|Q14457\|BECN1 | HUMAN | Beclin-1 | *Homo sapiens* |
| sp\|Q5R878\|BECN1 | PONAB | Beclin-1 | *Pongo abelii* |
| sp\|Q91XJ1\|BECN1 | RAT | Beclin-1 | *Rattus norvegicus* |
| sp\|O88597\|BECN1 | MOUSE | Beclin-1 | *Mus musculus* |
| sp\|Q4A1L5\|BECN1 | PIG | Beclin-1 | *Sus scrofa* |
| sp\|Q4A1L4\|BECN1 | BOVIN | Beclin-1 | *Bos Taurus* |

### Supplemental Experimental and Analytical Procedures

#### Purification and reconstitution of $F_1F_o$-ATP synthase

All chemicals where from Sigma-Aldrich (St. Louis, MO), except where otherwise noted. Rat and mouse heart mitochondria were solubilized in 1% lauryl maltoside (LM; Anatrace Inc., Maumee, OH) or 1.5% - 2% digitonin for 10 min, unsolubilized material was removed by centrifugation at 25000 x g for 15 min. $F_1F_o$ was immunocaptured using a Complex V immunocapture kit (MitoSciences Inc., Eugene, OR). Purified $F_1F_o$ was reconstituted into proteoliposomes using a modified procedure based on a method developed originally by (Paucek et al., 1992). Briefly, for ATP measurements, cardiolipin and phosphatidylcholine in a 1:9 ratio were dried under nitrogen and dispersed at 100 mg/ml in an outwardly-directed potassium gradient was employed with an internal buffer containing (in mM): 10 $NaH_2PO_4$, 200 $K_2SO_4$ (or KCl; and $Na_2SO_4$ or NaCl), 0.25 EDTA, 20 HEPES, pH 7.0. Lipids were solubilized by octylpentaoxyethylene detergent. ~ 5 µg of isolated $F_1F_o$ was added to 10 mg of solubilized lipids. Proteoliposome (PL) formation was induced by removal of the detergent in Bio-beads SM2 column (Bio-Rad Laboratories, Hercules, CA).

#### Protein detection

BCA (bicinchoninic acid) protein assay (Pierce Biotechnology, Inc., Rockford, IL) was used for measurements of protein concentration.

Integrity and purity of the isolated $F_1F_o$ was determined by gel electrophoresis under native and denaturing conditions. NativePage Novex 3-12% Bis-Tris gel system (Invitrogen) was applied to analyze samples under native clear and native blue (0.002% Coomassie Blue G-250) conditions. Gels were silver or Coomassie stained, or blotted to a PVDF membrane (GE Healthcare Bio-Sciences, Marlborough, MA) for protein immunodetection.

For analysis under denaturing conditions, mitochondria were lysed for 30 min on ice with RIPA buffer (0.15 mol/L NaCl, 10 mmol/L Tris (pH 7.4), 1 % NP-40, 0.1 % SDS, 0.5 % deoxycholate,1 mmol/L NaF, 1 mmol/L sodium orthovanadate) and protease inhibitors (Roche Diagnostics), and the lysates were centrifuged (25 min at 4°C; 25,000 x g). Lysates or purified $F_1F_o$ were heated (10 min at 75 ºC) in the NuPage LDS sample buffer under reducing conditions. Proteins extracts were separated using pre-cast NuPAGE Bis-Tris 4-12% gels (Invitrogen Corp. Carlsbad, CA). Gels were then silver stained or the proteins were transferred to a PVDF membrane. The membranes were blocked with 5% nonfat dry milk in tris(hydroxymethyl)aminomethane (Tris)-buffered saline with 0.1% Tween 20 (TBST) and incubated overnight with primary antibodies against individual Complex V subunits, $IF_1$ (MitoSciences), KCNJ1 Sigma Prestige (Sigma-Aldrich) and ANT (Santa Cruz Biotechnology, Inc., Santa Cruz, CA), followed by horseradish peroxidase-linked secondary antibodies (Amersham Biosciences, Piscataway, NJ). Protein bands were visualized using ECL system (Amersham Biosciences) diluted in TBST.

#### In-gel ATPase assay

The in-gel ATPase assay described in (Wittig et al., 2006) was slightly modified: Clear native (CN) and blue native (BN)-gels were preincubated for 2 hr in 270 mM glycine, 35 mM Tris, pH 8.4. In a parallel assay ATPase was inhibited with 5 µg/ml oligomycin. Gels were incubated for 45 min in ATPase reaction medium (in mM): 270 glycine, 35 Tris, 8 ATP, 14 $MgSO_4$, 0.2% $PbNO_3$, pH 8.4. Gels were washed with water and precipitated white lead phosphate bands were transformed to dark brown lead sulfide bands by immersion of the gels in 5% ammonium sulfide followed by a water rinse. Gels were then fixed (40% methanol, 10 % acetic acid) and protein visualized using a colloidal blue staining kit (Invitrogen).

#### Cell and proteoliposome treatment

The following compounds were applied (alone or in combination) to cells and PL during experimental measurements: 2-4 µM $F_o$-inhibitor venturicidin B (VENT; A.G. Scientific, Inc., San Diego, CA). MgATP (at various concentrations as indicated); 10 nM Tyr-D-Ala-Gly-Phe-D-Leu (DADLE); 100 µM glibenclamide; 30 µM diazoxide (Dz); 50 µM pinacidil; 500 µM 5-hydroxydecanoate (5HD); 1 µM oligomycin, 0.4-0.6 mM DCCD, 1 µM valinomycin; 1 µM FCCP; 20-100 nM Bcl-2, Bcl-xl and Mcl-1 (Sigma-Aldrich, St. Louis, MO); 3 mM LiCl; 2 µM Bcl-2 binding peptide (Bad BH3 peptide) and its negative control (Calbiochem Corp., San Diego, CA).

#### Quantitative comparison of $H^+$ and $K^+$ current magnitudes through ATP synthase

To compute the ratio of $H^+$ and $K^+$ currents through the ATP synthase we consider the contribution of each ion as described by the Goldman-Hodgkin-Katz formalism:

$$\frac{I_{H^+}}{I_{K^+}}=\frac{P_H {Z_H}^2\frac{\Delta\Psi_m F^2}{RT}\left(\frac{[H]_i-[H]_0\exp\left(\frac{-Z_H\Delta\Psi_m F}{RT}\right)}{1-\exp\left(\frac{-Z_H\Delta\Psi_m F}{RT}\right)}\right)}{P_K {Z_K}^2\frac{\Delta\Psi_m F^2}{RT}\left(\frac{[K]_i-[K]_0\exp\left(\frac{-Z_K\Delta\Psi_m F}{RT}\right)}{1-\exp\left(\frac{-Z_K\Delta\Psi_m F}{RT}\right)}\right)}$$

Taking the limit of the ratio of currents when $\Delta\Psi_m$ tends to - ∞, i.e., in the direction of ATP synthesis,

$$\lim_{\Delta\Psi_m\to-\infty}\left(\frac{I_{H^+}}{I_{K^+}}\right)=\frac{P_H {Z_H}^2\left(\frac{[H]_i-[H]_0\exp\left(\frac{Z_H\infty F}{RT}\right)}{1-\exp\left(\frac{Z_H\infty F}{RT}\right)}\right)}{P_K {Z_K}^2\left(\frac{[K]_i-[K]_0\exp\left(\frac{Z_K\infty F}{RT}\right)}{1-\exp\left(\frac{Z_K\infty F}{RT}\right)}\right)}$$

the linear terms ([H]i, [K]i and the 1 in the numerators and denominators, respectively), become negligible with respect to the exponential terms containing the dependence on $\Delta\Psi_m$ as follows:

$$\lim_{\Delta\Psi_m\to-\infty}\left(\frac{I_{H^+}}{I_{K^+}}\right)=\frac{P_H {Z_H}^2\left(\frac{\cancel{[H]_i}-[H]_0\exp\left(\frac{Z_H\infty F}{RT}\right)}{\cancel{1}-\exp\left(\frac{Z_H\infty F}{RT}\right)}\right)}{P_K {Z_K}^2\left(\frac{\cancel{[K]_i}-[K]_0\exp\left(\frac{Z_K\infty F}{RT}\right)}{\cancel{1}-\exp\left(\frac{Z_K\infty F}{RT}\right)}\right)}$$

rendering

$$\lim_{\Delta\Psi_m\to-\infty}\left(\frac{I_{H^+}}{I_{K^+}}\right)=\frac{P_H {Z_H}^2[H^+]_0}{P_K {Z_K}^2[K^+]_0}$$

This limit will be (asymptotically) approached as the magnitude of $\Delta\Psi_m$ increases, but under realistic ionic conditions, is practically achieved for $|\Delta\Psi_m| \geq 100$ mV. The currents ratio through ATP synthase will be proportional to the ratio of extra-mitochondrial concentrations (in mole-equivalents) of $[H^+] = 6.3\times10^{-8}$ over $[K^+] = 0.140$, at pH 7.2, corresponding to ~ $5\times10^{-7}$ : 1. This results in a ratio of $H^+$: $K^+$ currents ~ 1:3.7 since the ratio of permeability $P_H$:$P_K$ is equal to $5.2\pm0.9 \times 10^{-11}$ : $8.7\pm2.9 \times 10^{-17}$ as determined in our experiments.

**Mitochondria and submitochondrial membrane particles isolation**

Mitochondria were isolated by differential centrifugation of the heart homogenate.

Fresh submitochondrial membrane particles (SMP) were made by sonicating rat heart mitochondria until cloudy preparation turns clear, then centrifugation at 16,000 x g for 15 minutes and subsequent centrifugation of the supernatant at 140,000 x g for 20 minutes to pellet the SMPs, which were resuspended in isolation medium. For electrophysiological measurements the isolated SMPs were added directly into the *trans* chamber.

**ATP measurements**

External buffer (EB) contained (in mM): 10 $NaH_2PO_4$, 200 TEA-$SO_4$ (or TEA-Cl), 2.5 $MgCl_2$, 0.090 ADP (ultra-pure; Cell Technology Inc., Mountain View, CA), 1 mM EDTA, 0.001 FCCP, 20 TEA-HEPES pH 7.0, and using a Vapro osmometer (Wescor, Inc., Logan, UT) the osmotic activity of the internal solution ($K^+$=200 mEq) was matched to that of the EB ($K^+$=0, $TEA^+$=200 mEq) with an appropriate amount of mannitol. 5 µl of formed PL was resuspended in 495 µl of EB. After 5 min incubation at room temperature the ATP production was determined by luciferase/luciferin system using ATP bioluminescence assay kit HS II (Roche Diagnostics GmbH, Penzberg, Germany) in a Tropix TR 717 microplate luminometer (Bedford, MA).

To measure ATP production/consumption kinetics we developed a novel on-command acceleration-triggered inertial injector system, contained entirely inside individual wells of a 96 well plate, capable of accurately and reproducibly introducing and admixing 1-2 µL volumes in 1-5 sec (as desired, and without optical interference) during continuous chemiluminescence recording protocols. Specifically, individual wells of a 96 well plate were loaded with 198 µL of EB (see above) supplemented with 30 µM luciferin (Sigma Aldrich) and 30 pmol/well of luciferase (Promega Corp., Madison, WI); a meniscus-stabilized microcarrier containing 1-2 µL of PL (loaded with internal 200 $K^+$ buffer) was carefully placed on top of the EB. After the background-level luminescence was acquired (Victor3 plate reader; PerkinElmer Inc.; Waltham, MA), the ATP generating system (PL) was injected into the buffer-containing well (which initiates the ATP synthesis reaction). This method enables measurement the relatively fast reaction kinetics of ATP synthesis (from microliter reaction quantities) from baseline to completion of the reaction.

**$K^+$ flux measurements**

A similar procedure as described above was used to prepare PLs for determination of $K^+$ fluxes; additionally, vesicles were loaded with the potassium sensitive probe PBFI (Invitrogen Corp., Carlsbad, CA). Extra-vesicular PBFI was removed by passing the vesicles through 2 ml Bio-Spin columns (Bio-Rad Laboratories), which had been pre-equilibrated with internal buffer without the probe. In these experiments, an inwardly-directed potassium gradient was employed and the internal buffer contained (in mM): 125 TEA-$SO_4$, 1 EDTA, 25 TEA-HEPES pH 7.0, and 0.3 PBFI. The osmotic activity of the internal solution ($K^+$=0) was matched to that of the external buffer ($K^+$=150 mEq) with an appropriate amount of mannitol, using an osmometer. 5 µl of suspension was transferred with vigorous stirring into a cuvette containing 495 µl: in mM 150 KCl, 1 EDTA, 25 HEPES, 1 µM FCCP, pH 7.0. PBFI-fluorescence ratio was measured continuously in a Perkin-Elmer LS 50B fluorescence spectrophotometer and the normalized initial rate of rise of the signal was taken as the index of $K^+$ influx.

**Membrane potential measurements**

The potential-sensitive fluorescent probe oxonol VI (20 nM; (Apell and Bersch, 1987; Holoubek et al., 2003) was used to monitor $F_1F_o$-generated $\Delta\psi$ in a $K^+$ gradient. It has been documented that in the presence of an inside-positive membrane potential, this negatively charged dye accumulates into the proteoliposomes according to the Nernst equilibrium. The detailed protocol employed in this study was described in (Holoubek et al., 2003) and the experimental proteoliposomes were prepared as described in the **Supplemental Experimental Procedures** section: **$K^+$ flux measurements**.The lumen of the liposomes was osmotically balanced with the outside medium when different $K^+$ gradients were applied using potassium/choline system (osmolality was measured using a Vapro vapor pressure osmometer). Potential-dependent accumulation of the dye molecules inside the vesicles was monitored by a PTI QuantaMaster spectrofluorometer (Photon Technology International Inc.). Emission intensity ratio measurements at two wavelengths 640 nm/615 nm, using 560 nm excitation were used to track the membrane potential changes. The dye response to $\Delta\psi$ was calibrated by imposing defined $K^+$ gradients on the membrane where the $K^+$ permeability was selectively increased by the ionophore valinomycin (10 nM) thereby establishing different $K^+$ diffusion membrane potentials. The created $\Delta\psi$ was calculated from the Nernst equation, $\Delta\psi=(RT/F)\ln([K^+]out/[K^+]in)$. The membrane potential developed by the reconstituted ATP synthase was dissipated by 1 µM FCCP. PL-reconstituted $F_1F_o$ activity was blocked by 1 µM oligomycin and 500 µM DCCD.

**Electrophysiological measurements**
Electrophysiological characterization of reconstituted complex V was performed using conventional bilayer lipid membrane techniques. Briefly, "black" lipid membranes were formed across a 200 µm aperture that separated the *cis*- and *trans*-compartments of a custom designed and fabricated chamber. The lipid ratio of phosphatidylcholine (PC) : phosphatidylethanolamine (PE) : cardiolipin (CL) was 4.5:4.5:1 by mass. Proteoliposomes were added to the *trans* chamber (simulating the cytosolic side) which was held at virtual ground and contained a solution composed of (in mM): 150 KCl, 20 Tris-HCl, 1 EGTA, 0.5 ADP, 10 $PO_4^{3-}$ (with 50 µM DTT, pH 7.2). Measurements were made after spontaneous vesicle fusion and protein incorporation into the lipid bilayer membrane had been achieved. Voltage was applied (Warner BC-535 Bilayer Clamp amplifier) to the *cis* chamber (simulating the mitochondrial matrix) with the same composition as the *trans* solution except having 50 mM KCl. Single channel currents were filtered by a low-pass 8-pole Bessel filter at a corner frequency of 2 kHz and digitized at a sampling rate of 50 kHz. All measurements were obtained at room temperature (22-23°C). Current-time integral was calculated with the Clampfit and Origin softwares for steady voltages, and for voltage ramp protocol from -60 mV to +60 mV with duration of 8.5 seconds.

**Combined single photon and electrophysiological measurements**
We sought to further analyze these data by quantifying and comparing the ATP synthesis rates independently derived from single molecule electrical (described in main text) and bioluminescence measurements (described below). To derive a quantitative estimate of ATP production rate we used bioluminescence-emitted photon measurements (as generated by the ATP synthesized from one or a small number of ATP synthase molecules), concatenated with the efficiencies of the ATP-luciferase/luciferin reaction (using the bioluminescence quantum yield for firefly (*Photinus pyralis*) luciferin ($\Phi_{Bl}$=0.48; (Niwa et al., 2010; Wang et al., 2011), the optical collection efficiency of the microscope (Spring, 2003) and camera detection characteristics (using quantum efficiency and calibration-constants supplied by the manufacturer with the EM gain factor used in the experiments).
A bilayer was formed on the glass pipettes at the hydrogel/hydrogel interface based on the protocol described by (Ide et al., 2008). Glass pipettes for bilayer were made from borosilicate capillary tubing with outer and inner diameters of 1.5 and 0.9 mm (Warner Instruments), respectively, using pipet puller (P-97, Sutter Instruments). The diameter of the tip was 30-60 µM, the tip was fire-polished with coating and polishing microforge CPM-2 (ALA Scientific Instruments, NY). Hydrogel 1.5-2% agarose, type IX-A (ultra low cooling temperature) in 50 mM KCl solution was dissolved by heating, cooled down at 21 C, mixed with luciferase/luciferin solution (ATP Bioluminescence Assay Kit HS II, Roche Diagnostics) and then aspirated into the pipette from the tip, and hydrogel was polymerased by cooling in refrigerator protected from the light. Glass pipette was back filled with 50 KCl solution just before experiment and put in Axopatch 200B clamp holder. Heated and dissolved hydrogel of 1.5-2.0% agarose, type VII-A (low cooling temperature) in 150 KCl solution, was put on the bottom of the glass of handmade chamber. A small amount of 2 µL of proteoliposomes suspension containing 1-5 mg/ml of proteins was added to the hydrogel (trans chamber, simulating the mitochondrial matrix) which was held at virtual ground and contained a solution composed of (in mM): 150 KCl, 20 Tris-HCl, 1 EGTA, 0.1 ADP, 10 $PO_4^{3-}$ (pH 7.2). A lipid solution (10 mg/ml in n-decan) of L-α-Phosphatidylcholine and Cardiolipin from bovine heart (Sigma) was layered over the hydrogel with proteoliposomes. Measurements were made after spontaneous vesicle fusion and protein incorporation into the lipid bilayer membrane had been achieved. Voltage was applied (Axopatch 200B patch clamp amplifier) to the glass pipette (cis chamber, simulating the cytosolic side) with the same composition as the trans solution except having 50 mM KCl. Single channel currents were recorded at frequency of 2 kHz and digitized at a sampling rate of 50 kHz. All measurements were obtained at room temperature (22-23°C). Oligomycin (0.1 -1.0 µM) and venturicidin (1.5 µM) solution was injected into the chamber with Nanoliter2000 (WPI) microinjector.
Bioluminescence was detected using an Andor iXon$^{EM}$+ 897D back-illuminated EMCCD camera with 95% quantum efficiency at 560 nm (the wavelength of the equivalent photon from the mean of the integrated luminescence-spectrum energy; see below), attached to a Nikon inverted microscope inside a custom-built light-tight Faraday cage enclosure. The luminescence spectrum of the luciferase-luciferin ATP-detection system was measured with a PTI scanning spectrometer (sealed from external light contamination) in the 500-650 nm range for the EMCCD camera's signal calibration under the same physico-chemical conditions of the present experiments (i.e., same pH and ionic conditions, etc.; initiating the reaction with ATP at a level compatible with detection, and making the measurements at a point of stable light output in a quartz cuvette). The dark current/background noise (mean+1σ)-subtracted EMCCD camera signal (A/D counts) was converted to incident photon energy (eV) using physical- and calibration-constants supplied by the manufacturer together with the EM gain factor used in the experiments, in turn yielding the luminescence-related photon counts**.**
Bioluminescence-generated light was collected using a 40x/1.15 NA water-immersion long-working-distance Nikon objective lens and directed to the camera by an optical-grade front-surface mirror (light path transmission estimated

at 90%). The quantitative assessment of the bioluminescence light collected by the lens was estimated as follows. First, we calculated the amount of light able to enter a lens from an isotropically-emitting point source at its focal point, such as would be the case in the present experiments with the photons produced from the ATP synthesized from one or a small number of ATP synthase molecules being emitted from a diffraction-limited nanoscopic volume. This is straightforwardly derived from the geometry of the optics problem, as an equivalent cone of light (its tip at the emission point source) with a 2D acceptance angle, α, being able to enter the lens (determined by the properties of the lens, the numerical aperture (NA), and the refractive index (n) of the light path). The solid angle (Ω) cut out of the sphere by the cone is a spherical cap, where the complete sphere has a solid angle of 4π steradians. The ratio of this cone's cap to the complete sphere solid angle (Ω/4π) is the fraction of the emitted photons that can be collected by the lens. The solid angle Ω that results from the plane angle of the cone α is given by $\Omega = 2\pi*(1-\cos(\alpha/2))$. Since $NA = n*\sin(\alpha/2)$, the solid angle of light able to enter the lens is given by $\Omega = 2\pi*(1 - \cos(\sin^{-1}(NA/n)))$. Thus, the maximum fraction of the emitted light collected by the lens used in the present experiments Ω/4π is 0.249.
Propagated uncertainties in the calculated ATP synthesis rates derived from photon measurements were estimated as the root-sum-square of the products of each of the component-variable errors (approximating σ) with their respective partial derivatives. Uncertainties in the calculated ATP synthesis rates derived from electrical measurements were estimated from the background electrical noise of the lipid bilayer.

**$IF_1$ gene silencing by RNA interference**
Neonatal cardiac myocytes were isolated from 1- to 3-day-old Wistar rats by enzymatic digestion (Chesley et al., 2000) and cultured for two days before transfection with a pool of four short interfering RNAs (siRNAs; 100 nM) targeted specifically to $IF_1$ using GeneSilencer reagent (Gene Therapy Systems Inc., San Diego, California, USA) according to the protocol provided by the company. Four siRNA duplexes were designed and synthesized by Dharmacon Inc. (Lafayette, Colorado, USA) to target the rat $IF_1$: (a) 5'-AAACAGATCGAACGGCATA-3'; (b) 5'-AAAGAATAGTGAGCATTGA-3'; (c) 5 '-GGAGATAGAGCGTCTGCAA-3'; (d) 5'-CGTATGAGGGTCCTGCAAA-3'. As a negative control, cells were transfected with siRNA against GFP (Dharmacon Inc.). Experiments were performed 72 hours after transfection**.**

**Immunofluorescence**
Cultured neonatal cardiac myocytes were fixed with 2% formaldehyde in PBS (phosphate buffered saline; Sigma) followed by permeabilization with 0.5% Triton X-100/PBS. We blocked nonspecific cross reactivity by incubating the samples for 4 h in 1% BSA/PBS (Jackson ImmunoResearch, West Grove, PA, USA) and then performed cell immunolabeling via incubation with primary antibodies diluted in 1% BSA/PBS overnight, washing in PBS followed by appropriate fluorescence label-conjugated secondary antibodies (Jackson ImmunoResearch). Labeled cells were imaged by a Zeiss LSM-510 (Carl Zeiss, Jena, Germany) confocal microscope using a 63x/1.4 N.A. oil immersion lens.

**Cell isolation.**
Adult cardiac myocytes were isolated from Sprague-Dawley rats (2–4 months old), and WT or $IF_1$ KO mice by using standard enzymatic techniques, as described previously (Capogrossi et al., 1986). Isolated cardiomyocytes were suspended in a solution containing (in mM): NaCl 137, KCl 4.9, $MgSO_4$ 1.2, $NaH_2PO_4$ 1.2, glucose 15, HEPES 20, and $CaCl_2$ 1.0 (pH 7.4). Handling of animals was conducted in accordance with NIH guidelines for animal care and use.

**Determination of mitochondrial permeability transition threshold**
We have previously developed and extensively tested a model enabling the precise determination of the mPTP sensitivity to oxidant stress in intact cardiac myocytes (Juhaszova et al., 2004; Zorov et al., 2000). Briefly, small numbers of mitochondria inside isolated cardiomyocytes were exposed in situ to conditions of oxidative stress by repetitive (2 Hz) laser line-scanning (with imaging) of a single row of mitochondria in a cell loaded with 100 nM tetramethylrhodamine methyl ester (TMRM), using a Zeiss LSM-510 inverted confocal microscope. This results in incremental, additive exposure of only the laser-exposed area to the photodynamic production of ROS and consequent mPTP induction. The ROS-threshold for mPTP induction ($\boldsymbol{t}_{\mathbf{MPT}}$) was measured as the average time necessary to induce the mPTP due to the local buildup of ROS in a row consisting of ~25 mitochondria.

**Measurements of the mitochondrial volume in intact cardiomyocytes**
Transmitted optics line-scan imaging to assess changes in mitochondrial volume in individual cardiomyocyte due to exposure to 30 μM DZ, 10 nM DADLE, 10 μM HOE with or without the $F_o$-inhibitor VENT. Changes in

mitochondrial volume were assessed using a Zeiss 63x/1.4 N.A. oil immersion lens, 633 nm laser illumination, scanning 14.1 pixels/µm along the cell long axis for 72.6 or 145.1 µm). Fourier analysis (ImageJ, W. Rasband, NIH, Bethesda, Maryland, USA) of repeating intensity of the linescan image provided the long-axis spacing of the sarcomere and mitochondrial compartments (from the 1st and 2nd order spectral peaks, enabling resolution of changes in dimension of ~1% in 1 µm structures). The position of the major Fourier spectral peak corresponding to ~1 µm spatial structures gives the average mitochondrial diameter, and the peak-width gives an index of the variability about the mean (details of this procedure are described in (Juhaszova et al., 2004)).

**Flavoprotein fluorescence and $VO_2$ measurements**

Oxidation of the flavoprotein (FP) pool is known to increase the blue light-excited cellular autofluorescence (Juhaszova et al., 2004; Sato et al., 1998). Endogenous flavoprotein fluorescence was excited by an argon 488 nm laser line (attenuated to 1%) on a Zeiss LSM 510 using a 40x 1.3 NA oil objective. Emitted fluorescence was recorded at LP 505 nm. Relative fluorescence was calibrated to the values measured after 2,4-dinitrophenol (DNP) (fully oxidized) and cyanide exposure (fully reduced). Cells were continuously perfused (1-2 ml/min) and a time series of images was collected at intervals of 20 seconds. After baseline fluorescence was acquired (5 min), cells were exposed to diazoxide (10 min) and the experiment was completed by exposing the cells to DNP. Fluorescence traces from individual cell in the acquisition view were analyzed by MetaMorph image analysis software (Molecular Devices, Sunnyvale, CA) and then averaged. Results are normalized to the dynamic range achieved in each cell when exposed to DNP, which by producing sufficient respiratory uncoupling, yields the maximum degree of autofluorescence change possible due to FP oxidation.

Cell respiration ($VO_2$) was measured by the Clark-type $O_2$ electrode based Mitocell S200 micro respirometry system (Strathkelvin Instruments Ltd. North Lanarkshire, UK).

**Mitochondrial swelling, membrane potential and $K^+$ flux determination in isolated mitochondria**

Mitochondria from guinea pig heart were isolated and assayed as described elsewhere (Aon et al., 2010). Briefly, mitochondrial swelling, membrane potential (ΔΨm), and PBFI were monitored simultaneously with a spectrofluorimeter (Photon Technology, Inc.) in an experimental solution containing (in mM): 250 sucrose (or 137 KCl), 0.5 EGTA, 2.5 $MgCl_2$, 20 HEPES, 2 Pi, pH 7.2. Mitochondrial swelling was measured as a decrease in the 90° light scattering (LS) signal using 520 nm excitation. ΔΨm was recorded using 100 nM TMRM and quantified with a ratiometric method (Scaduto and Grotyohann, 1999). PBFI fluorescence was monitored ratiometrically (340/380 nm excitation at 495 nm emission).

**High throughput assay of mitochondrial respiration, OCR**

High-throughput–automated 96-well microplate reader analysis of respiration (XF96 extracellular flux analyzer; Seahorse Bioscience) (Aon et al., 2012; Cortassa et al., 2017a) was performed with freshly isolated mitochondria from rat hearts. The $O_2$ consumption rate (OCR) was evaluated in the linear range of detection, using the equivalent of 5µg of mitochondrial protein with glutamate and malate (G/M), 5mM each, in the following assay media, 200µl final volume, pH 7.2 at 37°C: *(i) KCl-based*, containing (in mM): 137 KCl, 2 $KH_2PO_4$, 0.5 $Na^+$-EGTA, 2.5 $MgCl_2$, and 20 $Na^+$-HEPES at pH 7.2, in presence of 0.2% fatty acid–free BSA (Aon et al., 2012); or *(ii) sucrose-based* containing (in mM): 250 sucrose (substituting KCl), 2 $NaH_2PO_4$ (substituting $KH_2PO_4$), 0.5 $Na^+$-EGTA, 2.5 $MgCl_2$, and 20 $Na^+$-HEPES. Both assay media were at the same osmolality (260 mOsm) as measured with Vapro, a vapor pressure-based osmometer.

The OCR corresponding to states 3 and 4 respiration was determined upon addition of 1mM ADP followed by 10µM Oligomycin, respectively. Respiratory Control Ratio (RCR=state3/state4) of 4 to 5 were obtained. The OCR is expressed in ng-at O $min^{-1}$ $mg^{-1}$ mitochondrial protein to allow comparison with reported data.

The day before the experiment, 120µl polyethylenimine (1:15000 dilution in buffer B of a 50% solution of polyethylenimine) were added to the wells of the XF96 plate, and incubated overnight at 37°C. Before the experiment, the solution of polyethylenimine was removed. After transfer of appropriate amounts of mitochondrial suspension into each well (the equivalent of 5µg of mitochondrial protein), the microplate was centrifuged at 2,000 x *g* for 14 min at 4°C using a swinging bucket rotor (Sorvall RT 6000D). To avoid temperature inhomogeneity effects, the plate was incubated at 37°C for 20 min before starting the assay in the Seahorse Bioscience equipment.

**Determination of ΔΨm, ΔpH and mitochondrial volume**

We used radioactive tracer methods to measure the proton motive force (PMF), and its components, membrane potential (ΔΨm) and ΔpH (Rottenberg, 1979, 1989). These measurements were accompanied by mitochondrial volume determinations using $^3H$-labeled water ($^3H_2O$) and $^{14}C$-labeled mannitol ($^{14}C$ mannitol) in states 4 and 3 respiration, which are needed to quantify $\Delta\Psi m$ and $\Delta pH$. The mitochondrial volume in $K^+$- or sucrose-based medium (see composition above) was performed in triplicate in each of three independent mitochondrial preparations, under the same conditions in which OCR was measured. For the timing, we used as a reference our fluorometric experiments in which the kinetics of volume (90º light scattering) and $\Delta\Psi m$ (TMRM ratiometric method) changes were measured in energized mitochondria with G/M 5/5 mM (state 4) followed by addition of different ADP concentrations (state 3) (Supplemental Fig S4 G-M). Mitochondrial volume, $\Delta\Psi m$ and $\Delta pH$ were determined in parallel for state 4 (5 min with G/M 5 mM each) or state 3 mitochondria (5 min with G/M and 0.5mM ADP) (Rottenberg, 1979, 1989). The radioactive tracers: 1μCi of $^3H_2O$ and 0.2 μCi of $^{14}C$ Mannitol for volume, or 1 μCi of $^3H$ Tetraphenylphosphonium bromide [phenyl-$^3H$] [$^3H$ TPP] for $\Delta\Psi m$ or 0.2 μCi of $^{14}C$ Dimethyl oxazolidine-2,4 dione-5,5 [2-$^{14}C$] [$^{14}C$ DMO] for $\Delta pH$, were added immediately before the substrates, to 2 ml Eppendorf microcentrifuge tubes containing 0.25 ml of assay $K^+$- or sucrose-based medium with 1.4 - 1.8 mg of mitochondrial protein/ml, and incubated the indicated times. After incubation, the tubes were centrifuged at 13,000 x g for 3 min. The whole volume of the supernatant was recovered and counted, and the pellet was also counted after being dissolved with 0.1 ml of perchloric acid 20% (wt/wt), using a scintillation counter (Beckman LS 6500, Beckman Coulter liquid scintillation analyzer., Fullerton CA).

The calculations for determining mitochondrial volume, $\Delta\Psi m$ and $\Delta pH$ were performed according to (Rottenberg, 1979, 1989).

Several controls were performed in order to account for *i)* count spillover: mitochondria were incubated with one radioactive tracer at a time, centrifuged, and supernatant and pellet counted in the $^{14}C$ and $^3H$ channels in duplicate; *ii)* Nonspecific binding of radioactive tracers: mitochondria were subjected to osmotic lysis, and the same centrifugation as well as counting protocol was followed; and *iii)* quenching of the $^{14}C$ and $^3H$ tracers introduced by perchloric acid: 0.01 μCi of each radioactive tracer was dissolved in 0.5ml of water; half of the volume was acidified with perchloric acid whereas the other half was counted directly.

**P/O ratio determination by high resolution respirometry**

The P/O ratio was determined in a High-Resolution Respirometry Oxygraph-2k Oroboros instrument, equipped with a Clark type polarographic oxygen sensor evaluating $PO_2$ in a stirred 2 mL chamber. The dissolved $O_2$ was continuously monitored during incubation of the mitochondrial suspension in the same $K^+$- or sucrose-based medium used in Seahorse OCR determinations. The mitochondrial suspension (200 μg mitochondrial protein) was equilibrated in the absence of ADP until a stable rate of $O_2$ consumption ($VO_2$) was attained followed by the addition of 50 μM ADP. $VO_2$ was monitored until ADP was consumed, and the area under the $VO_2$ curve measured to quantify the amount of $O_2$ consumed in response to the pulse of ADP.

**Materials**: [$^3H$]-$H_2O$ and D-[2-$^{14}C$]-Mannitol from NEN Radiochemicals were purchased from PerkinElmer (Walthan, MA) whereas [phenyl $^3H$] TPP and [2-$^{14}C$ DMO] were obtained from American Radiolabeled Chemicals, Inc (St Louis, MO).

**Protein binding ($K_d$) measurements**

For dissociation constant ($K_d$) measurements of $IF_1$ to Bcl-2-family proteins (Bcl-2, Bcl-xL), 100 nM of Bcl-protein was admixed with 0-200 nM of $IF_1$. Binding was monitored via changes in intrinsic Trp fluorescence, excited at 295 nm and fluorescence detected at 350 nm. Measurements were performed using a PTI (Photon Technology, Inc.) fluorimeter. Extracted intensities were fitted to a generalized binding equation:

$$F = F_0 + A \cdot \frac{([I]+[B_t]+K_d) - \sqrt{([I]+[B_t]+K_d)^2 - 4[I][B_t]}}{2[B_t]}$$

where $[B_t]$ is the total Bcl-protein concentration, $[I]$ is the concentration of $IF_1$, $F_0$ and $A$ are background fluorescence and a scaling factor, respectively.

**Inhibition of GSK-3β activity via Ser-9 phosphorylation**

We identified GSK-3β activity as a pivotal kinase a diverse range of signaling pathways to cause functional protection of the mitochondrial permeability transition pore (mPTP) against induction by oxidant stress (Juhaszova et al., 2004). The activity of GSK-3β is inversely related to the phosphorylation status of serine (Ser)-9. Dephosphorylation of this site, or mutations that prevent phosphorylation, result in activation of the kinase (Antos et al., 2002). We found that constitutive activation of GSK-3β prevents the ability to engage protective signaling in cardiomyocytes through pathways including PKA, PKB, PKC, PKG and p70s6K. We also found that Dz (as well as numerous other distinct protection-signaling triggers (Juhaszova et al., 2004)), resulted in GSK-3β phosphorylation on regulatory serine-9 (including the fraction isolated from mitochondrial membranes). Thus, in the present experiments we compared the ability of Dz to phosphorylate GSK-3β on regulatory Ser-9 in cardiomyocytes with $IF_1$-expression knocked down by ~75% through siRNA, compared to controls (Fig 4M). While Dz causes a significant increase in P-GSK-3β in control cells, this was largely prevented in $IF_1$-siRNA treated cells. We conclude that $mK_{ATP}$-related protection signaling via GSK-3β requires the functional presence of $IF_1$, thus implicating the role of the $F_1F_o$. Further support for this role comes from the ability of Vent to similarly block phosphorylation of GSK-3β by Dz (data not shown).

**Generation of IF1$^{-/-}$ mice**

Mice carrying a genetically inactivated allele at the Atpif1 locus (C57BL/6NTac-Atpif1$^{tm1a(EUCOMM)Wtsi}$/WtsiCnbc), originally generated by the Wellcome Trust Sanger Institute for the EUCOMM project, within the International KnockOut Mouse Consortium (IKMC) (Bradley et al., 2012), were obtained from the Spanish node of the European Mouse Mutant Archive (EMMA, http://www.infrafrontier.eu, mouse strain EM:05233) at the National Centre for Biotechnology in Madrid (Spain) and bred to homozygosity. Resulting homozygous mice are referred to as IF1$^{-/-}$ in this manuscript. Mice were genotyped through a combination of separate PCR reactions that detect LacZ, the gene-specific wild type allele, and a mutant allele-specific short range PCR.

PCRs primer pairs and expected size bands:

| PCR type | Forward primer | Reverse primer | Expected size (bp) |
|---|---|---|---|
| Mutant PCR | Atpif1_46360_F | CAS_R1_Term x | 114 |
| Wild type PCR | Atpif1_46360_F | Atpif1_46360_R | 396 |
| LacZ PCR | LacZ_2_small_F | LacZ_2_small_R | 108 |

Primer sequences:

| Primer names: | Primer sequence (5' > 3') |
|---|---|
| CAS_R1_Term | TCGTGGTATCGTTATGCGCC |
| Atpif1_46360_F | TGCCTGACATTGGTATTGGG |
| Atpif1_46360_R | GTGCAGCTTGTGGGAGTCAG |
| LacZ_2_small_F | ATCACGACGCGCTGTATC |
| LacZ_2_small_R | ACATCGGGCAAATAATATCG |

**Phylogenetic tree of the BH3 extended peptides**

The $IF_1$ peptide BH3 motif appears to have diverged from the evolutionary branch originating BID and BIK Bcl-2 family members. The analysis of a 35 residue fragment from $IF_1$ ranging from fungi to human aligned together with 79 fragments containing BH3 motifs with similarities to the 26 residue BAD peptide used in our functional studies (Fig 5B,C) leads to the tree shown in Fig S7A. MEGA enabled the calculation of the divergence time of $1690\times10^6$ years when $IF_1$ ancestor proteins separated from the ones that would originate BID, BIK and Bcl-2-like protein 13. The tree obtained from the Clustal Omega alignment shows largely the same group division that has been considered elsewhere (Aouacheria et al., 2013). In spite of our analysis being limited to only a fragment containing the BH3 motif, the same groups were obtained, i.e., BID-like, Bcl-2-like, also containing Bcl-xL (Lanave et al., 2004), and a Bax-like (Aouacheria et al., 2013). Additionally, an independent assessment of the alignment and phylogenetic analysis was performed with the program BAli-Phy. This rendered essentially a similar tree which only differed from the tree in Fig S7A regarding the relative position of the Beclin and BAX groups (Fig S7B). In our case there was no drift in the evolution of the hidden Markov Model (HMM) and the analysis of several Markov Chain Monte Carlo (MCMC) runs demonstrated convergence of the various chains toward a consensus alignment and phylogenetic tree (convergence criteria for PSRF-80%CI <= 1.001 and ASDSF[min=0.010] = 0.008). The main difference with the HMM described by Aouacheria (Aouacheria et al., 2013) is that longer fragments containing the so-called affinity enhancing motifs flanking the BH3 motif (Yang, 2010) were considered in the present work which renders the alignment more stringent.

**Phylogenetic tree of the BH3 extended peptides**

*Sequence Alignment and tree computation methods*

The sequences aligned were obtained from the UniProt website (EMBL-EBI; http://www.uniprot.org). First, the BH3-containing proteins were obtained by performing a BLAST search of the BAD peptide (26 AA peptide used in our functional studies). A collection of 79 sequences of BH3 containing proteins was retrieved expanding from yeast to mammals including proteins from the BAD, BAK, BAX, BID, BIK, PUMA, MCL-1 and BECLIN groups. Then, a collection of 28 IF1 sequences, mostly manually annotated and reviewed, was gathered and aligned using Clustal Omega (http://www.clustal.org/omega). The fragments of the sequences obtained by this alignment were saved and used for subsequent analysis. These fragments correspond to 35 amino acids that contain the pattern of the original BAD peptide that was used as the seed. The unaligned peptides were used as input sequences for BAli-Phy (v 2.3.6 (Redelings and Suchard, 2005)) a Linux program that uses Bayesian estimation and Markov Chain Monte Carlo (MCMC) methods to sample from the posterior distribution of alignments. Both alignments obtained from Clustal Omega and from BAli-Phy were utilized to construct the respective phylogenetic trees using the program MEGA 6.0 (Tamura et al., 2013) which allows computation of the the distances and the evolutionary times after linearization of the distances of divergence. For those calculations the reference point for the divergence between man and an ascomycete was $1540\times10^6 \pm 250\times10^6$ years (1540 Myr). The calculation of the tree was performed with the Neighbor Joining method (Saitou and Nei, 1987).
